# Dose-dependent disruption of hepatic zonation by 2,3,7,8-tetrachlorodibenzo-*p*-dioxin in mice: integration of single-nuclei RNA sequencing and spatial transcriptomics

**DOI:** 10.1101/2022.06.15.496321

**Authors:** R. Nault, S. Saha, S. Bhattacharya, S. Sinha, T. Maiti, Tim Zacharewski

**Affiliations:** Department of Biochemistry & Molecular Biology, Michigan State University, East Lansing, MI, USA 48824; Institute for Integrative Toxicology, Michigan State University, East Lansing, MI, USA 48824; Department of Statistics and Probability, Michigan State University, East Lansing, MI, USA 48824; Biomedical Engineering Department, Pharmacology & Toxicology, Institute for Quantitative Health Science and Engineering, Michigan State University, East Lansing, MI, USA 48824; Department of Statistics, Texas A&M University, College Station, TX, USA 77840

**Keywords:** Liver, Metabolic zonation, Nonalcoholic fatty liver disease, Toxicant associated fatty liver disease, Toxicology

## Abstract

2,3,7,8-Tetrachlorodibenzo-*p*-dioxin (TCDD) dose-dependently induces the development of hepatic fat accumulation and inflammation with fibrosis in mice initially in the portal region. Conversely, differential gene and protein expression is first detected in the central region. To further investigate cell-specific and spatially resolved dose-dependent changes in gene expression elicited by TCDD, single-nuclei RNA sequencing and spatial transcriptomics were used for livers of male mice gavaged with TCDD every 4 days for 28 days. The proportion of 11 cell (sub)types across 131,613 nuclei dose-dependently changed with 68% of all portal and central hepatocyte nuclei in control mice being overtaken by macrophages following TCDD treatment. We identified 368 (portal fibroblasts) to 1,339 (macrophages) differentially expressed genes. Spatial analyses revealed initial loss of portal identity that eventually spanned the entire liver lobule with increasing dose. Induction of R-spondin 3 (*Rspo3*) and pericentral *Apc*, suggested dysregulation of the Wnt/β-catenin signaling cascade in zonally resolved steatosis. Collectively, the integrated results suggest disruption of zonation contributes to the pattern of TCDD-elicited NAFLD pathologies.

**SYNOPSIS:** 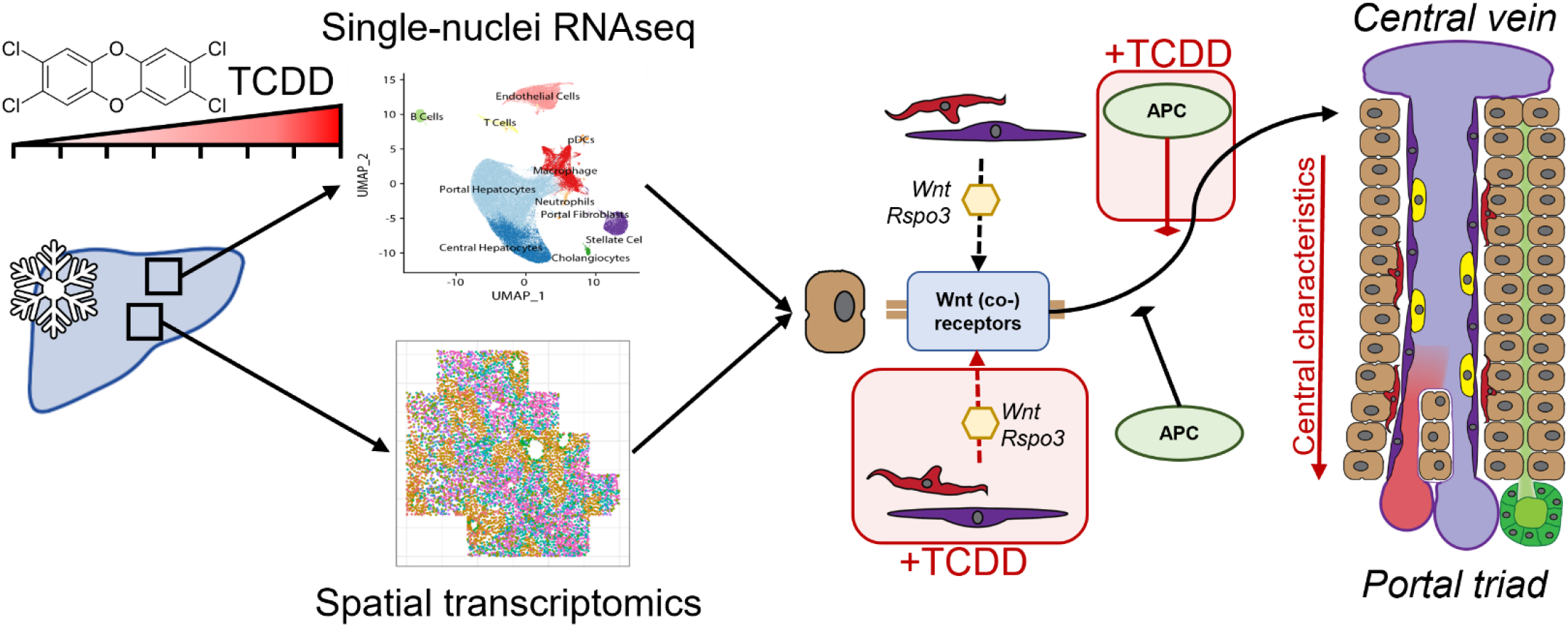

Single-nuclei RNA sequencing (snRNAseq) and spatial transcriptomics were integrated to investigate cell-specific and spatially resolved dose-dependent changes elicited by TCDD. We show that TCDD causes a loss of zonal characteristics that disrupts spatially defined metabolic functions.

- Dose-dependent analyses show higher responsiveness of central hepatocytes despite hepatotoxicity occurring initially in the portal region.
- Integration of snRNAseq and spatial transcriptomics demonstrates a loss of hepatocytes with portal characteristics.
- TCDD disrupted spatially resolved expression of β-catenin signaling members that are critical in maintaining liver zonation.
- Spatial transcriptomics and snRNAseq shows induction of R-spondin3 from nonparenchymal cells which serve as cue for the β-catenin pathway.

## INTRODUCTION

The spatial organization of the different liver cell types is essential for normal function. The adult liver consists of at least 11 predominant cell types that includes hepatocytes, cholangiocytes, hepatic stellate cells (HSCs), resident macrophages (aka Kupffer cells), and liver sinusoidal endothelial cells (LSECs) (Halpern *et al.*, 2018; Halpern *et al.*, 2017; Nault *et al.*, 2021; Xiong *et al.*, 2019). Other single-cell studies have also identified additional cell (sub)types such as spatially resolved LSECs and HSCs, as well as immune cell subtypes with distinct gene expression characteristics (Bleriot and Ginhoux, 2019; Dobie *et al.*, 2019; Halpern *et al.*, 2018). Oxygen and nutrient gradients that run from the portal triad to the central vein further influence the cellular and spatial organization of liver lobules. Nutrient rich blood from the portal vein, mixed with oxygenated blood from the hepatic artery, impart zone specific expression of enzymes for β-oxidation and gluconeogenesis in the portal region (zone 1) while lipogenesis, glycolysis, and phase I xenobiotic metabolic activities are focused in the central region (zone 3) (Cunningham and Porat-Shliom, 2021; Jungermann, 1995). This spatial organization allows for interdependent metabolic pathways to co-localize and optimize metabolism while avoiding interference and energy waste from opposing pathways (Gebhardt and Hovhannisyan, 2010). Disruption of the functional relationship within this cellular heterogeneity has been associated with several adverse health consequences (Cunningham and Porat-Shliom, 2021; Hall *et al.*, 2017; Panday *et al.*, 2022; Soto-Gutierrez *et al.*, 2017).

In addition to genetics, lifestyle, and diet, accumulating evidence suggests exposure to structurally diverse chemicals and environmental contaminants promotes the development of metabolic disorders such as NAFLD, type II diabetes, cardiovascular disease, and hepatocellular carcinoma (Cave *et al.*, 2010; Lee *et al.*, 2006; Taylor *et al.*, 2013). For example, the persistent organic pollutant and potent agonist of the aryl hydrocarbon receptor (AHR) 2,3,7,8-tetrachlorodibenzo-*p*-dioxin (TCDD) induces hepatic lipid accumulation (steatosis) that progresses to steatohepatitis with fibrosis (Boverhof *et al.*, 2005; Fader *et al.*, 2017b; Nault *et al.*, 2015). In humans, TCDD and related compounds are associated with dyslipidemia and inflammation (Pelclova *et al.*, 2006; Taylor *et al.*, 2013; Wahlang *et al.*, 2019; Warner *et al.*, 2013). In mice, AHR activation by TCDD elicits cell-specific and spatially resolved histological and gene expression responses. Specifically, at lower doses of TCDD CYP1A1 induction occurs in the central region while lipid accumulation and inflammation first appears in the portal region (Andersen *et al.*, 1997; Boverhof *et al.*, 2005; Fader *et al.*, 2017b). A recent single-nuclei RNA sequencing (snRNAseq) study reported cell-specific differential gene expression, with a putative loss of portal hepatocytes (Nault *et al.*, 2021). Additional studies are needed to determine if this hepatocyte specific zonal loss is truly a decrease in cell number or repression of characteristic portal hepatocyte marker expression.

A complex network of cues, receptors, and signaling cascades interact within and between cell subtypes to establish and maintain hepatic zonation. This includes the Wnt/β-catenin, RAS/ERK, Hippo, hedgehog, glucagon, HNF4α, and Dicer signaling pathways (Burke *et al.*, 2009; Cunningham and Porat-Shliom, 2021; Gebhardt and Hovhannisyan, 2010; Kietzmann, 2019). The Wnt/β-catenin signaling pathway is a central driver of liver zonation mediated by hepatic, angiocrine, and extrahepatic secreted factors and their cognate receptors (Burke *et al.*, 2009; Gebhardt and Hovhannisyan, 2010; Rocha *et al.*, 2015; Zhang *et al.*, 2020). In the canonical pathway, secreted Wnt ligands bind to Frizzled (FZD) and low-density lipoprotein receptor-related proteins (LRP) preventing the degradation of β-catenin by a destruction complex comprising adenomatous polyposis coli (APC) and glycogen synthase kinase (GSK3β). R-spondins and leucine-rich repeat-containing G protein-coupled (LGR) receptors further potentiate Wnt/β-catenin signaling (Park *et al.*, 2020; Rocha *et al.*, 2015). The Wnt/β-catenin pathway is also shown to interact with the AHR that is highly expressed in the central hepatic region (Braeuning *et al.*, 2011; Gerbal-Chaloin *et al.*, 2014; Nault *et al.*, 2021; Prochazkova *et al.*, 2011; Vondracek and Machala, 2016; Yang *et al.*, 2022). Developmental AHR deletion results in nuclear β-catenin localization and induction of target genes *Axin2, Ccnd1, Myc*, and *Lef1* as well as the modulation of basal *Cyp1a1* expression (Moreno-Marin *et al.*, 2018). Furthermore, β-catenin enhanced the induction of ligand activated AHR target genes. β-Catenin also formed a complex with the AHR further suggesting interactions between the Wnt/β-catenin and AHR signaling pathways (Moreno-Marin *et al.*, 2018).

Our previous work has suggested a global loss of hepatic function following TCDD treatment, especially functions associated with hepatocyte metabolism (Nault *et al.*, 2021; Nault *et al.*, 2017a). Moreover, it is established that AHR activation by TCDD perturbs mechanisms associated with the maintenance of zonation including HNF4α binding and Wnt signaling (Cholico *et al.*, 2022; Moreno-Marin *et al.*, 2018; Yang *et al.*, 2022). In this study we aimed to further characterize the cell-cell interactions in the mouse liver that were disrupted by TCDD and resulted in the loss of hepatic zonation. We hypothesize that the loss of liver function and portal hepatocytes following treatment with TCDD involves the disruption of spatially resolved Wnt/ β-catenin signaling. To accomplish this, dose-dependent snRNAseq and spatial transcriptomic datasets were integrated to investigate zone-specific effects at lower doses that progress to panacinar steatosis at higher doses.

## MATERIALS AND METHODS

### Animals and treatment

Male C57BL/6 mice were received from Charles Rivers Laboratories (Portage, MI) at postnatal day (PND) 25 and acclimated until PND 28. Mice were randomly assigned to Innocages (Innovive, San Diego, CA) with ALPHA-dri bedding (Shepherd Specialty Papers, Chicago, IL) at 30-40% humidity and a 12-hour light/dark cycle (3 mice/cage and dose group). Mice had free access to Harlan Teklad 22/5 Rodent Diet 8940 (Envigo, Indianapolis, IN) and Aquavive water (Innovive). On PND 28 and every 4 days thereafter, for a total of 7 administrations (24 days), mice were orally gavaged with sesame oil vehicle (Sigma-Aldrich, St. Louis, MO), 0.01, 0.03, 0.1, 0.3, 1, 3, 10, or 30 μg/kg TCDD (AccuStandard, New Haven, CT) between ZT00 and ZT01. The dose range induced a range of reported histopathologies and transcriptomic responses. On PND 56, 28 days after the initial gavage, mice were euthanized by CO2 asphyxiation and livers were immediately collected, snap frozen in liquid nitrogen and stored at −80°C. All procedures were approved by the Michigan State University Institutional Animal Care and Use Committee and this report meets the ARRIVE guidelines (Percie du Sert *et al.*, 2020). Deposited metadata fulfilled the Minimum Information about Animal Toxicology Experiments (MIATE) reporting guidelines (https://fairsharing.org/FAIRsharing.wYScsE).

### Single-nuclei RNA sequencing

Nuclei were isolated from frozen samples of the right lobe (~200mg) as previously described (https://doi.org/10.17504/protocols.io.3fkgjkw). Two to three biological replicates passing QC were analyzed for each dose group with at least 2 replicates providing sufficient power to identify cell types and assess cell-specific differential expression (Datlinger *et al.*, 2021; McGinnis *et al.*, 2019; Nault *et al.*, 2021). Briefly, livers were diced in EZ Lysis Buffer (Sigma-Aldrich), homogenized using a disposable Dounce homogenizer, and incubated on ice for 5 minutes. The homogenate was filtered using a 70-μm cell strainer, transferred to microcentrifuge tube, and centrifuged at 500 g and 4°C for 5 minutes. The supernatant was removed, and fresh EZ lysis buffer was added for an additional 5 minutes on ice following by centrifugation at 500 g and 4°C for 5 minutes. The nuclei pellet was washed twice in nuclei wash and resuspend buffer (1× phosphate-buffered saline, 1% bovine serum albumin, 0.2-U/μL RNAse inhibitor) with 5-minute incubations on ice. Following the washes, the nuclei pellet was resuspended in nuclei wash and resuspended in buffer containing DAPI (10 μg/mL). The resuspended nuclei were filtered with 40-μm strainer and immediately underwent fluorescence-activated cell sorting using a BD FACSAria IIu (BD Biosciences, San Jose, CA) with 70-μm nozzle at the MSU Pharmacology and Toxicology Flow Cytometry Core (drugdiscovery.msu.edu/facilities/flow-cytometry-core).

Libraries from sorted nuclei were prepared using the 10x Genomics Chromium Single Cell 3’ v3 kit and submitted for 150-bp paired-end sequencing at a depth ≥ 50,000 reads/cell using the HiSeq 4000 at Novogene (Beijing, China). Raw sequencing data were deposited in the Gene Expression Omnibus (GEO; GSE148339). Following the assessment of sequencing quality, CellRanger v3.0.2 (10x Genomics) was used to align reads to a custom reference genome (mouse mm10 release 93 genome build) which included introns and exons to consider pre-mRNA and mature mRNA present in the nuclei. Raw counts were further analyzed using Seurat v4.0.5. Each sample was filtered for (1) genes expressed in at least 3 nuclei, (2) nuclei that express at least 100 genes, and (3) ≤1% mitochondrial genes. Additional quality control assessments was performed using the scater package (v1.18.6). The DoubletFinder v2.0.3 package excluded putative doublets from subsequent analyses. Raw and processed data has been deposited in GEO with the accession ID GSE184506 and the Broad Single Cell Portal (SCP1871).

### Clustering, annotation, and analysis of snRNAseq data

Integration, clustering, and annotation was performed using Seurat. Clustering at varying resolutions (0.05, 0.1, 0.15, 0.2, and 0.3) was assessed to determine changes in cell type populations (**Figure S1**). At 0.1 resolution 11 distinct clusters were identified comparable to previous characterizations of liver cell populations (Halpern *et al.*, 2018; Halpern *et al.*, 2017; Nault *et al.*, 2021; Xiong *et al.*, 2019). For cell annotation, a semi-automated strategy (Nault *et al.*, 2021) was used with published data (GSE148339) as reference. Annotations were manually verified using highly expressed marker genes. Initially unidentified clusters were further examined for marker genes using both literature and data repositories (*e.g.*, panglaoDB and Broad Single Cell Portal).

Differential expression analysis was performed using a single cell Bayesian Test (scBT), a fit-for-purpose test method that effectively controls the false positive rate when testing multiple dose groups (Nault *et al.*, 2022). Genes were considered differentially expressed when the gene 1) was detected in at least 5% of nuclei in at least 1 dose group, 2) had an absolute fold-change ≥ 1.5, and 3) had an adjusted p-value ≤ 0.05. Gene set enrichment analysis was performed using bc3net v1.0.4 in R (4.0.3) with KEGG and MPO gene sets obtained from the Gene Set Knowledgebase (GSKB; http://ge-lab.org/gskb/). Gene sets were collapsed when ≥ 60% of the genes were common and manually annotated based on original annotations.

Cell-cell interactions were determined using CellPhoneDB (Efremova *et al.*, 2020). Cell-cell interactions were determined for each dose independently using 10 iterations and a ligand/receptor percent expression threshold of 25%. For each dose group interactions, the mean value of the ligand/receptor pair was used to calculate fold-change relative to vehicle control. Only interactions with a ≥ 2-fold increase following treatment and a p-value ≤ 0.05 were considered as new treatment induced cell-cell interactions. Raw interaction data for each dose group is available in **File S1**.

### Hepatocyte pseudospatial analysis

Pseudospatial analysis of hepatocyte snRNAseq data was performed as previously described (Nault *et al.*, 2021). In short, nuclei annotated as hepatocytes were extracted from the complete dose-response dataset and reintegrated using Seurat based on the expression of previously characterized spatially resolved genes (Halpern *et al.*, 2017). Following dimensionality reduction using Seurat, Slingshot was used for trajectory analysis along the spatial continuum (Street *et al.*, 2018). Next, we fit negative binomial generalized additive models (NB-GAM) to individual genes using tradeSeq v1.7.4 to evaluate treatment related effects that were considered significant (p ≤ 0.05). Only a small subset of genes were examined by NB-GAM and therefore the p-values were not adjusted. Center of expression (CoE), an indicator of zonal bias, was calculated for each dose and gene with a log normalized expression ≥ 0.5 and expressed in ≥ 50% of nuclei as previously described (Halpern *et al.*, 2017). The maximum CoE travel distance was calculated as the differences between the minimum and maximum CoE values to reflect a dose-dependent change in zonation bias.

### Spatial transcriptomics

Spatial transcriptomics was performed using the Resolve BioScience Molecular Cartography™ system, a probe-based platform with subcellular resolution for detecting gene expression in a tissue section. Two biological replicates for each dose group (0, 0.3, 3, and 30 μg/kg TCDD) were examined by placing 10 μm thick frozen liver sections in O.C.T compound (Sakura, Torrance, CA) in one of 8 designated custom slide regions by the MSU Investigative Histopathology Laboratory (https://sites.google.com/msu.edu/ihpl/home). Frozen slides were shipped to Resolve BioSciences for analysis following manufacturer’s instructions (protocol version 3.0) (D’Gama *et al.*, 2021; Groiss *et al.*, 2021; Guilliams *et al.*, 2022). In short, sections were thawed and fixed with 4% v/v formaldehyde. A total of 99 genes were probed (**Table S1**) overnight, fluorescently tagged, then manually selected regions of interest (ROI) were imaged. Probes were designed using a proprietary algorithm (Resolve Biosciences) for Ensembl gene targets outlined in **Table S1** that were verified for off-target hybridization. Imaging, segmentation, preprocessing, and decoding was performed as previously described (D’Gama *et al.*, 2021; Groiss *et al.*, 2021; Guilliams *et al.*, 2022). Raw and SpatialExperiment formatted data (Righelli *et al.*, 2022) was deposited in GEO (GSENNNN) as well as the Broad Single Cell Portal (SCP1875).

#### Visualization and analysis of spatial transcriptomic images

Visualization of spatial expression was performed using custom code developed by Resolve BioSciences for ImageJ and Python (https://github.com/ViriatoII/polylux_python). Baysor (v0.5.0) was used for cell segmentation (Petukhov *et al.*, 2021) that considered both transcript expression localization as well as DAPI images with a scale of 50 and standard deviation of 12.5%. MERINGUE was used to evaluate co-localization of gene expression (Miller *et al.*, 2021). Analysis was performed independently on each tissue section using the spatially aware strategy with a filtering distance of 150. Spatial autocorrelation and spatial cross correlation were determined for all genes after which groups of spatially co-expressed genes underwent hierarchical clustering to identify putative cell types. Because each section was processed independently, the number of clusters varied between tissue sections. For network analysis, gene pairs demonstrating co-expression (p-value ≤ 0.05) in at least 2 biological or technical replicates for each dose were considered as directly connected (2 nodes 1 edge). A network was drawn from the interactions showing each gene (node) and interaction (edge) at each dose using igraph v1.2.7. From the network, the number of edges between each pair of nodes was calculated (closeness) and used to identify a spatially resolved network of co-expressed genes.

## RESULTS

### Clustering and cell type annotation

snRNAseq was used to further investigate the dose-dependent effects of TCDD on cell-specific differential gene expression. Doses were selected to induce complete gene expression dose response curves as well as the previously reported spectrum of pathologies using the same dosing regimen to elicit lipid accumulation (≥ 0.3 μg/kg), inflammation (≥ 3 μg/kg), and fibrosis (30 μg/kg) (Fader *et al.*, 2017a; Fader *et al.*, 2017b; Nault *et al.*, 2016). At 30 μg/kg, TCDD modestly increased serum ALT levels with body weight loss ≤ 15% within treatment groups suggesting minimal overt toxicity following oral gavage every 4 days for 28 days (Nault *et al.*, 2016). A total of 131,613 nuclei passed quality control across all samples with an average of 14,624 (ranged from 8,717 to 18,131 nuclei) per dose (**Table S2**). Approximately 1,665 differentially expressed genes were expressed across all nuclei with a median of 3,294 unique molecular identifiers (UMI; transcripts). An average of 18,317 genes were detected in individual samples with negligible mitochondrial gene contamination. There was a noticeable representation of long non-coding RNAs, consistent with our previous independent snRNAseq study (Nault *et al.*, 2021). Conversion of this snRNAseq data to pseudobulk, for comparison to a published bulk RNAseq dataset (GSE203302) that were taken from the same liver samples, demonstrated good agreement between both technologies for the identification of differentially expressed genes (**Figure S2**). Among the genes that showed poor correlation in response to TCDD were low abundance genes (*e.g., Atp1a2*) and previously reported cytosol biased genes (*e.g.*, *Pgm1, Pgm2*, and *Slc2a4*) (Bahar Halpern *et al.*, 2015) (**Table S3**). Small differences with the use of different technologies and sample source (tissue vs nuclei) is not unexpected.

Data dimensionality reduction and clustering was used to assess hepatic cell populations at various levels of resolutions. Since clustering is influenced by the dataset size, unique assignments were made at various resolution levels to identify the optimal parameters to distinguish unique cell types (**Figure S1**). A resolution of 0.1 produced 11 distinct clusters that identified the previously characterized liver cell types (Halpern *et al.*, 2018; Halpern *et al.*, 2017; Nault *et al.*, 2021; Xiong *et al.*, 2019). A semi-automated strategy that involved label transferring based on published data complemented by manual cluster identification using marker genes as outline in the Materials and Methods was used to annotate clusters as specific cell types (**Figure 1A-B**). In addition to the 9 previously identified cell types, two potentially new cell types were identified. By comparison to other hepatic datasets and examining marker gene expression, one cluster was identified as a dendritic cell type (pDCs) based on the expression of *Fnbp1, Wdfy4*, and *Ciita* markers (**Figures 1C, S3**). The other novel cluster was identified as portal fibroblasts based on *Gas6* and *Msln* marker expression (Koyama *et al.*, 2017).

**Figure 1.**
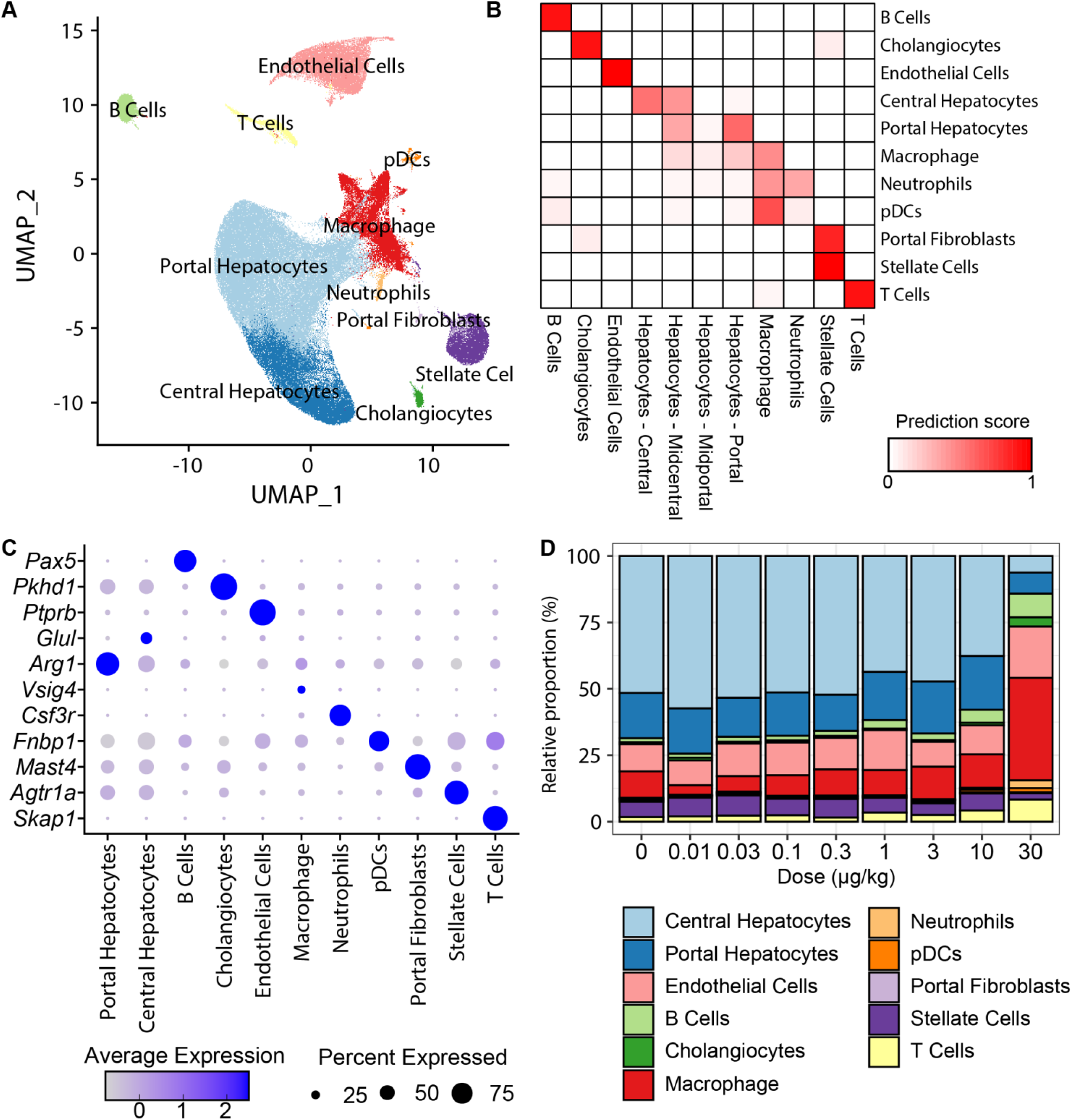
Clustering and annotation of snRNAseq data from liver samples of male mice gavaged with TCDD every 4 days for 28 days. (A) UMAP visualization of 131,613 (N = 2 – 3 biological replicates) annotated nuclei expression profiles across all doses and treatments. (B) Label transfer prediction scores for each cluster was estimated based on published hepatic snRNAseq data (GSE148339). (C) Dot plot of marker expression distinguishing individual clusters where the color represents average normalized expression and size represents percent of nuclei expressing the marker gene. (D) Relative proportion of individual cell types across TCDD dose levels. Colors represent distinct cell types matched to UMAP colors in (A). The bar size represents mean values (N = 2 – 3 biological replicates).

TCDD dose-dependently altered the relative proportions for liver cell types (**Figure 1D**). Hepatocytes (all zones) comprised the most abundant cell type representing 68.6 ± 2.0% in vehicle samples. However, in treated liver samples, hepatocytes (all zones) only represented 14.2 ± 5.3% following treatment with 30 μg/kg TCDD. HSCs also exhibited a modest decrease. In contrast, macrophages which only made up 9.9% ± 2.4 of hepatic cells in control livers increased to 38.6 ± 3.5% after treatment with 30 μg/kg TCDD. This large change in relative cell population is consistent with the reported increase in F4/80 staining in TCDD treated mice (Fader *et al.*, 2017b; Li *et al.*, 2020). In addition, the relative proportions of cholangiocytes, LSECs, B cells, T cells, and neutrophils dose-dependently increased. The proportion of pDCs (0.5% - 1.5%), and portal fibroblasts (0.4% – 0.6%) did not change, though assessing population changes for rarer cell types was more difficult.

#### Differential gene expression and functional enrichment

The number of differentially expressed genes (DEGs) (Bayes Factor adjusted false discovery rate (FDR) ≥ 0.05 and |fold-change| ≥ 1.5) for individual cell types ranged from 368 in portal fibroblasts to 1,339 in macrophages. In agreement with our previous report (Nault *et al.*, 2021), central hepatocytes were more responsive with 900 DEGs compared to 734 in portal hepatocytes. We examined DEGs for cell-specific (only differentially expressed in 1 cell type) or common (differentially expressed in two or more cell types) DEGs (**Figure 2A**). The largest sets were primarily cell specific DEGs with only 84 DEGs shared across all cell types. The 84 genes largely consisted of known AhR target genes including *Cyp1a1* (7.8-fold; central hepatocytes), *Cyp1a2* (8.6-fold; central hepatocytes), *Fmo3* (26.2-fold; central hepatocytes), *Tiparp* (5.8-fold; central hepatocytes), and *Nfe2l2* (7.6-fold; central hepatocytes) (**Figure S4; Table S4**). The next largest multiple cell type DEG intersections occurred between macrophages and pDCs (91 DEGs), both derived from the myeloid lineage, followed by portal and central hepatocytes (80 DEGs) consistent with the clustering in **Figure S1**.

**Figure 2.**
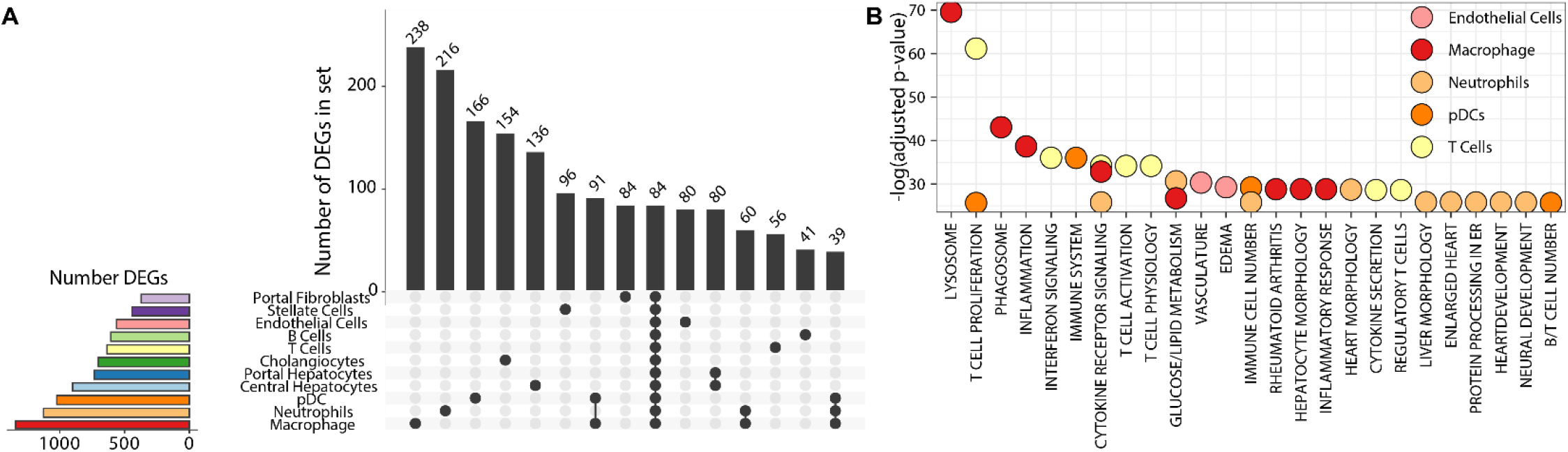
Set and functional analysis of hepatic differentially expressed genes (DEGs) from male mice gavaged with TCDD every 4 days for 28 days. (A) UpSet plot of the 15 largest gene sets, in rank order, based on set analysis that identified both unique and common differentially expressed genes (DEGs; Bayes Factor adjusted FDR ≥ 0.05 and |fold-change| ≥ 1.5) among all identified cell types. Set sizes represent the number of genes identified in only the cells as indicated by black circles. (B) Functional analysis of DEGs for each cell type using gene lists from KEGG and MPO from the Gene Set Knowledgebase (GSKB; http://ge-lab.org/gskb/). Gene sets with ≥ 60% overlap were combined and manually annotated. The top 30 enriched functions (adjusted p-value) across all cell types are shown. A complete list is available in **Table S2**.

Differential gene expression within individual cell types reflected their specific physiological roles such as phagocytosis in macrophages, the proliferation and differentiation of B cells, T cells, and pDCs, the vascular function of LSECs, and the expression of extracellular matrix related genes by HSCs (**Figure 2B; Table S5**). Interestingly, macrophages and neutrophil DEGs were also enriched for glucose/lipid metabolism. This included *Acadm* (0.74-fold), *Atf6* (0.69-fold), *Clock* (0.52-fold), *Hsd17b4* (0.61-fold), *Ldlr* (0.54-fold), *Pex7* (0.70-fold), *Ppargc1a (0.64-fold), Pten* (0.58-fold) in neutrophils, while *Lipa* (1.79-fold), *Nr1h3* (1.51-fold), *Pparg* (1.58-fold), and *Yap1* (0.66-fold) were differentially expressed in macrophages (**Figure S5**). Neutrophils have previously been linked to hepatic glucose and lipid homeostasis (Ou *et al.*, 2017) consistent with abnormal glucose metabolism gene expression in neutrophil DEGs (**Figure 2B**).

#### Cell-cell communication

CellphoneDB was used to identify emerging cell-cell interactions following treatment with TCDD (**Figures 3, S6**). Hepatocytes and portal fibroblasts interacted with other cell types via the NRG1-ERBB4 ligand-receptor pathway (**Figure 3B**). Neuregulin-1 (*Nrg1*), an epidermal growth factor family member increases hepatic glucose uptake, inhibits gluconeogenesis, and regulates cholesterol biosynthesis (Arai *et al.*, 2017; Haskins *et al.*, 2015; Zhang *et al.*, 2018). *Nrg1* was detected in 99% of central hepatocytes with a 14.4-fold induction at 30 μg/kg TCDD while *Erbb4* was induced 5.8-fold and detected in 97% of neutrophils at 3 μg/kg TCDD (**Figure 3C**).

**Figure 3.**
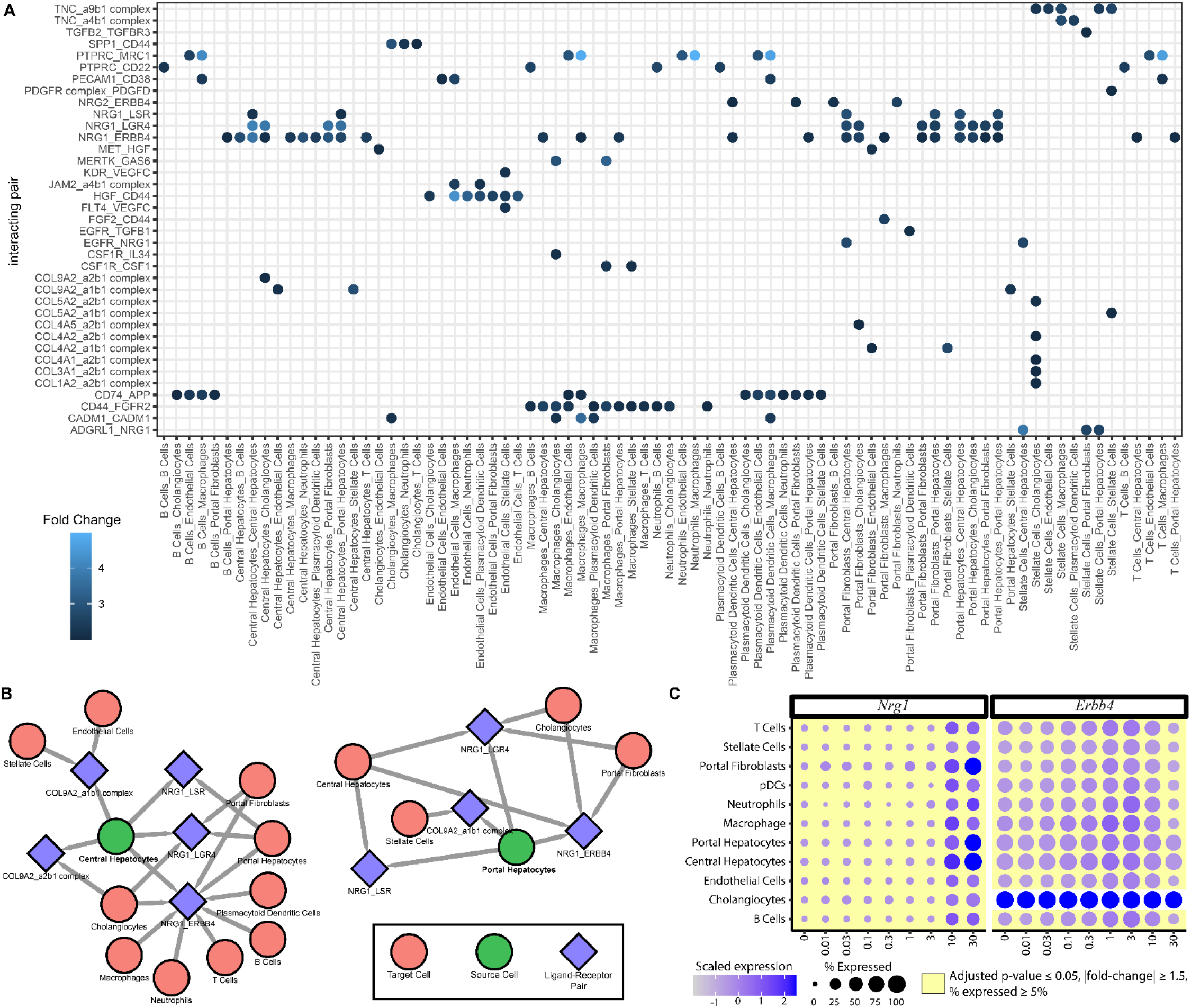
Cell-cell interaction analysis between liver cell (sub)types in male mice gavaged with TCDD every 4 days for 28 days. (A) Ligand-receptor interacting pairs were identified using CellPhoneDB at each dose including genes expressed in >25% of nuclei. Mean values for ligand-receptor pairs (mean of the means for each member of the interaction) were calculated at each dose and only interactions exhibiting a ≥ 2-fold increase relative to control and p-value ≤ 0.05 in any dose group are shown. Cell combinations are shown as the cell expressing the ligand (source) followed by the cell expressing the receptor (target). (B) The network of interactions is shown for central and portal hepatocytes (green nodes). (C) Expression of *Nrg1* and *Erbb4*. The color of the dot represents the average normalized expression, and the size of the dot represents the percent of nuclei expressing the gene. Each dose group consists of 2 – 3 biological replicates representing a total of 131,613 nuclei.

LSECs communicate with cholangiocytes, macrophages, HSCs, portal fibroblasts, T cells, and neutrophils using hepatocyte growth factor (HGF) and the cell surface receptor, CD44. Endothelial *Hgf* was induced 4.0-fold but repressed 1.5-fold in stellate cells. CD44 is an important factor in NAFLD progression, with its deletion reducing fibrosis and infiltration consistent with its presence in fibroblasts (portal fibroblasts and HSCs; up to 1.63-fold) as well as immune cells (macrophages, T cells, and neutrophils; up to 4.6-fold). LSECs also interacted with VEGFC in HSCs (*Vegfc;* 3.1-fold) via production of KDR (*Kdr;* 1.6-fold) and FLT4 (*Flt4;* 1.7-fold) ligands. Stellate cells are also associated with extracellular matrix and cell-cell contact through collagens (*Col1a2, Col3a1, Col4a1, Col4a2, Col5a2*; ≥2-fold induction) and TNC (*Tnc;* 4.0-fold) with alpha-2-macroglobulin (A2M) complexes, indications of fibrosis and protease inhibition (Chrostek and Panasiuk, 2014; Ratziu *et al.*, 2006). In addition, HSCs were found to interact with portal fibroblasts through TGFB2-TGFBR3 and ADGRL1-NRG1 (**Figure S6**). Interestingly, TGF-β signaling promotes portal fibroblast differentiation into HSCs *in vitro* (Li *et al.*, 2007). SPP1 (*Spp1* aka Osteopontin; 1.9-fold) released from cholangiocytes also contributed to cell interactions and is implicated in NAFLD and hepatocellular carcinoma (Song *et al.*, 2021).

Immune cells largely interact through PTPRC (CD45; 5.3-fold) and PECAM1 (1.8-fold) with other cell surface receptors for recruitment and maintenance. B-cells, pDCs, and macrophages were also found to interact with cholangiocytes and LSECs via CD74 (*Cd74;* 2.6-fold in pDCs) and amyloid precursor protein (*App;* 2.7-fold in macrophages). *In vitro*, CD74 inhibits β-amyloid (Aβ) production by preventing APP processing resulting in endocytosis (Matsuda *et al.*, 2009). Hepatic Aβ maintains stellate cell quiescence by suppressing α-SMA, collagen, and TGF-β expression while inducing nitric oxide (NO) production (Buniatian *et al.*, 2020). Moreover, APP is linked to bile duct blockage and biliary atresia consistent with TCDD treatment increasing gall bladder volume and bile duct proliferation in mice (Babu *et al.*, 2020; Fader *et al.*, 2017b). Macrophage also interact with portal fibroblasts and cholangiocytes via CSF1R (CSF1R-CSF1 and CSF1R-IL34) and MERTK (MERTK-GAS6). Hepatic activation of CSF1R (*Csf1r*; 3.3-fold) promotes macrophage proliferation and activation consistent with the increased macrophage populations in TCDD treated livers (Nault *et al.*, 2021). These interactions further highlight the complex, dose-dependent interactions associated with the hepatotoxicity and progression of steatosis to steatohepatitis with fibrosis elicited by TCDD.

#### Spatial heterogeneity in hepatic gene expression

To investigate spatially resolved gene expression cell segmentation of molecular cartography analyses was examined using Baysor while spatial heterogeneity was characterized by MERINGUE (Miller *et al.*, 2021; Petukhov *et al.*, 2021). Individual cell types were identified based on marker gene expression with cell-cell relationships determined using network analysis (**Figures 4, S7**). For example, the central hepatocyte marker *Glul* was largely associated with nuclear receptors (*e.g., Nfe2l2, Nr1h4* [FXR], *Nr1i2* [PXR], *Nr1i3* [CAR], *Ppard*, and *Rxra*), xenobiotic metabolism (*e.g.*, *Ahr, Gsta3, Gstm3*, and *Nqo1*), and lipid cholesterol metabolism (*e.g.*, *Aldh3a2*, *Cd36*, *Ces1b*, *Cyp7a1*, *Hmgcs1*, *Mgll*, *Sqle*, *Srebf1*, and *Vldlr*) genes at lower doses (0 – 3 μg/kg). Conversely, the portal hepatocyte marker *Cyp2f2* was associated with *Cbs*, *Egfr, G6pc, Gldc, Gls2, Hal, Kynu, Pkhd1*, and *Sds* as expected with the portal region receiving nutrient- and oxygen-rich blood requiring antioxidant defenses (*Cbs*, *Gldc*). In addition, the portal zone is responsible for amino acid (*Hal, Gls2*) and glucose metabolism (*G6pc*). *Cyp2f2* was also associated with cholangiocyte markers *Sox9* and *Pkhd1* consistent with their co-localization in the portal triad. The HSC marker, *Tagln*, co-localized with the extracellular matrix expression (*e.g.*, *Adamtsl2, Col14a1, Col1a1, Col1a2, Col3a1*, and *Des*), Wnt signaling (*e.g.*, *Cacna2d1*, and *Ccnd1*), and LSECs (*e.g.*, *Rspo3*) genes. Similarly, at 0 and 0.3 μg/kg TCDD, macrophage markers were adjacent to HSCs in agreement with a previous report of a cell-cell interaction (Bonnardel *et al.*, 2019). With increasing dose, these spatial delineations eroded and became more ambiguous (**Figure 4D**). For example, *Cyp2f2* was adjacent to genes associated with xenobiotic metabolism genes such as *Gsta3*, nuclear receptors (*e.g.*, *Pparg, Ppara*, and *Nr1i3* [CAR], *Nfe2l2*, and *Ghr*), and other genes that typically co-localized with *Glul* at lower doses. Overall, genes distant from each other in control samples (**Figure 4A**) become more closely associated as indicated by the increased area with greater blue intensity (**Figure 4D).** The data indicates a loss of zonal gene expression across the portal to central axis.

**Figure 4.**
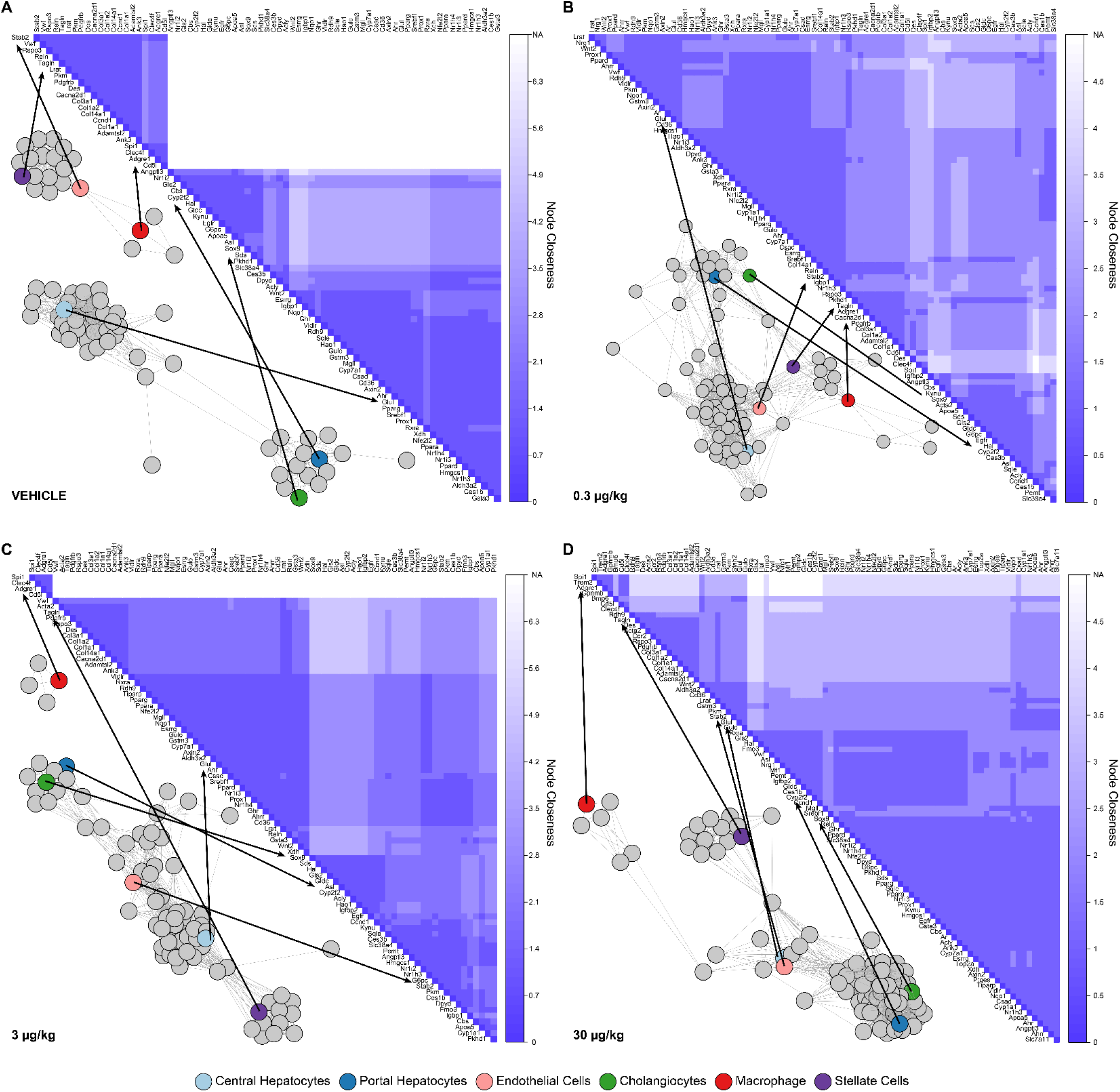
Network visualization of spatially resolved hepatic gene expression determined using MERINGUE and the Resolve Biosciences Molecular Cartography system. Mice were treated with (A) sesame oil vehicle control, (B) 0.3, (C) 3, or (D) 30 μg/kg TCDD every 4 days for 28 days. Correlations between gene co-expression was determined based on co-expression in at least 2 biological or technical replicates for each dose. Gene pairs that exhibited co-expression were considered directly connected are indicated by 2 nodes and 1 edge. Each gene (node) and interaction (edge) was used to draw a network using igraph v1.2.7. From the network, the number of edges between each pair of nodes was calculated (closeness) and used to identify a spatially resolved network of co-expressed genes. A closeness of 0 indicates nodes that could not be connected at all (*i.e.*, independent clusters; area is white). A marker gene for individual cell types is colored to illustrate their position within the network. Arrows show the location of marker genes in the associated heatmap where the intensity of the blue color represents the closeness of the genes.

#### Dose-dependent loss of hepatocyte zonal identity

Hepatocyte nuclei were extracted from the complete nuclei dataset and re-processed independently using spatially resolved genes for dimensionality reduction (Halpern *et al.*, 2017; Nault *et al.*, 2021). Two clear clusters were identified by UMAP visualization with the leftmost cluster enriched for the central hepatocyte markers, *Glul* and *Gulo*, while the portal markers, *Cyp2f2* and *Sds*, were over-represented in the rightmost clusters (**Figures 5A, C, S8**). In control mice, *Igfbp2*, considered a midzonal marker with a portal bias, was also more abundant in the rightmost cluster with modest expression in the leftmost cluster consistent with its midzonal expression (**Figure S8**). Trajectory analysis examined the continuum from central to portal zones following the patterns of marker expression. Spatial transcriptomics corroborated the zonal hepatocyte gene expression, showing a clear separation of *Glul* expressing hepatocytes from *Cyp2f2* expressing hepatocytes (**Figures 5B, D**).

**Figure 5.**
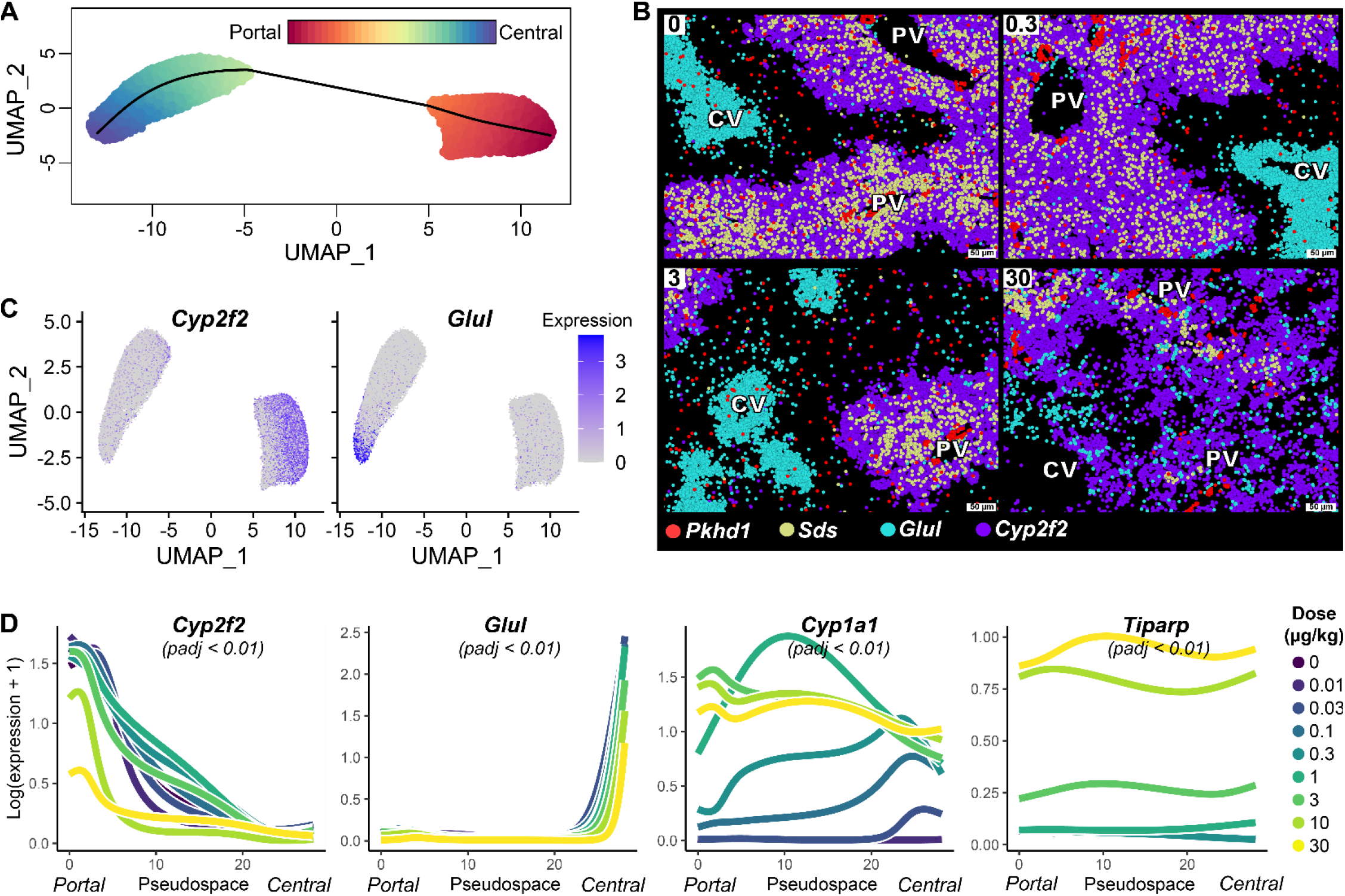
Effect of TCDD on zonal gene expression in the male mouse liver following oral gavage with TCDD every 4 days for 28 days. The hepatocyte nuclei subset was extracted from the snRNAseq dataset and re-integrated using spatially resolved genes. (A) Slingshot was used to evaluate the continuum from central to portal hepatocytes shown as a color gradient with the average represented by the black line. (B) Molecular cartography was used to visualize the spatial distribution of *Pkhd1*, Sds, Glul and Cyp2f2 zonal marker genes. Colors represent individual genes and individual spots reflect a single transcript. (C) UMAP visualization shows the expression of the portal (*Cyp2f2*) and central (*Glul*) hepatocyte markers. (D) Dose dependent expression of portal *Cyp2f2* and central *Glul* as well as AHR target genes (*Cyp1a1* and *Tiparp*) along the pseudospace continuum from portal to central zones. Adjusted p-values were calculated based on a Wald statistic for the TCDD treatment effect. Data represents 2 – 3 biological replicates per dose for snRNAseq and spatial transcriptomic images are representative of data obtained from 2 biological replicates and 2 technical replicates.

Examination of the distribution along the hepatocyte pseudospatial trajectory showed an increased proportion of hepatocytes with gene expression indicative of central hepatocytes suggesting a loss of portal hepatocyte identity and function (**Figure S8**). The spatial expression of several genes such as antioxidant, phase I, and II metabolism genes was changed following AHR activation by TCDD with their expression pattern disrupted compared to the typical localization observed in control lobules (**Figures 5, S9**). Specifically, *Cyp2f2*, the portal hepatocyte landmark, was dose-dependently repressed by TCDD in portal hepatocytes but was unchanged in central hepatocytes. Likewise, expression of the central hepatocyte landmark, *Glul*, was dose-dependently repressed by TCDD suggesting a loss of some central characteristics and functions in the central region (**Figure 5C**). Similarly, AHR target genes *Xdh* and *Gstm3*, which are primarily expressed in the central region in controls, were induced in all zones giving portal hepatocytes central characteristics (**Figure S9**). Although, spatially resolved transcriptomics indicated that marker genes such as *Cyp2f2* and *Glul* no longer exhibit definitive zonal distribution at 30 ug/kg TCDD, others including *Sds* and *Igfbp2* localized adjacent to portal triads marked by *Pkhd1* (cholangiocytes) revealing some zonal expression is preserved following treatment with TCDD (**Figures 5B**, **S8**). AhR target genes such as *Cyp1a1* and *Tiparp* also exhibited clear zone-specific differential expression with central induction at lower doses while snRNAseq and spatial transcriptomic analyses showed panacinar induction at higher doses (**Figure 5D**).

Using CoE to calculate pseudospatial peak expression at each dose (Halpern *et al.*, 2017), the maximum distance travelled was calculated as described in the Materials and Methods. Overall, TCDD-elicited a bias towards portal and midzonal (ΔCoE ≥ 0.5) characteristics (**Figure 6A**). Two broad gene clusters were identified: (GA) portal genes displaying more homogenous expression across all zones and (GB) homogenously expressed genes across all zones exhibiting a more biased central expression pattern. The GB cluster consisted of known AHR induced genes including *Cyp1a1, Ahrr, Tiparp, Fabp12*, and *Gk.* No GB genes were predominantly expressed in the central region in control animals and in the portal region at higher doses. *B4galt6* exhibited the largest ΔCoE (1.3; **Figure 6B**) while *B4galt5* also showed zone specific differential expression. Both *B4galt5* and *B4galt6* are implicated in lactosylceramide (LacCer) synthesis, a lipid associated with NASH, though only a modest increase in LacCer was observed at 3 μg/kg using a similar study design (Nault *et al.*, 2017b). Genes in group GA were largely repressed to levels comparable to the central region consistent with an overall loss of zonal functional organization. For example, *Acly*, is primarily expressed in the portal/midzonal region in control mice but was repressed to near central region levels starting at 3 μg/kg (**Figures 6C-D**) that likely contributed to impaired lipogenesis in the region that received the most nutrient rich blood. Overall, these results demonstrated TCDD dose dependently disrupted zonal gene expression.

**Figure 6.**
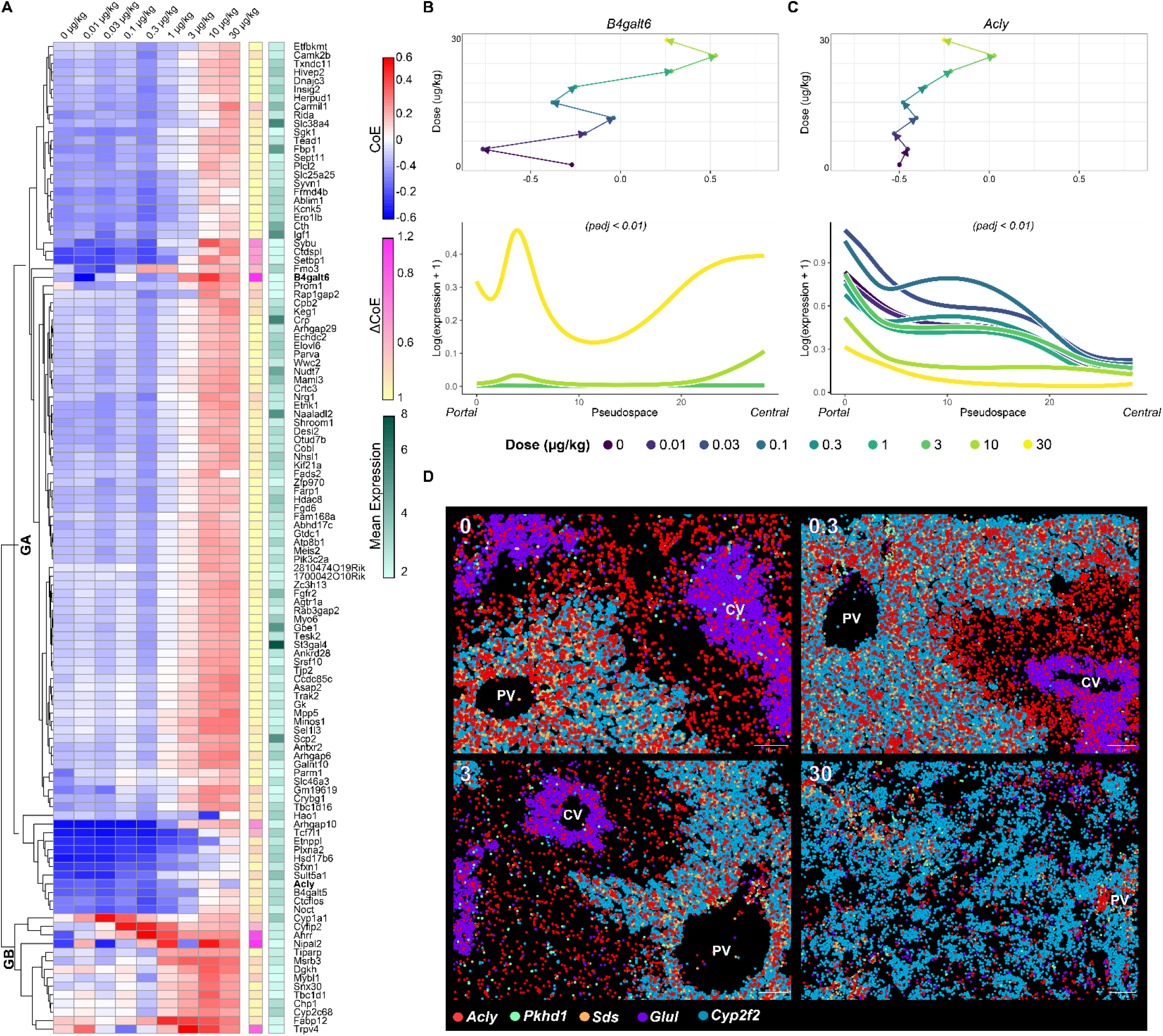
Center of Expression (CoE) analysis of hepatocyte gene expression in male mice gavaged with sesame oil vehicle control, 0.01, 0.03, 0.1, 0.3, 1, 3, 10, or 30 μg/kg TCDD every 4 days for 28 days. CoE was calculated as previously described (Halpern *et al.*, 2017). The CoE travel distance (ΔCoE; maximum CoE – minimum CoE) was calculated for each gene. (A) Hierarchical clustering identified two major clusters (GA and GB) for the top 5^th^ percentile of ΔCoE values. Dose-dependent changes of CoE values (top) and pseudospace distribution of expression (bottom) is shown for (B) *B6galt6* and (C) *Acly*. Adjusted p-values were calculated based on a Wald statistic for an effect of TCDD treatment. (D) Dose pendent molecular cartography analysis for *Acly* as well as the portal (*Cyp2f2, Sds*), central (*Glul*), and cholangiocytes (*Pkhd1*) marker genes. Scale bar represents 50 μm. Data represents 2 – 3 biological replicates per dose for snRNAseq and spatial transcriptomic images are representative of data obtained from 2 biological replicates and 2 technical replicates.

#### Disruption of the Wnt/β-catenin signaling cascade

Wnt/β-catenin signaling is a key mediator of liver zonation (Dobie *et al.*, 2019; Rocha *et al.*, 2015; Xiong *et al.*, 2019). Examination of genes involved in the Wnt/β-catenin signaling cascade showed dose-dependent and zone-specific dysregulation (**Figures 7, S10-11**). R-spondins (*Rspo1* and *Rspo3*) are ligand agonists of Wnt/β-catenin signaling that were highly expressed in portal fibroblasts, LSECs, and HSCs (**Figure 7A**). TCDD did not alter LSEC or HSC *Rspo* expression though *Rspo3*, which is highly expressed in stellate cells, was induced 1.90-fold but did not achieve significance due to the cytosolic bias of *Rspo3* mRNA, and the dose-dependent decrease in the number of stellate cells that limited the utility of the scBT test method (Bahar Halpern *et al.*, 2015) (Nault *et al.*, 2022). LRT linear analysis, a more appropriate test, identified significant *Rspo3* induction in stellate cells consistent with our spatial transcriptomic analysis (Nault *et al.*, 2022) (**Figure S11**). The induction of *Rspo3* following stellate cell activation by carbon tetrachloride is also reported in areas of fibrosis (Dobie *et al.*, 2019; Zhang *et al.*, 2020). The most highly expressed WNT ligands, *Wnt2* (1.4-fold, adjusted p-value ≤ 0.05) and *Wnt9a* (1.3-fold, adjusted p-value ≤ 0.05), were expressed in LSECs and portal fibroblasts with modest induction by TCDD that did not meet the fold-change threshold (**Figure 7A**). *Wnt2* showed both AHR enrichment and the presence of a putative DRE implicating potential direct AHR regulation (**Figure S12**).

**Figure 7.**
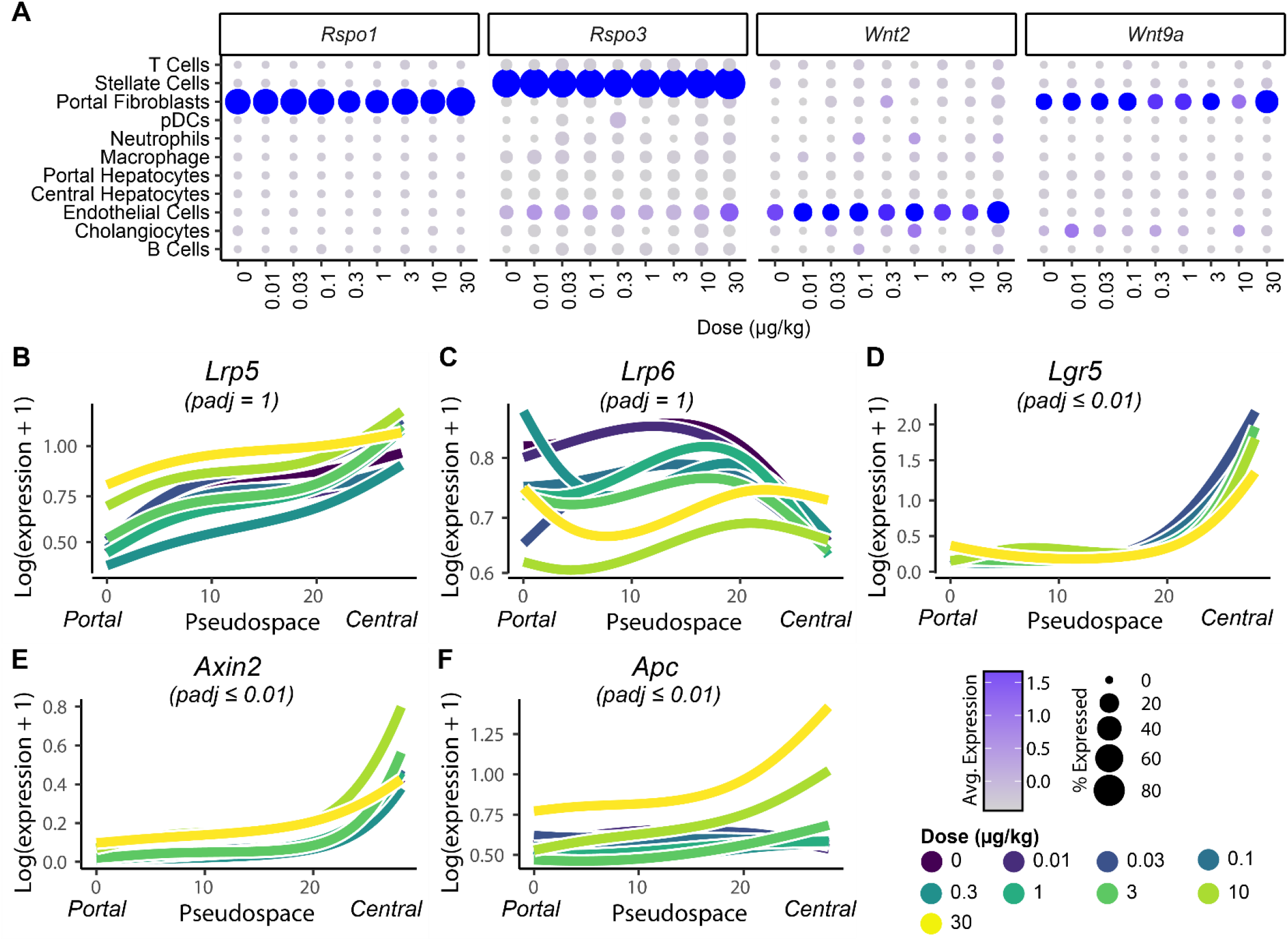
Dose dependent effects of TCDD on the cell-specific and pseudospatial expression of genes associated with Wnt/β-catenin signaling. (A) Expression of highly expressed Wnt/β-catenin ligands primarily expressed in nonparenchymal cells where the color of the dot represents the average normalized expression, and the size of the dot represents percent of nuclei expressing the gene. Dose-dependent expression along the hepatocyte pseudospace continuum is shown for receptors and downstream targets (B) *Lrp5*, (C) *Lrp6*, (D) *Lgr5*, (E) *Axin2* and (F) *Apc*. Adjusted p-values were calculated based on a Wald statistic for an effect of TCDD treatment. Data represents 2 – 3 biological replicates per dose.

The Frizzled family of receptors (*Fzd1, 3, 4, 5, 6, 7*) exhibited low levels of expression in hepatic nuclei with negligible changes following TCDD treatment. Conversely, the co-receptors *Lrp5* and *Lrp6* were highly expressed in central and portal hepatocytes, respectively, with *Lrp1* and *Lrp4* evenly distributed across all zones (**Figures 7B-C, S10**). *Lrp1* was modestly repressed in central hepatocytes while *Lrp4* was induced in both portal and central hepatocytes. While R-spondin receptors potentiate WNT signaling, *Lgr4* and *Lgr5* were the most abundantly expressed receptors in portal and central hepatocytes, respectively. *Lgr4* was not altered by TCDD while *Lgr5* exhibited modest dose dependent repression in the central zone (**Figures 7D, S10**). In agreement with altered zone-specific Wnt/β-catenin activation, the downstream target *Axin2* which is largely restricted to central hepatocytes at lower doses, exhibited greater dose dependent induction in the central zone (**Figures 7E, S11**). Additionally, *Apc*, a member of the β-catenin destruction complex that supports portal hepatocyte maintenance was dose dependently induced in all zones with the highest induction occurring in central hepatocytes (**Figure 7F**).

Under homeostatic conditions and following liver injury, the IGFBP2-mTOR-CCND1 axis drives the proliferation of midzonal hepatocytes (Wei *et al.*, 2021). Our spatial transcriptomic analysis shows that *Ccnd1* expression at low doses is limited to the portal triad with widespread portal and central expression at 30 μg/kg TCDD (**Figure S11**). While *Igfbp2* and *Ccnd1* co-localize, *Ccnd1* is more commonly co-localized with *Axin2* consistent with induction via Wnt/β-catenin signaling pathway rather than the IGFBP2-mTOR-CCND1 axis. The inhibition of liver regeneration and induction of *Ccnd1* is consistent with the perturbation of Wnt/β-catenin signaling and the G1 cell cycle arrest elicited by TCDD (Jackson *et al.*, 2014).

## DISCUSSION

The development and progression of hepatic steatosis to steatohepatitis with fibrosis involves interactions between distinct cell types as well as the disruption of the spatially organized and temporally separated metabolic functions of the liver lobule. We integrated snRNAseq and spatial transcriptomic datasets to investigate the role of AHR activation in the disruption of zonal gene expression in liver lobules following dose-dependent treatment with TCDD. In contrast to our previous report that examined only 16,015 nuclei (Nault *et al.*, 2021), pDCs and portal fibroblasts were also identified using this larger dose response snRNAseq dataset that included 131,613 nuclei. Although pDCs did not represent a significant proportion of cells in the mouse liver, increased numbers are reported in acute decompensated cirrhosis samples with higher levels of interferon (Cardoso *et al.*, 2021). Portal fibroblasts play a key role in maintaining the portal tract and produce extracellular matrix when activated (Karin *et al.*, 2016). However, the most striking cell population change was the increased proportion of macrophages from 10% in controls to 39% in treated liver samples. While the nuclei isolation protocol may be biased (*e.g.*, preferably capture macrophage nuclei), the increase in macrophages is consistent with increased F4/80 staining as well as other reports of immune cell infiltration in NAFLD models (Fader *et al.*, 2017b; Li *et al.*, 2020; Miura *et al.*, 2012; Xiong *et al.*, 2019).

Functional analysis of differentially expressed genes was in agreement with immune cells playing a central role in TCDD-elicited NAFLD pathologies. Macrophages, neutrophils, and pDCs were the three most responsive cell types followed by hepatocytes. Enrichment of lysosomal related genes in macrophages is consistent with lysosomes contributing to the internalization of cholesterol in Kupffer cells (KCs) driven in part by *Cd36* (Bieghs *et al.*, 2012). *Cd36* is a direct target of TCDD activated AHR that leads to cholesterol and cholesterol ester accumulation in the liver (Dornbos *et al.*, 2019; Lee *et al.*, 2010; Nault *et al.*, 2017b). Foam-like KCs secrete chemokines that recruit other immune cell types including neutrophils in addition to the recruitment by leukotrienes induced by TCDD (Doskey *et al.*, 2020; Takeda *et al.*, 2017). Interestingly, neutrophil infiltration is believed to contribute to NAFLD progression due to dysregulation of diurnal rhythm. While we have previously shown that TCDD dose-dependently disrupts diurnal regulation, in this study *Clock*, a master circadian regulator, was repressed in neutrophils from treated livers (Crespo *et al.*, 2020; Fader *et al.*, 2019).

In addition to cellular heterogeneity, spatial organization is a key determinant of liver function, (Braeuning *et al.*, 2011; Gerbal-Chaloin *et al.*, 2014; Kietzmann, 2019; Soto-Gutierrez *et al.*, 2017). Although AHR activation at lower TCDD doses regulates gene and protein expression primarily in the pericentral region, steatosis and inflammation initially occur in the periportal region (Andersen *et al.*, 1997; Boverhof *et al.*, 2005; Fader *et al.*, 2017b). Moreover, snRNAseq data suggests dose dependent induction of AhR target genes including *Cyp1a1* and *Cyp1a2*, as well as antioxidant defense genes initially in central hepatocytes. Spatial transcriptomic analyses also distinguished hepatocyte zonation based on markers associated with zone-specific metabolic pathways. However, TCDD elicited a dose-dependent loss of zonal identity. For example, *Acly* and *Cyp2f2*, two portal markers, were dose-dependently repressed. Nevertheless, other markers maintained their zone-specific expression such as the portal expression of serine dehydratase (*Sds*) associated with gluconeogenesis. This suggests a functional gradient was retained for some pathways, though TCDD caused its expression to further concentrate around the portal region.

Zonation of hepatic lipid accumulation depends on the model and exhibits species differences. It is dependent on the functional differences between portal and central hepatocytes (Hijmans *et al.*, 2014; Schleicher *et al.*, 2017). In human NAFLD and alcoholic fatty liver disease (AFLD), lipids accumulate in the central region consistent with the upregulation of lipogenesis (a central biased pathway) and the uptake of mobilized lipids originating from peripheral tissue stores. In contrast, lipid accumulation in pediatric NAFLD exhibits a periportal bias or no zonal preference in the absence of hepatocyte ballooning (Carter-Kent *et al.*, 2011; Hijmans *et al.*, 2014). In *ob/ob, db/db*, and high fat diet fed mouse models, fat accumulation occurs centrally (Flach *et al.*, 2011; Hijmans *et al.*, 2014; Wiegman *et al.*, 2003), while diets high in carbohydrates and the western diet primarily induce portal fat accumulation (Ghallab *et al.*, 2021; Hijmans *et al.*, 2014). Although zonation of steatosis may be model dependent, further studies are needed to elucidate the underlying mechanisms responsible for central versus portal steatosis and whether these factors are additive in panacinar steatosis. Furthermore, mice and humans exhibit zonal differences in metabolic activity, particularly for lipogenesis related genes (Massalha *et al.*, 2020). In this study, TCDD disrupted zonation with periportal hepatocytes losing characteristic functions and adopting central functions (*i.e.*, pericentralization). More specifically, TCDD repressed β-oxidation and increased triglyceride synthesis, functions typically associated with the portal and central region, respectively (Cholico *et al.*, 2021; Lee *et al.*, 2010; Nault *et al.*, 2017b). TCDD-elicited steatosis has been attributed to dietary sources consistent with elevated levels of fatty acids in the portal circulation following feeding and not impaired metabolism in the portal region (Angrish *et al.*, 2012; Nauli and Matin, 2019; Nault *et al.*, 2017b; Schleicher *et al.*, 2017).

Several mechanisms are implicated in the distribution of enzymatic activities along the portal-central axis that determines the metabolic characteristics of hepatocytes. The Wnt/β-catenin pathway plays a key role in determining this functional distribution (Braeuning *et al.*, 2011; Gerbal-Chaloin *et al.*, 2014; Nault *et al.*, 2021; Prochazkova *et al.*, 2011; Vondracek and Machala, 2016; Yang *et al.*, 2022). Deletion of β-catenin in hepatocytes, as well as *Lgr4* and *Lgr5* deletion, increase NAFLD severity in mice fed a methionine choline deficient diet or high fat diet (Behari *et al.*, 2010; Saponara *et al.*, 2021). Abundantly expressed Wnt ligands, *Wnt2* and *Wnt9a* showed only modest induction with TCDD. *Wnt2* was identified as abundantly expressed by central LSECs with its knockout eliciting a “periportal” hepatocyte phenotype (Halpern *et al.*, 2018; Hu *et al.*, 2022). More importantly, *Wnt2* in LSECs is co-expressed with *Rspo3*, another Wnt/β-catenin signaling member implicated in the maintenance of central hepatocyte characteristics (Halpern *et al.*, 2018; Rocha *et al.*, 2015). However, in TCDD treated livers, *Rspo3* was primarily expressed and induced in stellate cells as reported in other hepatic scRNAseq datasets (Dobie *et al.*, 2019; Xiong *et al.*, 2019). Levels of R-spondins, particularly *Rspo3*, are increased in human hepatic fibrotic lesions and induced following the activation of stellate cells *in vitro.* HSCs also exhibit zonation, with central HSCs producing collagen that is associated with elevated *Rspo3* expression (Dobie *et al.*, 2019). In our dataset, very few HSCs with low *Rspo3* expression were identified possibly due to a nuclei isolation bias or given the difference in the age between mouse models (*i.e.*, 4 – 8 weeks vs. 10 – 16 weeks). Spatial transcriptomics suggested the presence of zonated HSCs further implicating AHR-mediated induction of *Rspo3* in HSCs as another factor contributing to disrupted lobular zonation following treatment with TCDD.

In summary, dose-dependent AHR activation by TCDD elicited cell- and zone-specific dysregulation of gene expression along the portal-central axis of hepatic lobules. Spatial transcriptomics confirmed the initial loss of periportal hepatocyte characteristics progressed to disruption of positionally defined functions throughout the liver lobule following treatment with higher doses of TCDD. β-Catenin activation in the portal region and impaired activation in the central region is consistent with *Apc* expression and its role as a zonal gatekeeper (Benhamouche *et al.*, 2006). The loss of portal specific functions is also in agreement with impaired lipid metabolism (*i.e.*, β-oxidation), and initial increases in lipid accumulation in the periportal zone. The progression of pathologies likely involves aberrant Wnt and R-spondin signaling within LSECs and HSCs that contributes to metabolic reprogramming and hepatotoxicity that contribute to an inflammatory environment and further tissue damage. In contrast to other hepatotoxicants such as acetaminophen and ethanol that require metabolic activation to elicit zone-specific effects, we propose TCDD and related compounds cause hepatotoxicity by altering zone specific functions that disrupt intermediate metabolism. Further studies using cell-specific and zone-specific ablation of AHR and Wnt/β-catenin pathway members are required to further explore the disruption of enzymatic functions along the portal-central axis as a novel mechanism contributing to hepatotoxicity. Collectively, this study begins to elucidate the dose-dependent cell-cell interactions elicited by TCDD that underlie the progression of steatosis to steatohepatitis with fibrosis that parallel the pathologies associated with NAFLD.

## ACKNOWLEDGEMENTS

The authors thank Lewis Vann, Sam Stingley, and Ricardo Gerreiro at Resolve Biosciences for their support with performing spatial transcriptomics analyses. This work was supported by the National Institutes of Health Science (R01ES029541), NIEHS Superfund Research Program (P42ES004911), and National Human Genome Research Institute (R21HG010789). Tim Zacharewski and Sudin Bhattacharya are partially supported by AgBioResearch at Michigan State University.

## AUTHOR CONTRIBUTIONS

**Rance Nault:** Conceptualization; Data curation; Formal analysis; Investigation; Methodology; Software; Visualization; Writing - original draft; writing - review & editing. **Satabdi Saha:** Software; Writing - review & editing. **Sudin Bhattacharya:** Funding acquisition, writing - review & editing. **Samiran Sinha:** Software; Writing - review & editing. **Tapabrata Maiti:** Software; Writing - review & editing. **Tim Zacharewski:** Conceptualization; Funding acquisition; Project administration; Resources; Supervision, Writing - review & editing.

---

**Figure S1.**
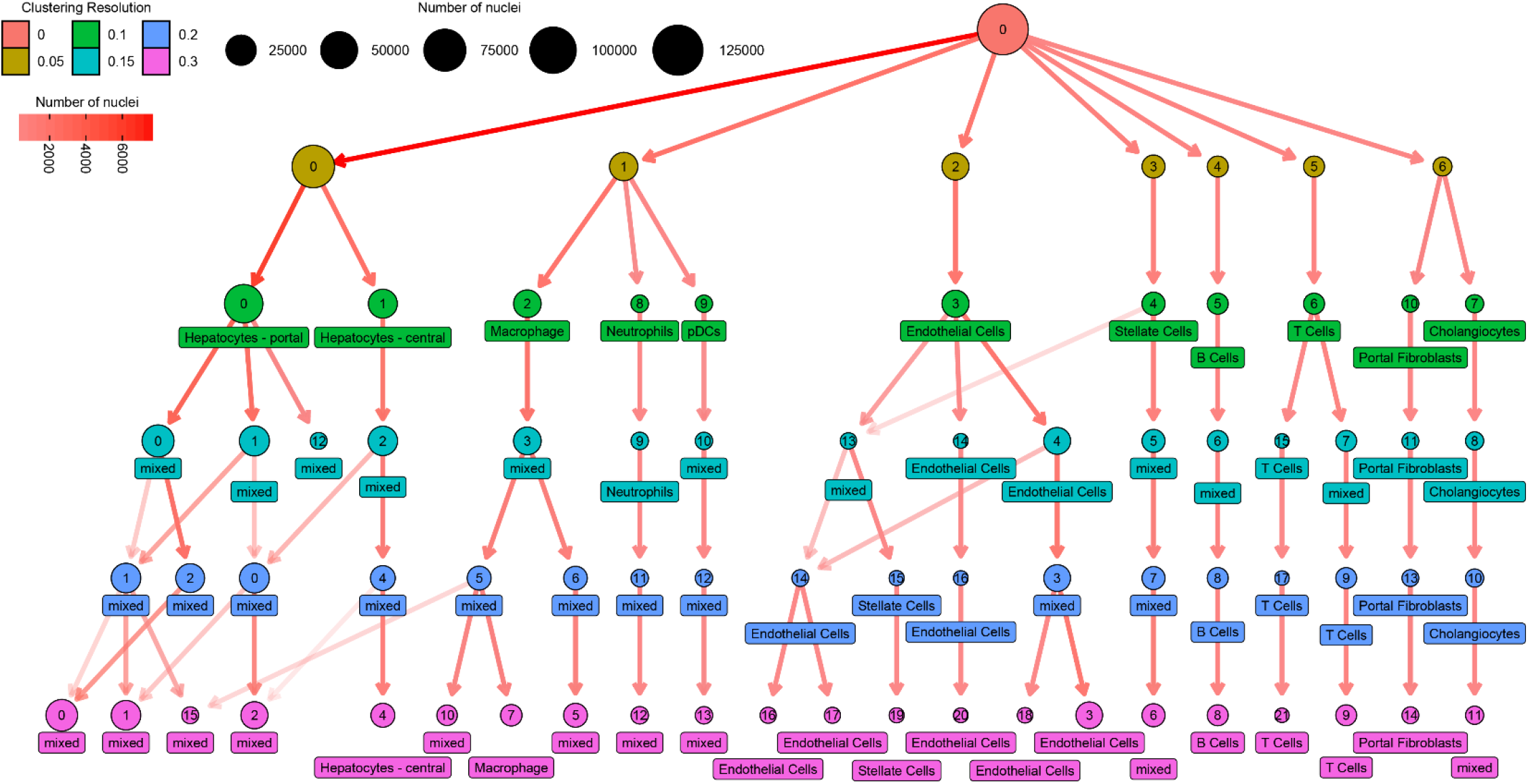
Clustering tree used to identify hepatic cell types using snRNAseq data. Seurat was used to cluster nuclei at resolutions ranging from 0 to 0.3. The number of nuclei was calculated and tracked as resolution increased to identify putative cell (sub)types. Labels were assigned at the lowest resolution where cells types could be clearly identified (0.1).

**Figure S2.**
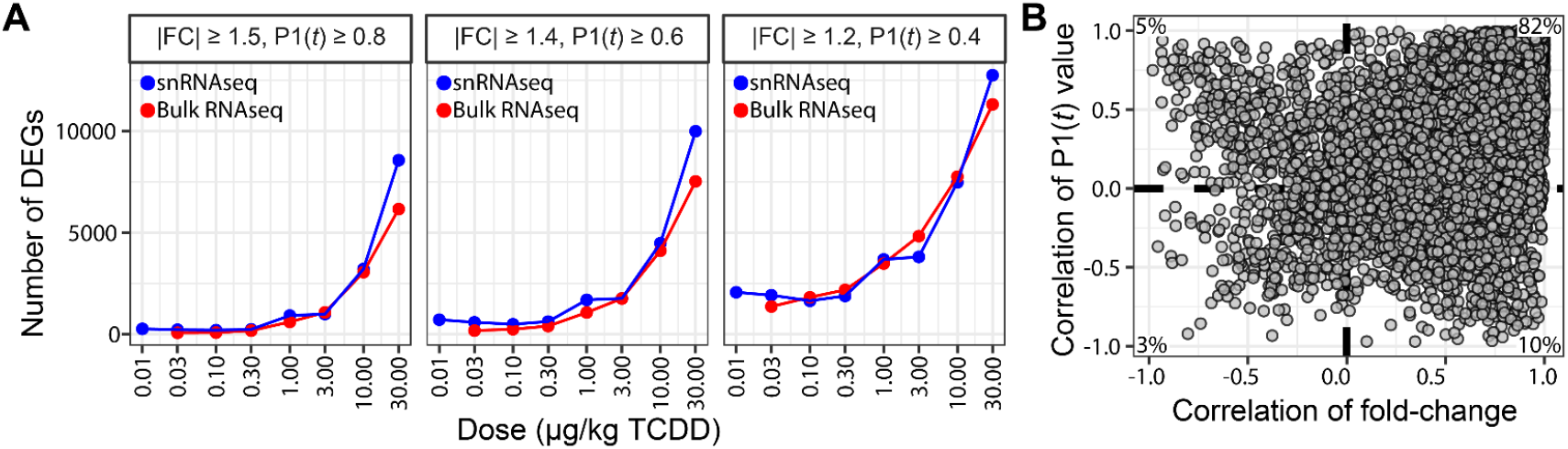
Pseudobulk data from converted dose-response snRNAseq compared to bulk RNAseq data from the same tissue. snRNAseq data was converted to pseudobulk and compared to bulk RNAseq (GSE203302) following semi-parametric normalization and empirical Bayes analysis as previously described (Nault *et al.*, 2015). (A) Differentially expressed genes were identified for each dose group at varying |fold-change| and P1(*t*) thresholds. (B) Correlation analysis of calculated fold-change and significance value (P1(*t*)) for each gene when comparing pseudobulk from snRNAseq and bulk RNAseq datasets.

**Figure S3.**
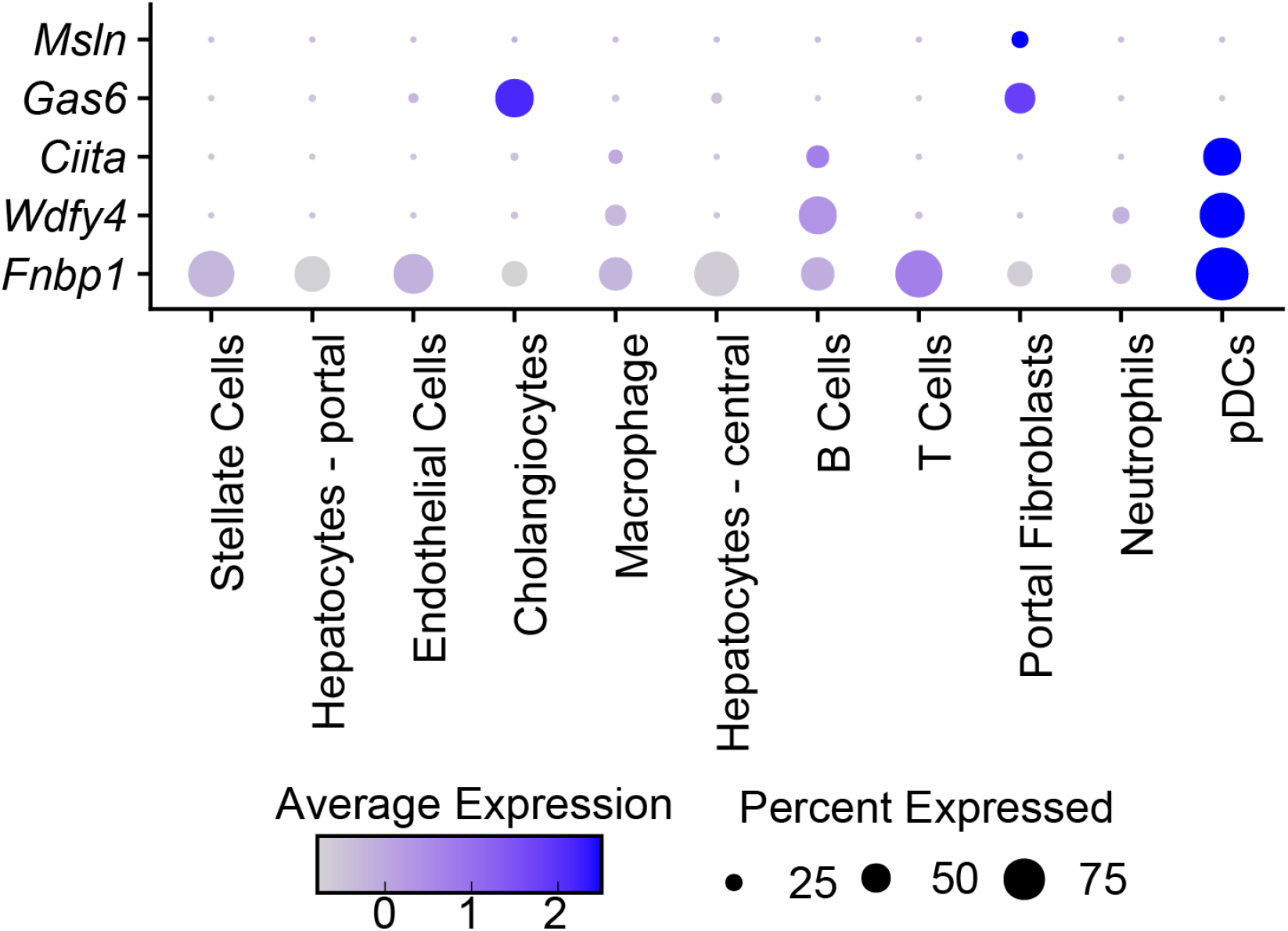
Manual identification of portal fibroblasts and plasmacytoid dendritic cells (pDCs) using marker genes. Portal fibroblasts and pDCs marker expression in individual clusters are shown as dot plot. The color of the dot represents the average normalized expression level while the size of the dot represents the percentage of nuclei expressing the gene.

**Figure S4.**
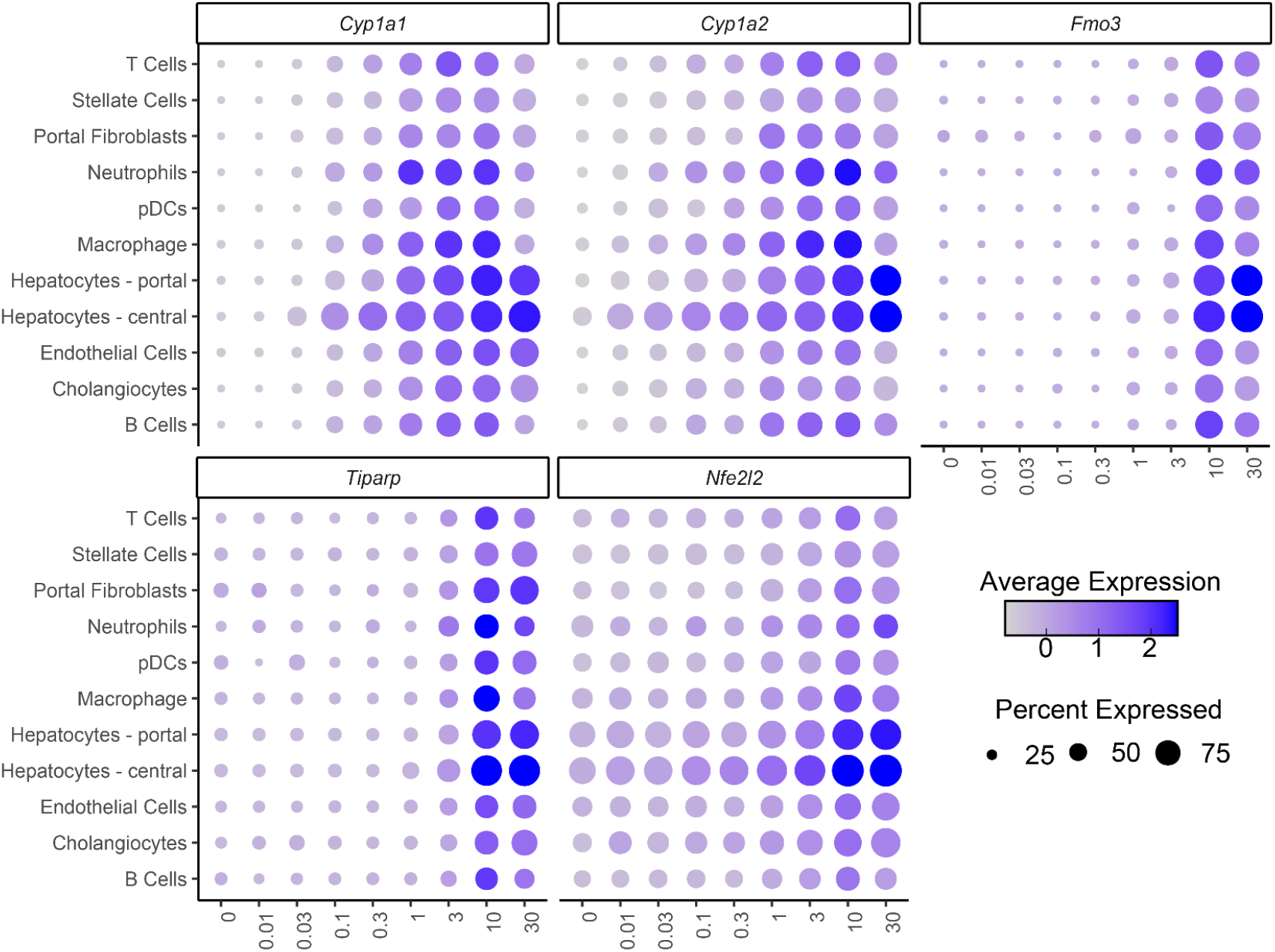
Scaled expression of AhR target genes in hepatic cells from male mice gavaged with TCDD every 4 days for 28 days. Expression of AhR regulated genes differentially expressed in specific cell types shown as dot plot where the color of the dot represents the average normalized expression, and the size of the dot represents percent of nuclei expressing the gene.

**Figure S5.**
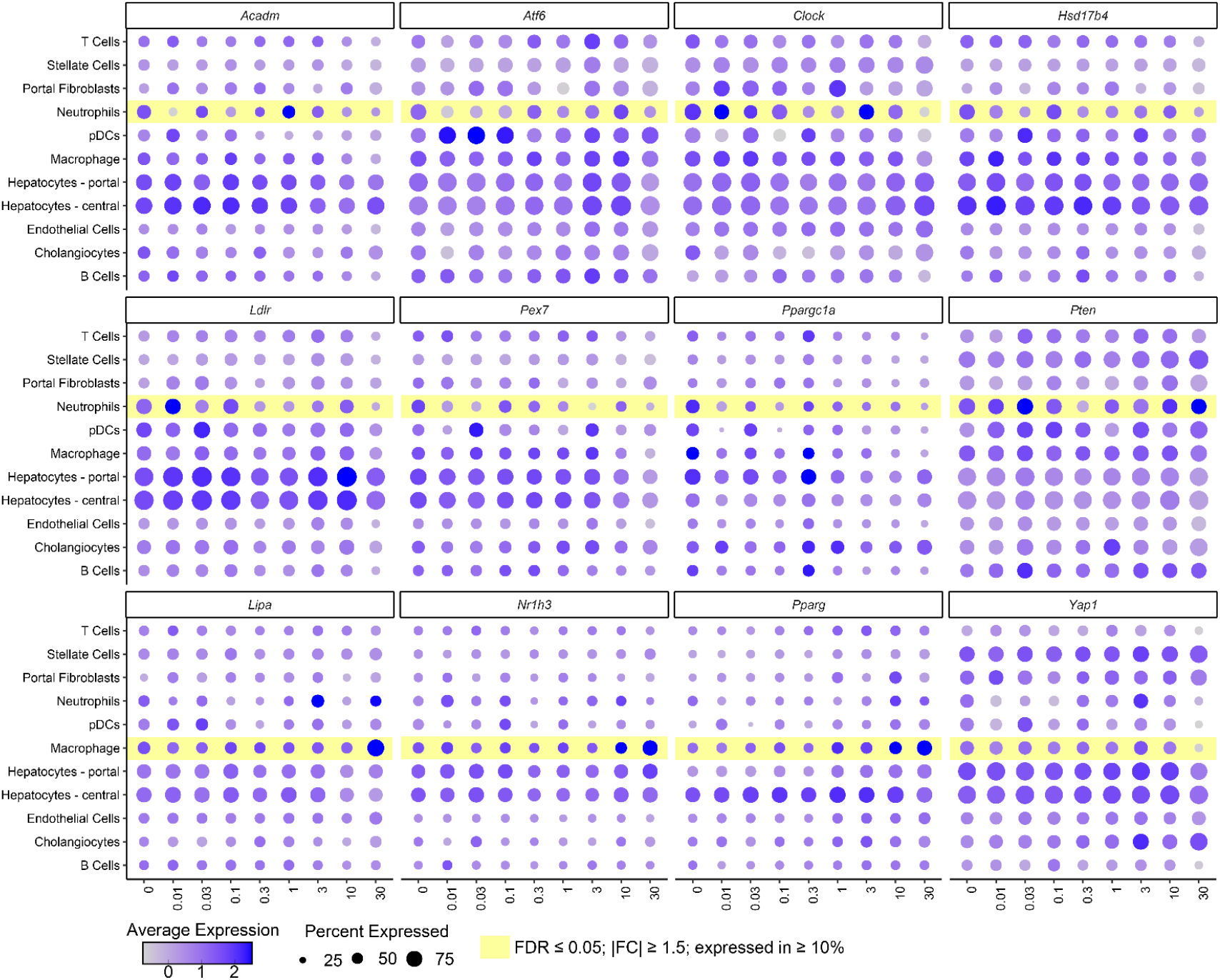
Scaled expression of DEGs associated with the steatosis gene list enriched in neutrophils and macrophages of male mice gavaged with TCDD every 4 days for 28 days. Expression of DEGs from the Mouse Phenome Ontology (MPO) Hepatic Steatosis gene set are shown as a dot plot where the color of the dot represents average normalized expression and size of the dot represents percent of nuclei expressing the gene.

**Figure S6.** See supplementary HTML file.

**Figure S7.**
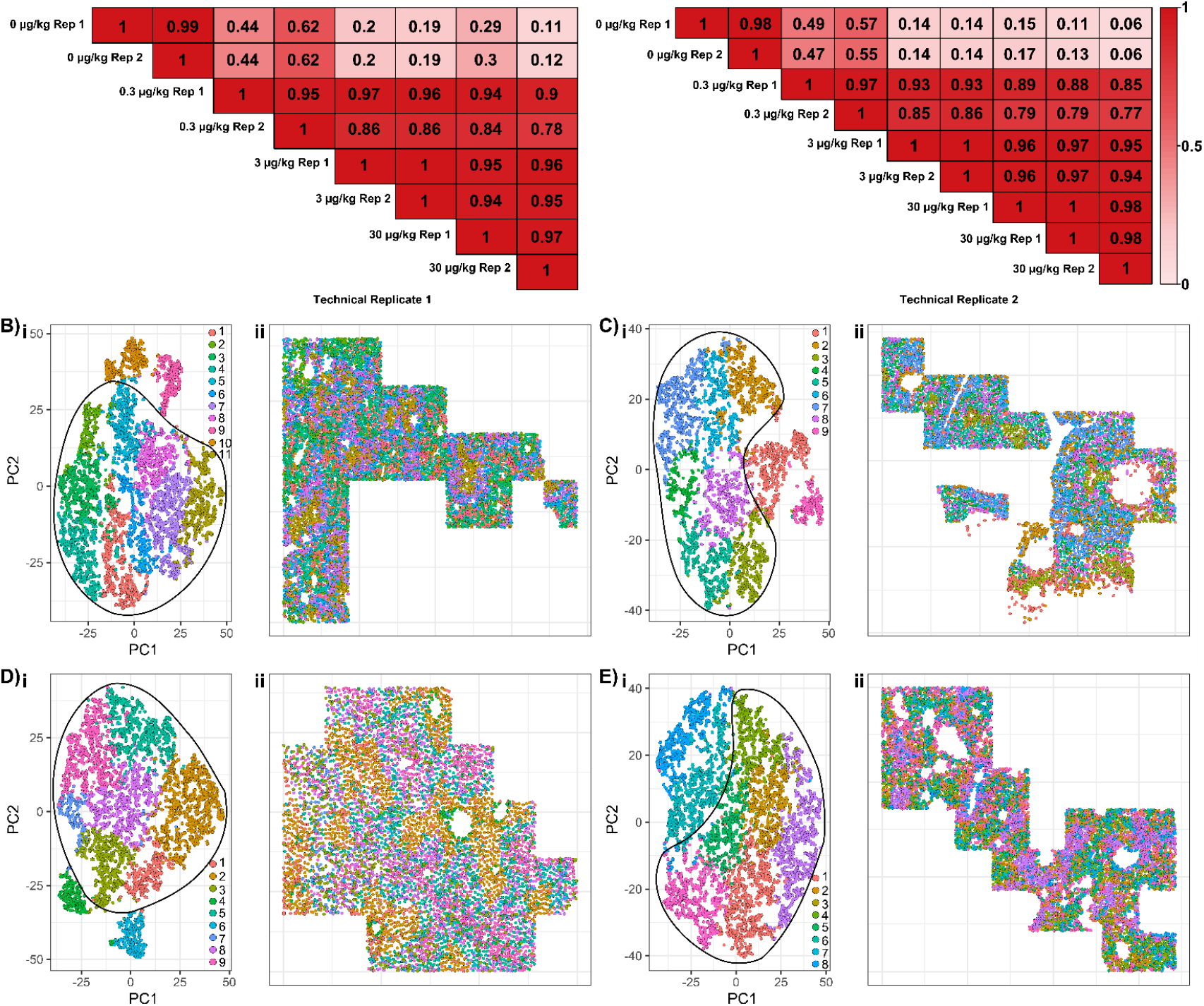
Biological and technical replicate analysis of spatial transcriptomic using Baysor and MERINGUE. (A) Pearson correlation of transcript counts for regions of interest (ROI) comparing biological replication across two technical replicates. ROIs were processed using Baysor v0.5.0 for cell segmentation based on DAPI staining and transcript abundances (Petukhov *et al.*, 2021). Co-expression and, dimensionality reduction, and clustering was performed using MERINGUE (Miller *et al.*, 2021). Representative PCA plots (subplots **i**) and color-coded ROIs for liver sections (subplots **ii**) from mice gavaged every 4 days for 28 days with (B) sesame oil vehicle control, (C) 0.3 μg/kg, (D) 3 μg/kg, or (E) 30 μg/kg TCDD. Each tissue section was processed independently (cluster numbers are not equivalent for each sample) and numbers associated with each color reflect an identified cluster but was not assigned a cell type. Examination of genes associated with individual clusters for each slide were used to draw circles around cells representing hepatocytes (inside circle) or macrophage and stellate cells (outside circle). Mapped ROIs show location of cells colored by cluster. Images are not equally scaled due to differences in ROI dimensions.

**Fig S8.**
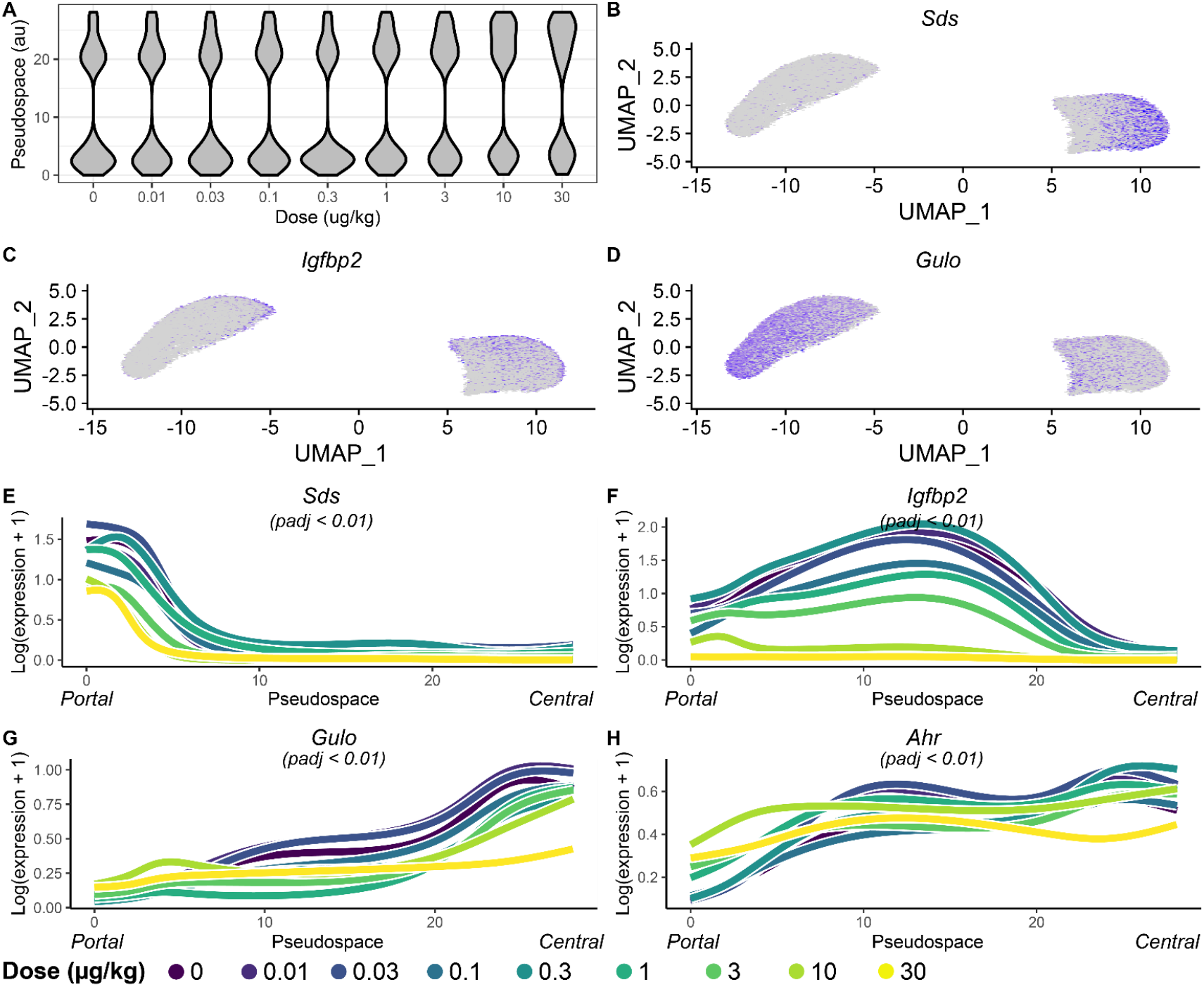
Distribution of hepatocytes following pseudospace trajectory analysis. (A) Violin plots show the distribution of hepatocytes along the pseudospace trajectory at each TCDD dose. UMAP visualization shows the hepatocyte expression of (B) portal marker *Sds*, (C) midzonal marker *Igfbp2*, and (D) central marker *Gulo.* Dosedependent expression along the pseudospace continuum is shown for (E) *Sds*, (F) *Igfbp2*, (G) *Gulo*, and (H) *Ahr.* Adjusted p-values were calculated based on a Wald statistic for the TCDD treatment effect.

**Fig S9.**
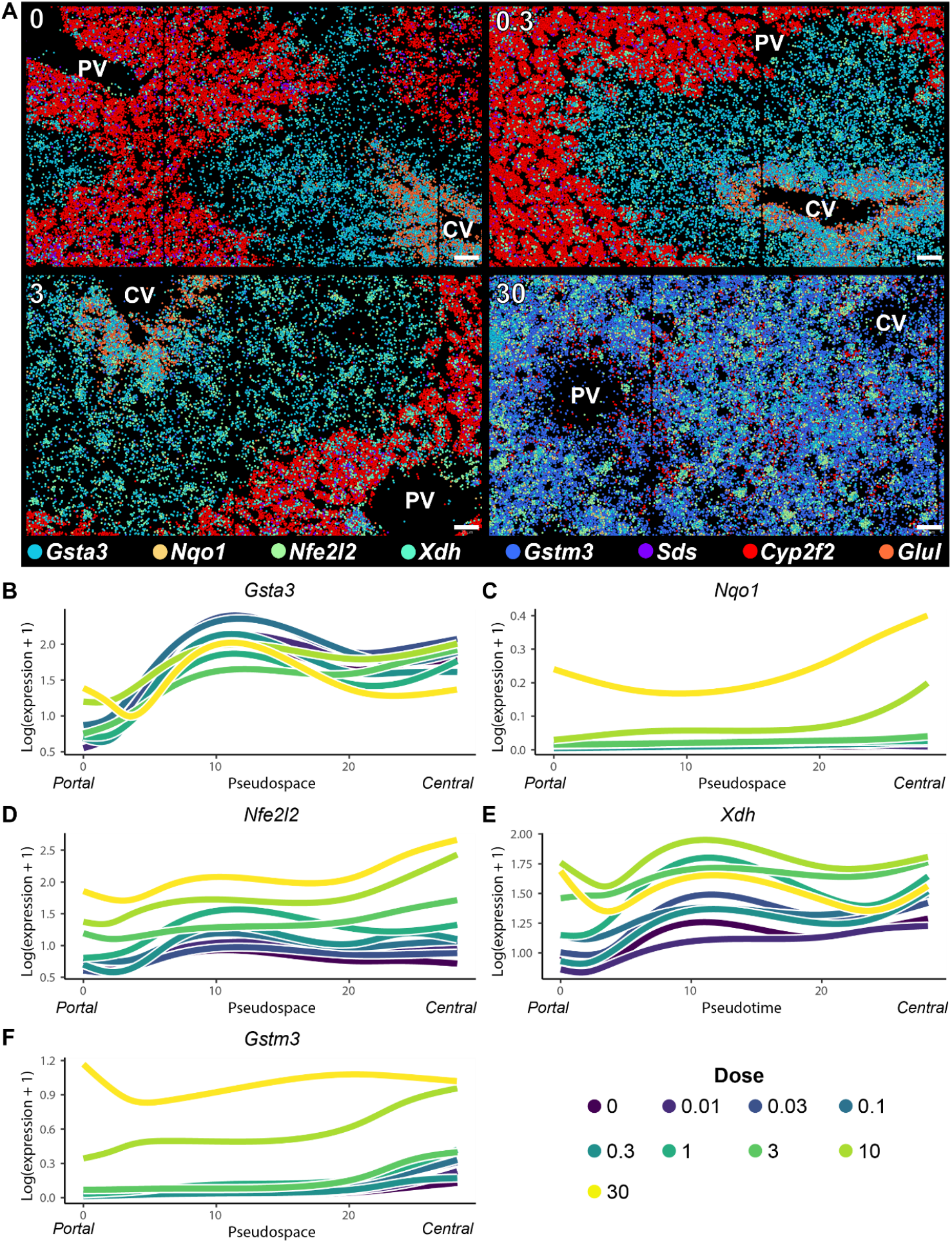
Spatial and pseudospatial expression of oxidative stress genes in liver sections from male mice gavaged with TCDD every 4 days for 28 days. (A) Molecular cartography was used to visualize the spatial distribution of *Gsta3, Nqo1, Nfe2l2, Xdh*, and *Gstm3* oxidative stress related genes along with the portal markers *Sds* and *Cyp2f2*, and the central marker *Glul*. Colors represent individual genes while individual spots reflect a single transcript. Genes are ordered from topmost layer to bottom layer for overlapping points. Dose-dependent expression along the pseudospace continuum is shown for (B) *Gsta2*, (C) *Nqo1*, (D) *Nfe2l2*, (E) *Xdh*, and (F) *Gstm3*.

**Fig S10.**
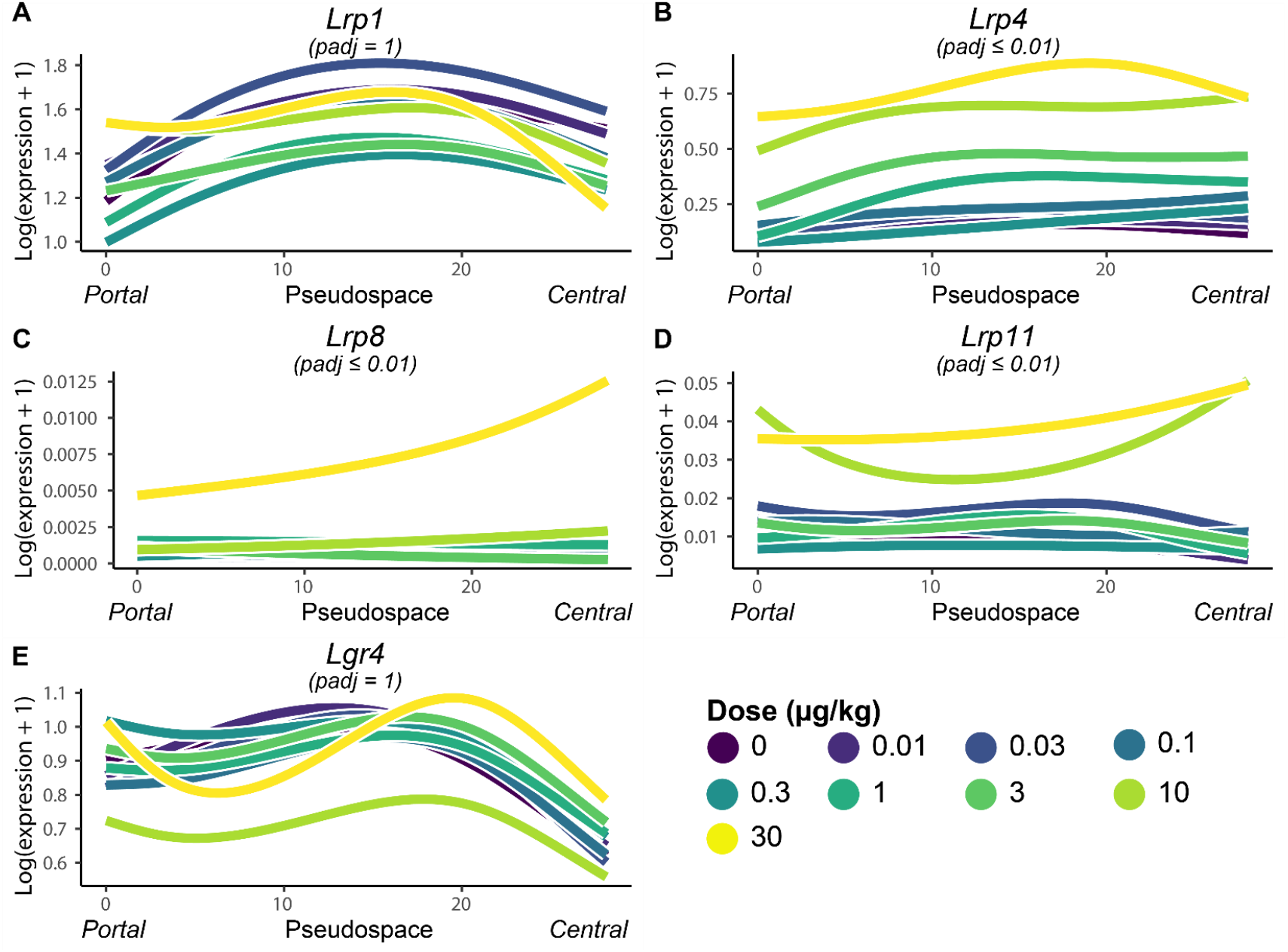
Dose dependent effects of TCDD on the pseudospatial expression of Wnt/β-catenin signaling related co-receptors. Dose-dependent expression along the pseudospace continuum is shown for (A) *Lrp1*, (B) *Lrp4*, (C) *Lrp8*, (D) *Lrp11*, and (E) *Lgr4.* Adjusted p-values were calculated based on a Wald statistic for an effect of TCDD treatment.

**Fig S11.**
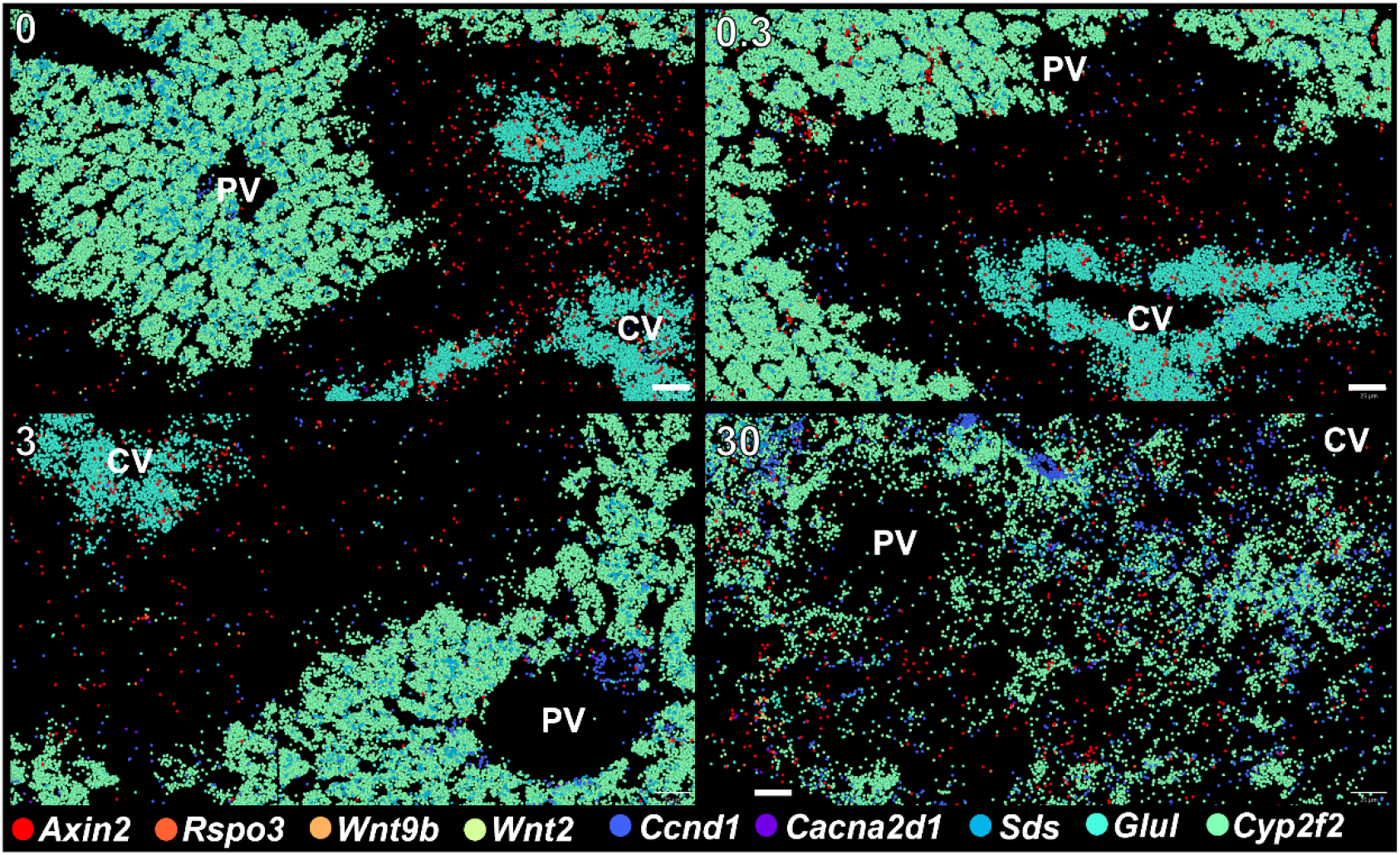
Spatial expression of Wnt/β-catenin signaling related genes in livers of male mice gavaged with TCDD every 4 days for 28 days. (A) Molecular cartography was used to visualize the spatial distribution of Wnt/β-catenin signaling related genes *Axin2, Rspo3, Wnt9b, Wnt2, and Ccnd1* along with portal markers *Sds* and *Cyp2f2*, and central marker *Glul*. Colors represent individual genes and individual spots reflect a single transcript molecule and genes are listed from topmost layer to bottom layer for overlapping points. CV indicated central vein, PV indicates portal vein.

**Fig S12.**
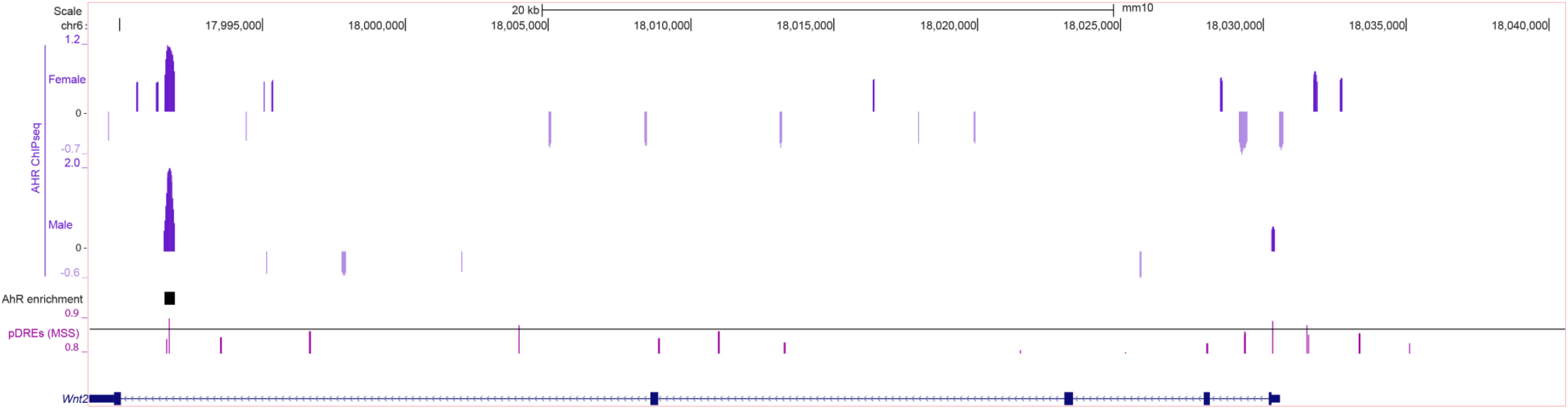
Presence of AHR binding and putative DREs (pDREs) in *Wnt2* genomic region. Chromatin immunoprecipitation (ChIP) sequencing data for AHR from mice treated with TCDD for 2hrs was obtained from GEO (GSE109865, GSE97636). Presence of pDREs determined as previously described is shown with (Nault *et al.*, 2016) a matrix similarity score (MSS) of 0.874.

## LITERATURE CITED

Andersen, M. E., Birnbaum, L. S., Barton, H. A. and Eklund, C. R. (1997). Regional hepatic CYP1A1 and CYP1A2 induction with 2,3,7,8-tetrachlorodibenzo-p-dioxin evaluated with a multicompartment geometric model of hepatic zonation. Toxicol Appl Pharmacol 144(1), 145–55.

Angrish, M. M., Mets, B. D., Jones, A. D. and Zacharewski, T. R. (2012). Dietary fat is a lipid source in 2,3,7,8-tetrachlorodibenzo-p-dioxin (TCDD)-elicited hepatic steatosis in C57BL/6 mice. Toxicol Sci 128(2), 377–86. 3493189

Arai, T., Ono, Y., Arimura, Y., Sayama, K., Suzuki, T., Shinjo, S., Kanai, M., Abe, S.-i., Semba, K. and Goda, N. (2017). Type I neuregulin1α is a novel local mediator to suppress hepatic gluconeogenesis in mice. Scientific reports 7(1), 42959.

Babu, R. O., Lui, V. C. H., Chen, Y., Yiu, R. S. W., Ye, Y., Niu, B., Wu, Z., Zhang, R., Yu, M. O. N., Chung, P. H. Y., Wong, K. K. Y., Xia, H., Zhang, M. Q., Wang, B., Lendahl, U. and Tam, P. K. H. (2020). Beta-amyloid deposition around hepatic bile ducts is a novel pathobiological and diagnostic feature of biliary atresia. Journal of hepatology 73(6), 1391–1403.

Bahar Halpern, K., Caspi, I., Lemze, D., Levy, M., Landen, S., Elinav, E., Ulitsky, I. and Itzkovitz, S. (2015). Nuclear Retention of mRNA in Mammalian Tissues. Cell reports 13(12), 2653–62. 4700052

Behari, J., Yeh, T. H., Krauland, L., Otruba, W., Cieply, B., Hauth, B., Apte, U., Wu, T., Evans, R. and Monga, S. P. (2010). Liver-specific beta-catenin knockout mice exhibit defective bile acid and cholesterol homeostasis and increased susceptibility to diet-induced steatohepatitis. The American journal of pathology 176(2), 744–53. PMC2808081

Benhamouche, S., Decaens, T., Godard, C., Chambrey, R., Rickman, D. S., Moinard, C., Vasseur-Cognet, M., Kuo, C. J., Kahn, A., Perret, C. and Colnot, S. (2006). Apc tumor suppressor gene is the “zonation-keeper” of mouse liver. Developmental cell 10(6), 759–70.

Bieghs, V., Verheyen, F., van Gorp, P. J., Hendrikx, T., Wouters, K., Lutjohann, D., Gijbels, M. J., Febbraio, M., Binder, C. J., Hofker, M. H. and Shiri-Sverdlov, R. (2012). Internalization of modified lipids by CD36 and SR-A leads to hepatic inflammation and lysosomal cholesterol storage in Kupffer cells. PloS one 7(3), e34378. PMC3314620

Bleriot, C. and Ginhoux, F. (2019). Understanding the Heterogeneity of Resident Liver Macrophages. Frontiers in immunology 10, 2694. PMC6877662

Bonnardel, J., T’Jonck, W., Gaublomme, D., Browaeys, R., Scott, C. L., Martens, L., Vanneste, B., De Prijck, S., Nedospasov, S. A., Kremer, A., Van Hamme, E., Borghgraef, P., Toussaint, W., De Bleser, P., Mannaerts, I., Beschin, A., van Grunsven, L. A., Lambrecht, B. N., Taghon, T., Lippens, S., Elewaut, D., Saeys, Y. and Guilliams, M. (2019). Stellate Cells, Hepatocytes, and Endothelial Cells Imprint the Kupffer Cell Identity on Monocytes Colonizing the Liver Macrophage Niche. Immunity 51(4), 638–654 e9. PMC6876284

Boverhof, D. R., Burgoon, L. D., Tashiro, C., Chittim, B., Harkema, J. R., Jump, D. B. and Zacharewski, T. R. (2005). Temporal and dose-dependent hepatic gene expression patterns in mice provide new insights into TCDD-Mediated hepatotoxicity. Toxicol Sci 85(2), 1048–63.

Braeuning, A., Kohle, C., Buchmann, A. and Schwarz, M. (2011). Coordinate regulation of cytochrome P450 1a1 expression in mouse liver by the aryl hydrocarbon receptor and the beta-catenin pathway. Toxicol Sci 122(1), 16–25.

Buniatian, G. H., Weiskirchen, R., Weiss, T. S., Schwinghammer, U., Fritz, M., Seferyan, T., Proksch, B., Glaser, M., Lourhmati, A., Buadze, M., Borkham-Kamphorst, E., Gaunitz, F., Gleiter, C. H., Lang, T., Schaeffeler, E., Tremmel, R., Cynis, H., Frey, W. H., 2nd, Gebhardt, R., Friedman, S. L., Mikulits, W., Schwab, M. and Danielyan, L. (2020). Antifibrotic Effects of Amyloid-Beta and Its Loss in Cirrhotic Liver. Cells 9(2). PMC7072823

Burke, Z. D., Reed, K. R., Phesse, T. J., Sansom, O. J., Clarke, A. R. and Tosh, D. (2009). Liver zonation occurs through a beta-catenin-dependent, c-Myc-independent mechanism. Gastroenterology 136(7), 2316–2324 e1-3.

Cardoso, C. C., Matiollo, C., Pereira, C. H. J., Fonseca, J. S., Alves, H. E. L., da Silva, O. M., de Souza Menegassi, V., Dos Santos, C. R., de Moraes, A. C. R., de Lucca Schiavon, L. and Santos-Silva, M. C. (2021). Patterns of dendritic cell and monocyte subsets are associated with disease severity and mortality in liver cirrhosis patients. Scientific reports 11(1), 5923. PMC7960697

Carter-Kent, C., Brunt, E. M., Yerian, L. M., Alkhouri, N., Angulo, P., Kohli, R., Ling, S. C., Xanthakos, S. A., Whitington, P. F., Charatcharoenwitthaya, P., Yap, J., Lopez, R., McCullough, A. J. and Feldstein, A. E. (2011). Relations of steatosis type, grade, and zonality to histological features in pediatric nonalcoholic fatty liver disease. J Pediatr Gastroenterol Nutr 52(2), 190–7.

Cave, M., Appana, S., Patel, M., Falkner, K. C., McClain, C. J. and Brock, G. (2010). Polychlorinated biphenyls, lead, and mercury are associated with liver disease in American adults: NHANES 2003-2004. Environmental health perspectives 118(12), 1735–42. 3002193

Cholico, G. N., Fling, R. R., Zacharewski, N. A., Fader, K. A., Nault, R. and Zacharewski, T. R. (2021). Thioesterase induction by 2,3,7,8-tetrachlorodibenzo-p-dioxin results in a futile cycle that inhibits hepatic beta-oxidation. Scientific reports 11(1), 15689. PMC8333094

Cholico, G. N., Nault, R. and Zacharewski, T. R. (2022). Genome-Wide ChIPseq Analysis of AhR, COUP-TF, and HNF4 Enrichment in TCDD-Treated Mouse Liver. International journal of molecular sciences 23(3). PMC8836158

Chrostek, L. and Panasiuk, A. (2014). Liver fibrosis markers in alcoholic liver disease. World journal of gastroenterology 20(25), 8018–23. PMC4081671

Crespo, M., Gonzalez-Teran, B., Nikolic, I., Mora, A., Folgueira, C., Rodriguez, E., Leiva-Vega, L., Pintor-Chocano, A., Fernandez-Chacon, M., Ruiz-Garrido, I., Cicuendez, B., Tomas-Loba, A., N, A. G., Caballero-Molano, A., Beiroa, D., Hernandez-Cosido, L., Torres, J. L., Kennedy, N. J., Davis, R. J., Benedito, R., Marcos, M., Nogueiras, R., Hidalgo, A., Matesanz, N., Leiva, M. and Sabio, G. (2020). Neutrophil infiltration regulates clock-gene expression to organize daily hepatic metabolism. eLife 9. PMC7723411

Cunningham, R. P. and Porat-Shliom, N. (2021). Liver Zonation - Revisiting Old Questions With New Technologies. Frontiers in physiology 12, 732929. PMC8458816

D’Gama, P. P., Qiu, T., Cosacak, M. I., Rayamajhi, D., Konac, A., Hansen, J. N., Ringers, C., Acuna-Hinrichsen, F., Hui, S. P., Olstad, E. W., Chong, Y. L., Lim, C. K. A., Gupta, A., Ng, C. P., Nilges, B. S., Kashikar, N. D., Wachten, D., Liebl, D., Kikuchi, K., Kizil, C., Yaksi, E., Roy, S. and Jurisch-Yaksi, N. (2021). Diversity and function of motile ciliated cell types within ependymal lineages of the zebrafish brain. Cell reports 37(1), 109775. PMC8524669

Datlinger, P., Rendeiro, A. F., Boenke, T., Senekowitsch, M., Krausgruber, T., Barreca, D. and Bock, C. (2021). Ultra-high-throughput single-cell RNA sequencing and perturbation screening with combinatorial fluidic indexing. Nature methods.

Dobie, R., Wilson-Kanamori, J. R., Henderson, B. E. P., Smith, J. R., Matchett, K. P., Portman, J. R., Wallenborg, K., Picelli, S., Zagorska, A., Pendem, S. V., Hudson, T. E., Wu, M. M., Budas, G. R., Breckenridge, D. G., Harrison, E. M., Mole, D. J., Wigmore, S. J., Ramachandran, P., Ponting, C. P., Teichmann, S. A., Marioni, J. C. and Henderson, N. C. (2019). Single-Cell Transcriptomics Uncovers Zonation of Function in the Mesenchyme during Liver Fibrosis. Cell reports 29(7), 1832–1847 e8. 6856722

Dornbos, P., Jurgelewicz, A., Fader, K. A., Williams, K., Zacharewski, T. R. and LaPres, J. J. (2019). Characterizing the Role of HMG-CoA Reductase in Aryl Hydrocarbon Receptor-Mediated Liver Injury in C57BL/6 Mice. Scientific reports 9(1), 15828. 6825130

Doskey, C. M., Fader, K. A., Nault, R., Lydic, T., Matthews, J., Potter, D., Sharratt, B., Williams, K. and Zacharewski, T. (2020). 2,3,7,8-Tetrachlorodibenzo-p-dioxin (TCDD) alters hepatic polyunsaturated fatty acid metabolism and eicosanoid biosynthesis in female Sprague-Dawley rats. Toxicol Appl Pharmacol 398, 115034. 7294678

Efremova, M., Vento-Tormo, M., Teichmann, S. A. and Vento-Tormo, R. (2020). CellPhoneDB: inferring cell-cell communication from combined expression of multi-subunit ligand-receptor complexes. Nature protocols 15(4), 1484–1506.

Fader, K. A., Nault, R., Doskey, C. M., Fling, R. R. and Zacharewski, T. R. (2019). 2,3,7,8-Tetrachlorodibenzo-p-dioxin abolishes circadian regulation of hepatic metabolic activity in mice. Scientific reports 9(1), 6514. 6478849

Fader, K. A., Nault, R., Kirby, M. P., Markous, G., Matthews, J. and Zacharewski, T. R. (2017a). Convergence of hepcidin deficiency, systemic iron overloading, heme accumulation, and REV-ERBalpha/beta activation in aryl hydrocarbon receptor-elicited hepatotoxicity. Toxicol Appl Pharmacol 321, 1–17. 5421516

Fader, K. A., Nault, R., Zhang, C., Kumagai, K., Harkema, J. R. and Zacharewski, T. R. (2017b). 2,3,7,8-Tetrachlorodibenzo-p-dioxin (TCDD)-elicited effects on bile acid homeostasis: Alterations in biosynthesis, enterohepatic circulation, and microbial metabolism. Scientific reports 7(1), 5921. 5517430

Flach, R. J., Qin, H., Zhang, L. and Bennett, A. M. (2011). Loss of mitogen-activated protein kinase phosphatase-1 protects from hepatic steatosis by repression of cell death-inducing DNA fragmentation factor A (DFFA)-like effector C (CIDEC)/fat-specific protein 27. J Biol Chem 286(25), 22195–202. PMC3121364

Gebhardt, R. and Hovhannisyan, A. (2010). Organ patterning in the adult stage: the role of Wnt/beta-catenin signaling in liver zonation and beyond. Dev Dyn 239(1), 45–55.

Gerbal-Chaloin, S., Dume, A. S., Briolotti, P., Klieber, S., Raulet, E., Duret, C., Fabre, J. M., Ramos, J., Maurel, P. and Daujat-Chavanieu, M. (2014). The WNT/beta-catenin pathway is a transcriptional regulator of CYP2E1, CYP1A2, and aryl hydrocarbon receptor gene expression in primary human hepatocytes. Molecular pharmacology 86(6), 624–34.

Ghallab, A., Myllys, M., Friebel, A., Duda, J., Edlund, K., Halilbasic, E., Vucur, M., Hobloss, Z., Brackhagen, L., Begher-Tibbe, B., Hassan, R., Burke, M., Genc, E., Frohwein, L. J., Hofmann, U., Holland, C. H., Gonzalez, D., Keller, M., Seddek, A. L., Abbas, T., Mohammed, E. S. I., Teufel, A., Itzel, T., Metzler, S., Marchan, R., Cadenas, C., Watzl, C., Nitsche, M. A., Kappenberg, F., Luedde, T., Longerich, T., Rahnenfuhrer, J., Hoehme, S., Trauner, M. and Hengstler, J. G. (2021). Spatio-Temporal Multiscale Analysis of Western Diet-Fed Mice Reveals a Translationally Relevant Sequence of Events during NAFLD Progression. Cells 10(10). PMC8533774

Groiss, S., Pabst, D., Faber, C., Meier, A., Bogdoll, A., Unger, C., Nilges, B., Strauss, S., Föderl-Höbenreich, E., Hardt, M., Geipel, A., Reinecke, F., Korfhage, C. and Zatloukal, K. (2021). Highly resolved spatial transcriptomics for detection of rare events in cells. bioRxiv, 2021.10.11.463936.

Guilliams, M., Bonnardel, J., Haest, B., Vanderborght, B., Wagner, C., Remmerie, A., Bujko, A., Martens, L., Thone, T., Browaeys, R., De Ponti, F. F., Vanneste, B., Zwicker, C., Svedberg, F. R., Vanhalewyn, T., Goncalves, A., Lippens, S., Devriendt, B., Cox, E., Ferrero, G., Wittamer, V., Willaert, A., Kaptein, S. J. F., Neyts, J., Dallmeier, K., Geldhof, P., Casaert, S., Deplancke, B., Ten Dijke, P., Hoorens, A., Vanlander, A., Berrevoet, F., Van Nieuwenhove, Y., Saeys, Y., Saelens, W., Van Vlierberghe, H., Devisscher, L. and Scott, C. L. (2022). Spatial proteogenomics reveals distinct and evolutionarily conserved hepatic macrophage niches. Cell 185(2), 379–396 e38.

Hall, Z., Bond, N. J., Ashmore, T., Sanders, F., Ament, Z., Wang, X., Murray, A. J., Bellafante, E., Virtue, S., Vidal-Puig, A., Allison, M., Davies, S. E., Koulman, A., Vacca, M. and Griffin, J. L. (2017). Lipid zonation and phospholipid remodeling in nonalcoholic fatty liver disease. Hepatology 65(4), 1165–1180. PMC5396354

Halpern, K. B., Shenhav, R., Massalha, H., Toth, B., Egozi, A., Massasa, E. E., Medgalia, C., David, E., Giladi, A., Moor, A. E., Porat, Z., Amit, I. and Itzkovitz, S. (2018). Paired-cell sequencing enables spatial gene expression mapping of liver endothelial cells. Nature biotechnology.

Halpern, K. B., Shenhav, R., Matcovitch-Natan, O., Toth, B., Lemze, D., Golan, M., Massasa, E. E., Baydatch, S., Landen, S., Moor, A. E., Brandis, A., Giladi, A., Avihail, A. S., David, E., Amit, I. and Itzkovitz, S. (2017). Single-cell spatial reconstruction reveals global division of labour in the mammalian liver. Nature 542(7641), 352–356. 5321580

Haskins, J. W., Zhang, S., Means, R. E., Kelleher, J. K., Cline, G. W., Canfran-Duque, A., Suarez, Y. and Stern, D. F. (2015). Neuregulin-activated ERBB4 induces the SREBP-2 cholesterol biosynthetic pathway and increases low-density lipoprotein uptake. Sci Signal 8(401), ra111. PMC4666504

Hijmans, B. S., Grefhorst, A., Oosterveer, M. H. and Groen, A. K. (2014). Zonation of glucose and fatty acid metabolism in the liver: mechanism and metabolic consequences. Biochimie 96, 121–9.

Hu, S., Liu, S., Bian, Y., Poddar, M., Singh, S., Cao, C., McGaughey, J., Bell, A., Blazer, L. L., Adams, J. J., Sidhu, S. S., Angers, S. and Monga, S. P. (2022). Dynamic control of metabolic zonation and liver repair by endothelial cell Wnt2 and Wnt9b revealed by single cell spatial transcriptomics using Molecular Cartography. bioRxiv, 2022.03.18.484868.

Jackson, D. P., Li, H., Mitchell, K. A., Joshi, A. D. and Elferink, C. J. (2014). Ah receptor-mediated suppression of liver regeneration through NC-XRE-driven p21Cip1 expression. Molecular pharmacology 85(4), 533–41. 3965890

Jungermann, K. (1995). Zonation of metabolism and gene expression in liver. Histochem Cell Biol 103(2), 81–91.

Karin, D., Koyama, Y., Brenner, D. and Kisseleva, T. (2016). The characteristics of activated portal fibroblasts/myofibroblasts in liver fibrosis. Differentiation 92(3), 84–92. PMC5079826

Kietzmann, T. (2019). Liver Zonation in Health and Disease: Hypoxia and Hypoxia-Inducible Transcription Factors as Concert Masters. International journal of molecular sciences 20(9). PMC6540308

Koyama, Y., Wang, P., Liang, S., Iwaisako, K., Liu, X., Xu, J., Zhang, M., Sun, M., Cong, M., Karin, D., Taura, K., Benner, C., Heinz, S., Bera, T., Brenner, D. A. and Kisseleva, T. (2017). Mesothelin/mucin 16 signaling in activated portal fibroblasts regulates cholestatic liver fibrosis. The Journal of clinical investigation 127(4), 1254–1270. PMC5373891

Lee, D. H., Lee, I. K., Song, K., Steffes, M., Toscano, W., Baker, B. A. and Jacobs, D. R., Jr. (2006). A strong dose-response relation between serum concentrations of persistent organic pollutants and diabetes: results from the National Health and Examination Survey 1999-2002. Diabetes care 29(7), 1638–44.

Lee, J. H., Wada, T., Febbraio, M., He, J., Matsubara, T., Lee, M. J., Gonzalez, F. J. and Xie, W. (2010). A novel role for the dioxin receptor in fatty acid metabolism and hepatic steatosis. Gastroenterology 139(2), 653–63. 2910786

Li, C., Liu, Y., Dong, Z., Xu, M., Gao, M., Cong, M. and Liu, S. (2020). TCDD promotes liver fibrosis through disordering systemic and hepatic iron homeostasis. J Hazard Mater 395, 122588.

Li, Z., Dranoff, J. A., Chan, E. P., Uemura, M., Sevigny, J. and Wells, R. G. (2007). Transforming growth factor-beta and substrate stiffness regulate portal fibroblast activation in culture. Hepatology 46(4), 1246–56.

Massalha, H., Bahar Halpern, K., Abu-Gazala, S., Jana, T., Massasa, E. E., Moor, A. E., Buchauer, L., Rozenberg, M., Pikarsky, E., Amit, I., Zamir, G. and Itzkovitz, S. (2020). A single cell atlas of the human liver tumor microenvironment. Molecular systems biology 16(12), e9682. PMC7746227

Matsuda, S., Matsuda, Y. and D’Adamio, L. (2009). CD74 interacts with APP and suppresses the production of Abeta. Mol Neurodegener 4, 41. PMC2770512

McGinnis, C. S., Patterson, D. M., Winkler, J., Conrad, D. N., Hein, M. Y., Srivastava, V., Hu, J. L., Murrow, L. M., Weissman, J. S., Werb, Z., Chow, E. D. and Gartner, Z. J. (2019). MULTI-seq: sample multiplexing for single-cell RNA sequencing using lipid-tagged indices. Nature methods 16(7), 619–626. PMC6837808

Miller, B. F., Bambah-Mukku, D., Dulac, C., Zhuang, X. and Fan, J. (2021). Characterizing spatial gene expression heterogeneity in spatially resolved single-cell transcriptomic data with nonuniform cellular densities. Genome research 31(10), 1843–1855. PMC8494224

Miura, K., Yang, L., van Rooijen, N., Ohnishi, H. and Seki, E. (2012). Hepatic recruitment of macrophages promotes nonalcoholic steatohepatitis through CCR2. American journal of physiology. Gastrointestinal and liver physiology 302(11), G1310–21. PMC3378163

Moreno-Marin, N., Merino, J. M., Alvarez-Barrientos, A., Patel, D. P., Takahashi, S., Gonzalez-Sancho, J. M., Gandolfo, P., Rios, R. M., Munoz, A., Gonzalez, F. J. and Fernandez-Salguero, P. M. (2018). Aryl Hydrocarbon Receptor Promotes Liver Polyploidization and Inhibits PI3K, ERK, and Wnt/beta-Catenin Signaling. iScience 4, 44–63. 6147018

Nauli, A. M. and Matin, S. (2019). Why Do Men Accumulate Abdominal Visceral Fat? Frontiers in physiology 10.

Nault, R., Colbry, D., Brandenberger, C., Harkema, J. R. and Zacharewski, T. R. (2015). Development of a computational high-throughput tool for the quantitative examination of dose-dependent histological features. Toxicologic pathology 43(3), 366–75. 4382446

Nault, R., Fader, K. A., Bhattacharya, S. and Zacharewski, T. R. (2021). Single-Nuclei RNA Sequencing Assessment of the Hepatic Effects of 2,3,7,8-Tetrachlorodibenzo-p-dioxin. Cell Mol Gastroenterol Hepatol 11(1), 147–159. PMC7674514

Nault, R., Fader, K. A., Harkema, J. R. and Zacharewski, T. (2017a). Loss of liver-specific and sexually dimorphic gene expression by aryl hydrocarbon receptor activation in C57BL/6 mice. PloS one 12(9), e0184842. PMC5602546

Nault, R., Fader, K. A., Kopec, A. K., Harkema, J. R., Zacharewski, T. R. and Luyendyk, J. P. (2016). From the Cover: Coagulation-Driven Hepatic Fibrosis Requires Protease Activated Receptor-1 (PAR-1) in a Mouse Model of TCDD-Elicited Steatohepatitis. Toxicol Sci 154(2), 381–391. 5139072

Nault, R., Fader, K. A., Lydic, T. A. and Zacharewski, T. R. (2017b). Lipidomic Evaluation of Aryl Hydrocarbon Receptor-Mediated Hepatic Steatosis in Male and Female Mice Elicited by 2,3,7,8-Tetrachlorodibenzo-p-dioxin. Chemical research in toxicology 30(4), 1060–1075.

Nault, R., Saha, S., Bhattacharya, S., Dodson, J., Sinha, S., Maiti, T. and Zacharewski, T. (2022). Benchmarking of a Bayesian single cell RNAseq differential gene expression test for dose-response study designs. Nucleic acids research.

Ou, R., Liu, J., Lv, M., Wang, J., Wang, J., Zhu, L., Zhao, L. and Xu, Y. (2017). Neutrophil depletion improves diet-induced non-alcoholic fatty liver disease in mice. Endocrine 57(1), 72–82.

Panday, R., Monckton, C. P. and Khetani, S. R. (2022). The Role of Liver Zonation in Physiology, Regeneration, and Disease. Seminars in liver disease 42(1), 1–16.

Park, S., Wu, L., Tu, J., Yu, W., Toh, Y., Carmon, K. S. and Liu, Q. J. (2020). Unlike LGR4, LGR5 potentiates Wnt-beta-catenin signaling without sequestering E3 ligases. Sci Signal 13(660). PMC7905944

Pelclova, D., Urban, P., Preiss, J., Lukas, E., Fenclova, Z., Navratil, T., Dubska, Z. and Senholdova, Z. (2006). Adverse health effects in humans exposed to 2,3,7,8-tetrachlorodibenzo-p-dioxin (TCDD). Reviews on environmental health 21 (2), 119–38.

Percie du Sert, N., Hurst, V., Ahluwalia, A., Alam, S., Avey, M. T., Baker, M., Browne, W. J., Clark, A., Cuthill, I. C., Dirnagl, U., Emerson, M., Garner, P., Holgate, S. T., Howells, D. W., Karp, N. A., Lazic, S. E., Lidster, K., MacCallum, C. J., Macleod, M., Pearl, E. J., Petersen, O. H., Rawle, F., Reynolds, P., Rooney, K., Sena, E. S., Silberberg, S. D., Steckler, T. and Wurbel, H. (2020). The ARRIVE guidelines 2.0: Updated guidelines for reporting animal research. BMC veterinary research 16(1), 242. 7359286

Petukhov, V., Xu, R. J., Soldatov, R. A., Cadinu, P., Khodosevich, K., Moffitt, J. R. and Kharchenko, P. V. (2021). Cell segmentation in imaging-based spatial transcriptomics. Nature biotechnology.

Prochazkova, J., Kabatkova, M., Bryja, V., Umannova, L., Bernatik, O., Kozubik, A., Machala, M. and Vondracek, J. (2011). The interplay of the aryl hydrocarbon receptor and beta-catenin alters both AhR-dependent transcription and Wnt/beta-catenin signaling in liver progenitors. Toxicol Sci 122(2), 349–60.

Ratziu, V., Massard, J., Charlotte, F., Messous, D., Imbert-Bismut, F., Bonyhay, L., Tahiri, M., Munteanu, M., Thabut, D., Cadranel, J. F., Le Bail, B., de Ledinghen, V., Poynard, T., Group, L. S. and group, C. s. (2006). Diagnostic value of biochemical markers (FibroTest-FibroSURE) for the prediction of liver fibrosis in patients with non-alcoholic fatty liver disease. BMC Gastroenterol 6, 6. PMC1386692

Righelli, D., Weber, L. M., Crowell, H. L., Pardo, B., Collado-Torres, L., Ghazanfar, S., Lun, A. T. L., Hicks, S. C. and Risso, D. (2022). SpatialExperiment: infrastructure for spatially resolved transcriptomics data in R using Bioconductor. Bioinformatics.

Rocha, A. S., Vidal, V., Mertz, M., Kendall, T. J., Charlet, A., Okamoto, H. and Schedl, A. (2015). The Angiocrine Factor Rspondin3 Is a Key Determinant of Liver Zonation. Cell reports 13(9), 1757–64.

Saponara, E., Penno, C., Matadamas Guzmán, M. L., Brun, V., Fischer, B., Brousseau, M., Wang, Z.-Y., ODonnell, P., Turner, J., Meyer, A. G., Bollepalli, L., d’Ario, G., Roma, G., Carbone, W., Orsini, V., Annunziato, S., Obrecht, M., Beckmann, N., Saravanan, C., Osmont, A., Tropberger, P., Richards, S., Genoud, C., Aebi, A., Ley, S., Ksiazek, I., Nigsch, F., Terraciano, L., Bouwmeester, T., Tchorz, J. and Ruffner, H. (2021). Loss of hepatic Lgr4 and Lgr5 promotes nonalcoholic fatty liver disease. bioRxiv, 2021.11.22.469602.

Schleicher, J., Dahmen, U., Guthke, R. and Schuster, S. (2017). Zonation of hepatic fat accumulation: insights from mathematical modelling of nutrient gradients and fatty acid uptake. J R Soc Interface 14(133). PMC5582132

Song, Z., Chen, W., Athavale, D., Ge, X., Desert, R., Das, S., Han, H. and Nieto, N. (2021). Osteopontin Takes Center Stage in Chronic Liver Disease. Hepatology 73(4), 1594–1608. PMC8106357

Soto-Gutierrez, A., Gough, A., Vernetti, L. A., Taylor, D. L. and Monga, S. P. (2017). Pre-clinical and clinical investigations of metabolic zonation in liver diseases: The potential of microphysiology systems. Exp Biol Med (Maywood) 242(16), 1605–1616. PMC5661767

Street, K., Risso, D., Fletcher, R. B., Das, D., Ngai, J., Yosef, N., Purdom, E. and Dudoit, S. (2018). Slingshot: cell lineage and pseudotime inference for single-cell transcriptomics. BMC genomics 19(1), 477. 6007078

Takeda, T., Komiya, Y., Koga, T., Ishida, T., Ishii, Y., Kikuta, Y., Nakaya, M., Kurose, H., Yokomizo, T., Shimizu, T., Uchi, H., Furue, M. and Yamada, H. (2017). Dioxin-induced increase in leukotriene B4 biosynthesis through the aryl hydrocarbon receptor and its relevance to hepatotoxicity owing to neutrophil infiltration. J Biol Chem.

Taylor, K. W., Novak, R. F., Anderson, H. A., Birnbaum, L. S., Blystone, C., Devito, M., Jacobs, D., Kohrle, J., Lee, D. H., Rylander, L., Rignell-Hydbom, A., Tornero-Velez, R., Turyk, M. E., Boyles, A. L., Thayer, K. A. and Lind, L. (2013). Evaluation of the association between persistent organic pollutants (POPs) and diabetes in epidemiological studies: a national toxicology program workshop review. Environmental health perspectives 121 (7), 774–83. 3701910

Vondracek, J. and Machala, M. (2016). Environmental Ligands of the Aryl Hydrocarbon Receptor and Their Effects in Models of Adult Liver Progenitor Cells. Stem Cells Int 2016, 4326194. 4870370

Wahlang, B., Jin, J., Beier, J. I., Hardesty, J. E., Daly, E. F., Schnegelberger, R. D., Falkner, K. C., Prough, R. A., Kirpich, I. A. and Cave, M. C. (2019). Mechanisms of Environmental Contributions to Fatty Liver Disease. Current environmental health reports 6(3), 80–94. PMC6698418

Warner, M., Mocarelli, P., Brambilla, P., Wesselink, A., Samuels, S., Signorini, S. and Eskenazi, B. (2013). Diabetes, metabolic syndrome, and obesity in relation to serum dioxin concentrations: the Seveso women’s health study. Environmental health perspectives 121(8), 906–11. PMC3734493

Wei, Y., Wang, Y. G., Jia, Y., Li, L., Yoon, J., Zhang, S., Wang, Z., Zhang, Y., Zhu, M., Sharma, T., Lin, Y. H., Hsieh, M. H., Albrecht, J. H., Le, P. T., Rosen, C. J., Wang, T. and Zhu, H. (2021). Liver homeostasis is maintained by midlobular zone 2 hepatocytes. Science 371(6532). PMC8496420

Wiegman, C. H., Bandsma, R. H., Ouwens, M., van der Sluijs, F. H., Havinga, R., Boer, T., Reijngoud, D. J., Romijn, J. A. and Kuipers, F. (2003). Hepatic VLDL production in ob/ob mice is not stimulated by massive de novo lipogenesis but is less sensitive to the suppressive effects of insulin. Diabetes 52(5), 1081–9.

Xiong, X., Kuang, H., Ansari, S., Liu, T., Gong, J., Wang, S., Zhao, X. Y., Ji, Y., Li, C., Guo, L., Zhou, L., Chen, Z., Leon-Mimila, P., Chung, M. T., Kurabayashi, K., Opp, J., Campos-Perez, F., Villamil-Ramirez, H., Canizales-Quinteros, S., Lyons, R., Lumeng, C. N., Zhou, B., Qi, L., Huertas-Vazquez, A., Lusis, A. J., Xu, X. Z. S., Li, S., Yu, Y., Li, J. Z. and Lin, J. D. (2019). Landscape of Intercellular Crosstalk in Healthy and NASH Liver Revealed by Single-Cell Secretome Gene Analysis. Mol Cell 75(3), 644–660 e5. PMC7262680

Yang, Y., Filipovic, D. and Bhattacharya, S. (2022). A Negative Feedback Loop and Transcription Factor Cooperation Regulate Zonal Gene Induction by 2, 3, 7, 8-Tetrachlorodibenzo-p-Dioxin in the Mouse Liver. Hepatol Commun 6(4), 750–764. PMC8948569

Zhang, M., Haughey, M., Wang, N. Y., Blease, K., Kapoun, A. M., Couto, S., Belka, I., Hoey, T., Groza, M., Hartke, J., Bennett, B., Cain, J., Gurney, A., Benish, B., Castiglioni, P., Drew, C., Lachowicz, J., Carayannopoulos, L., Nathan, S. D., Distler, J., Brenner, D. A., Hariharan, K., Cho, H. and Xie, W. (2020). Targeting the Wnt signaling pathway through R-spondin 3 identifies an anti-fibrosis treatment strategy for multiple organs. PloS one 15(3), e0229445. PMC7065809

Zhang, P., Kuang, H., He, Y., Idiga, S. O., Li, S., Chen, Z., Yang, Z., Cai, X., Zhang, K., Potthoff, M. J., Xu, Y. and Lin, J. D. (2018). NRG1-Fc improves metabolic health via dual hepatic and central action. JCI insight 3(5). PMC5922292

