## Supplementary figures and images for "Dose-dependent disruption of hepatic zonation by 2,3,7,8-tetrachlorodibenzo-*p*-dioxin in mice: integration of single-nuclei RNA sequencing and spatial transcriptomics"

### heatmap_count.pdf

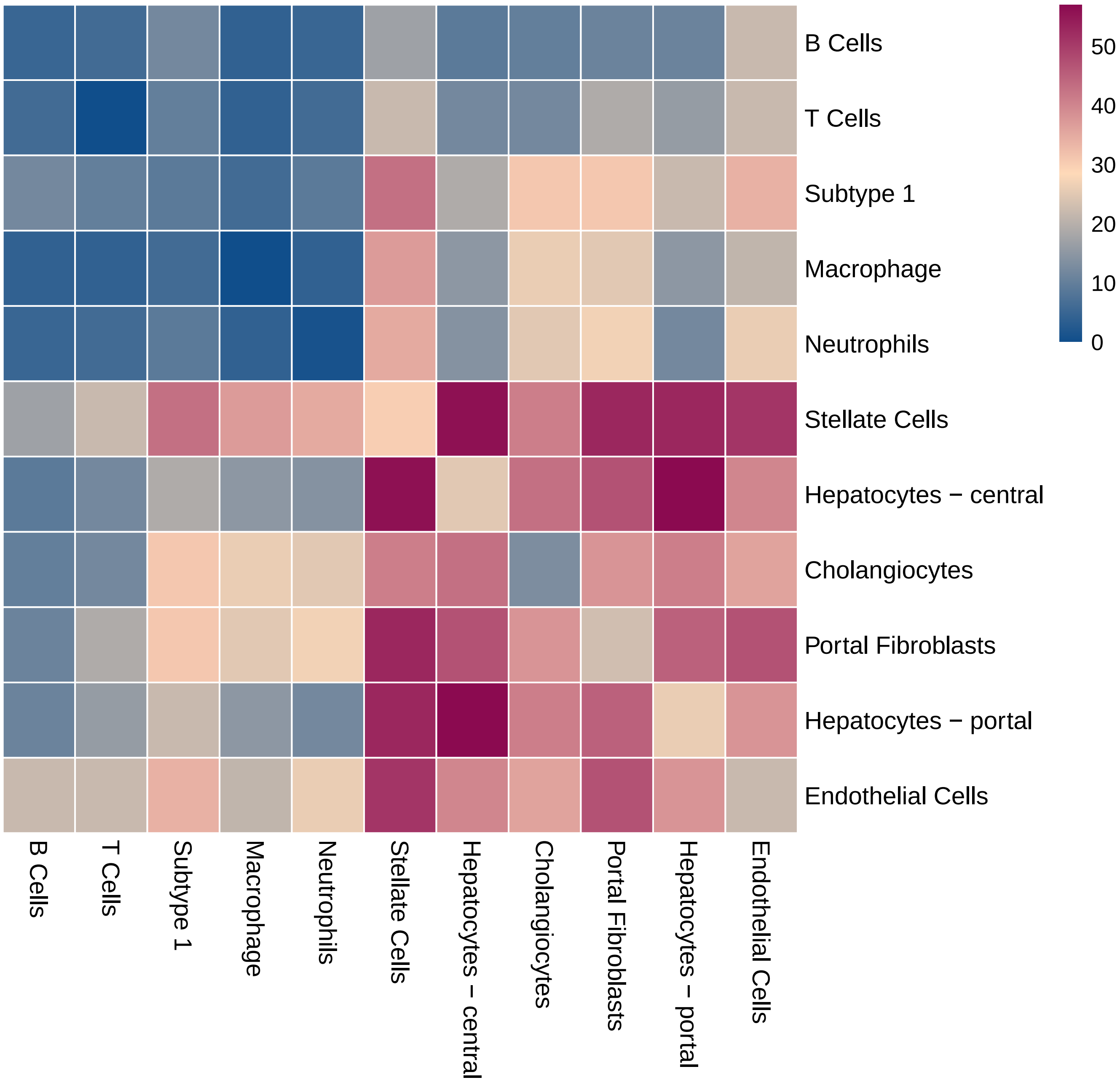

### heatmap_count.pdf

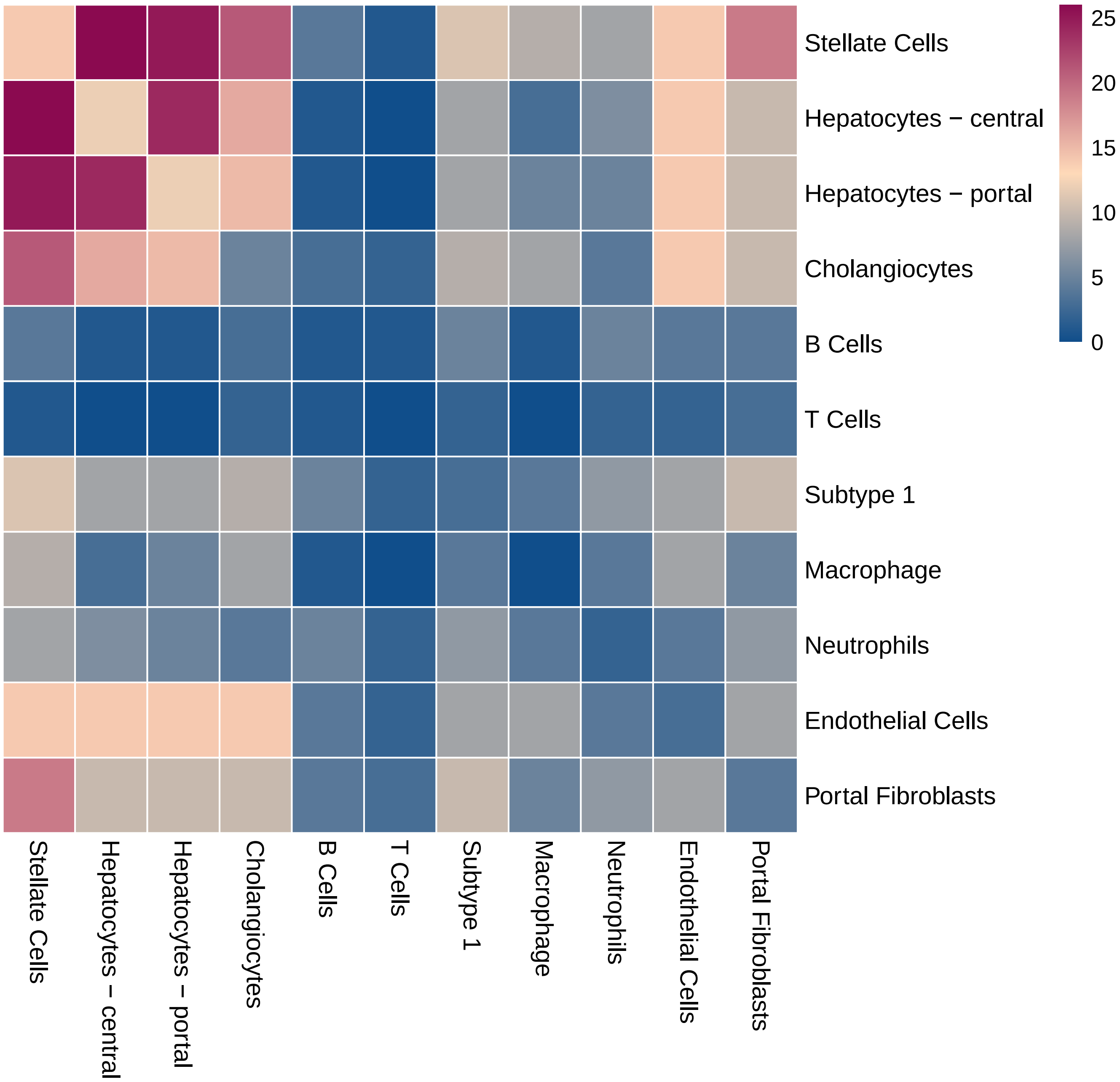

### heatmap_count.pdf

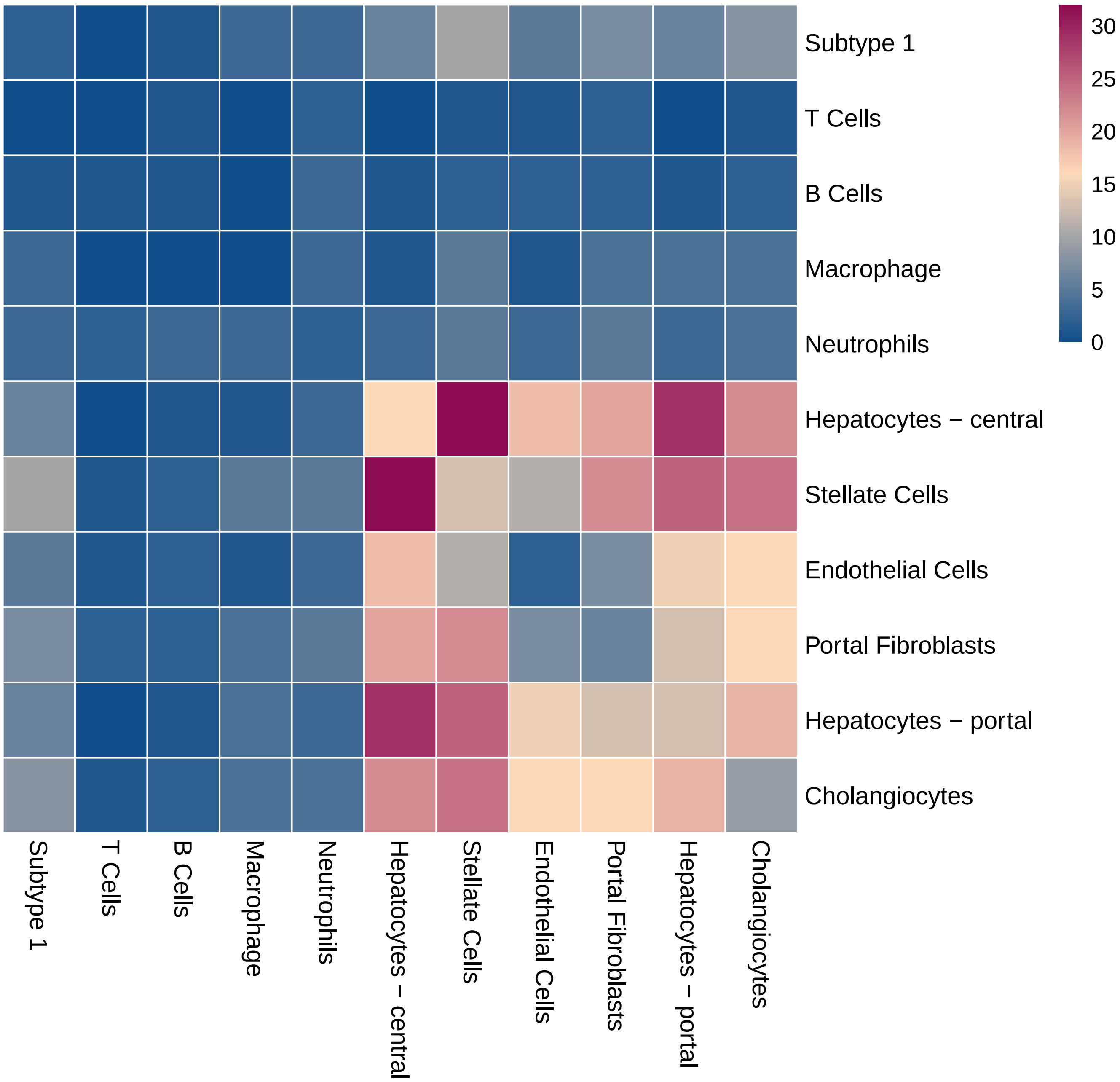

### heatmap_count.pdf

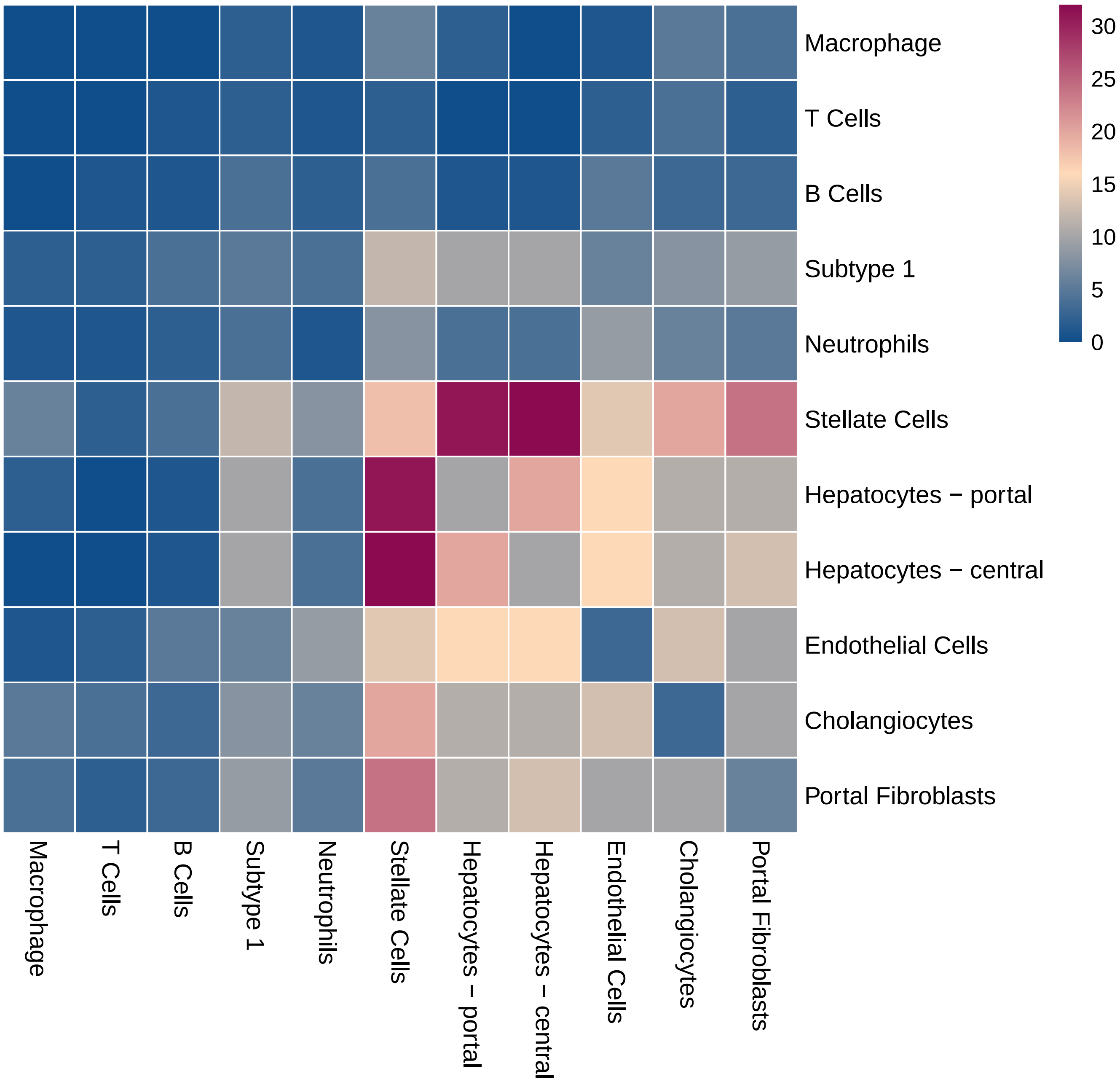

### heatmap_count.pdf

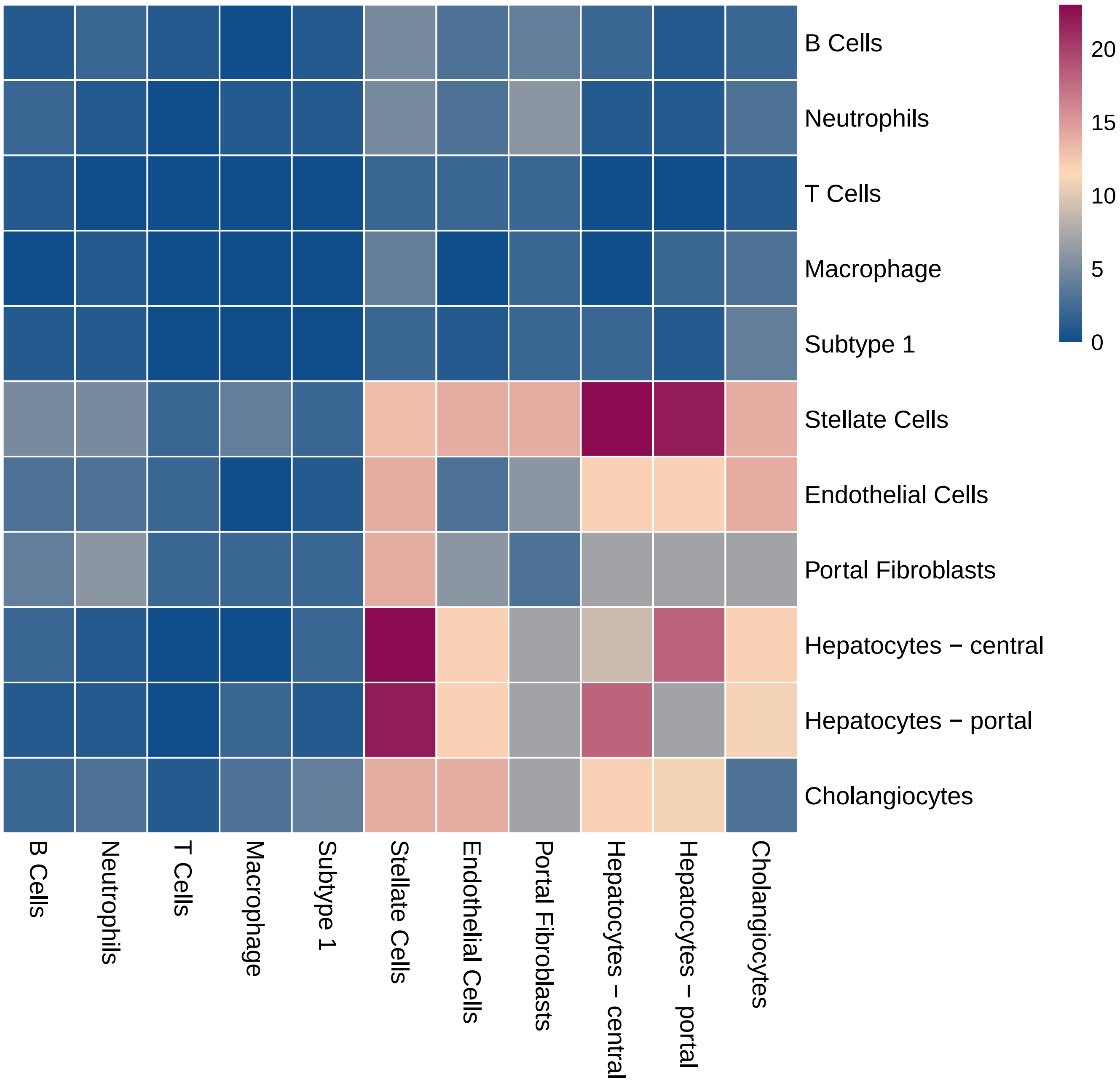

### heatmap_count.pdf

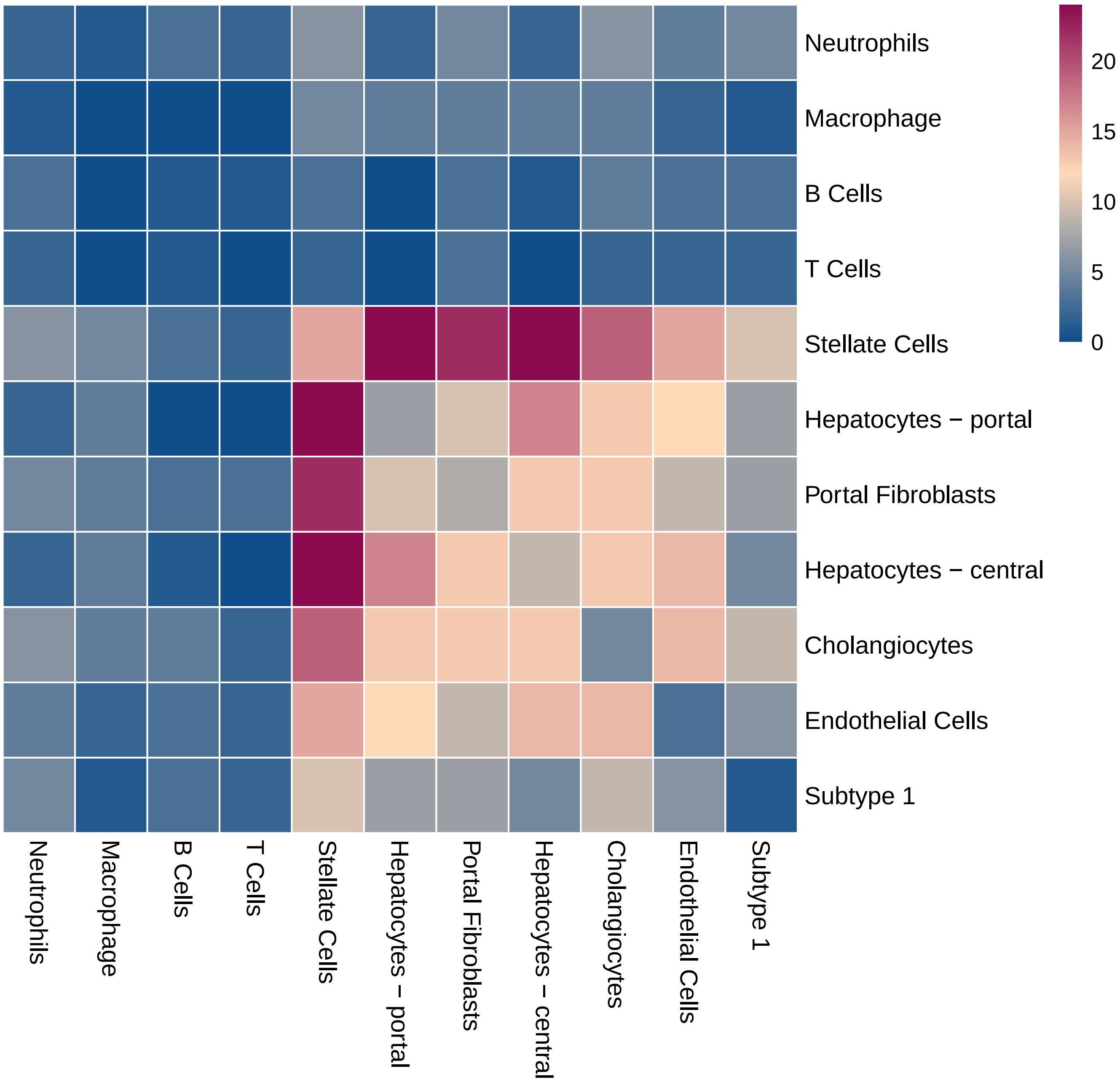

### heatmap_count.pdf

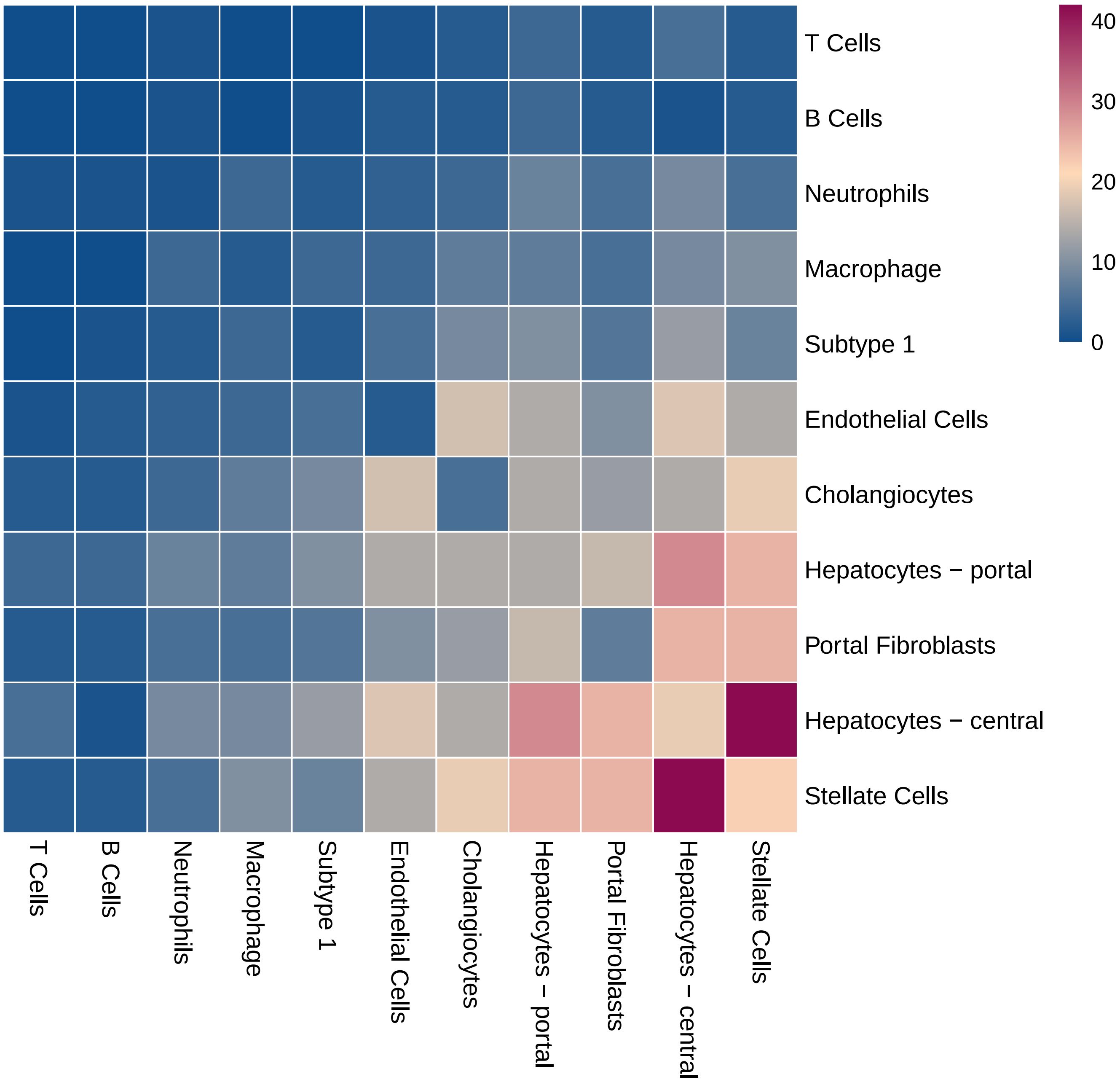

### heatmap_count.pdf

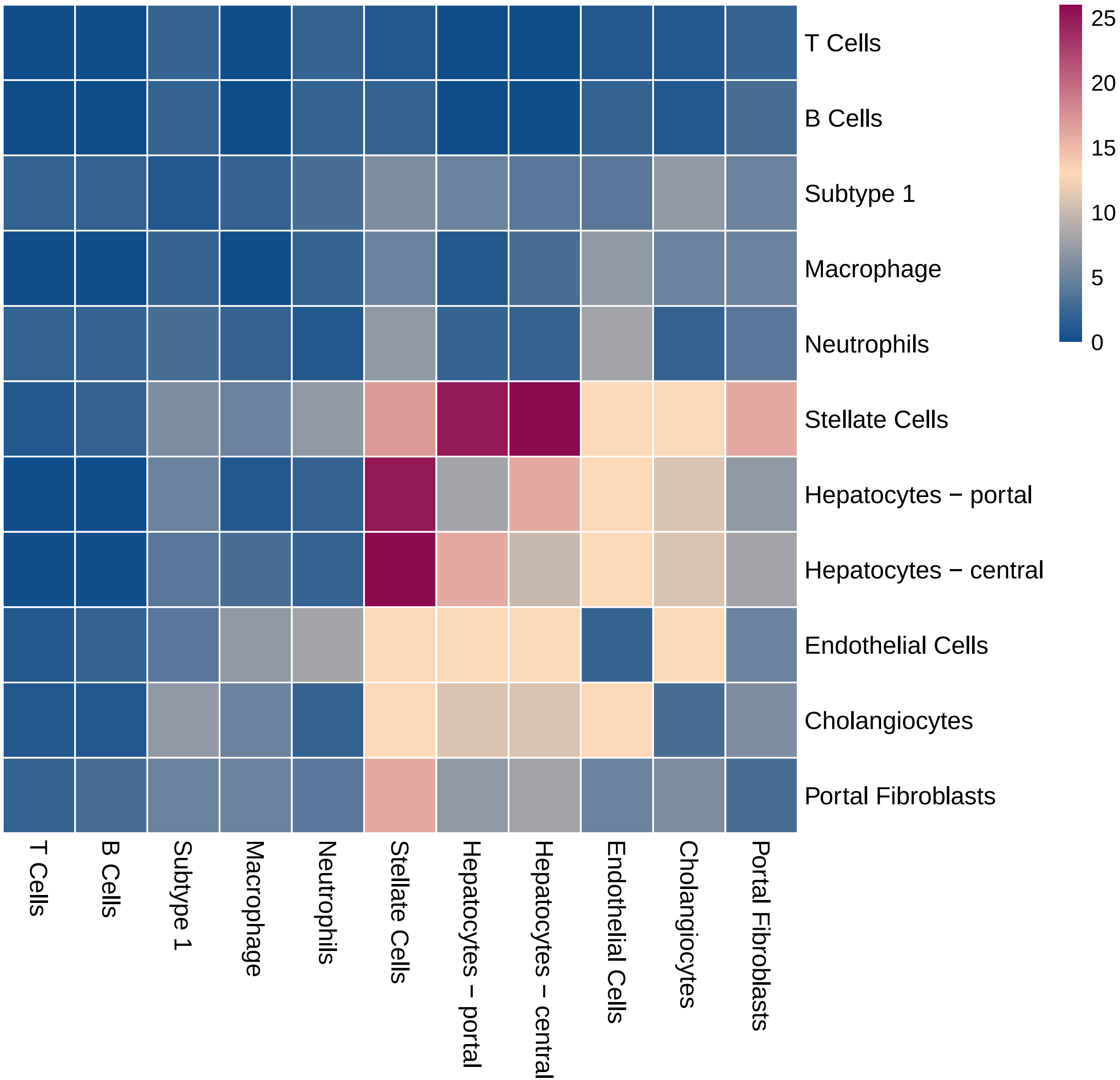

### heatmap_count.pdf

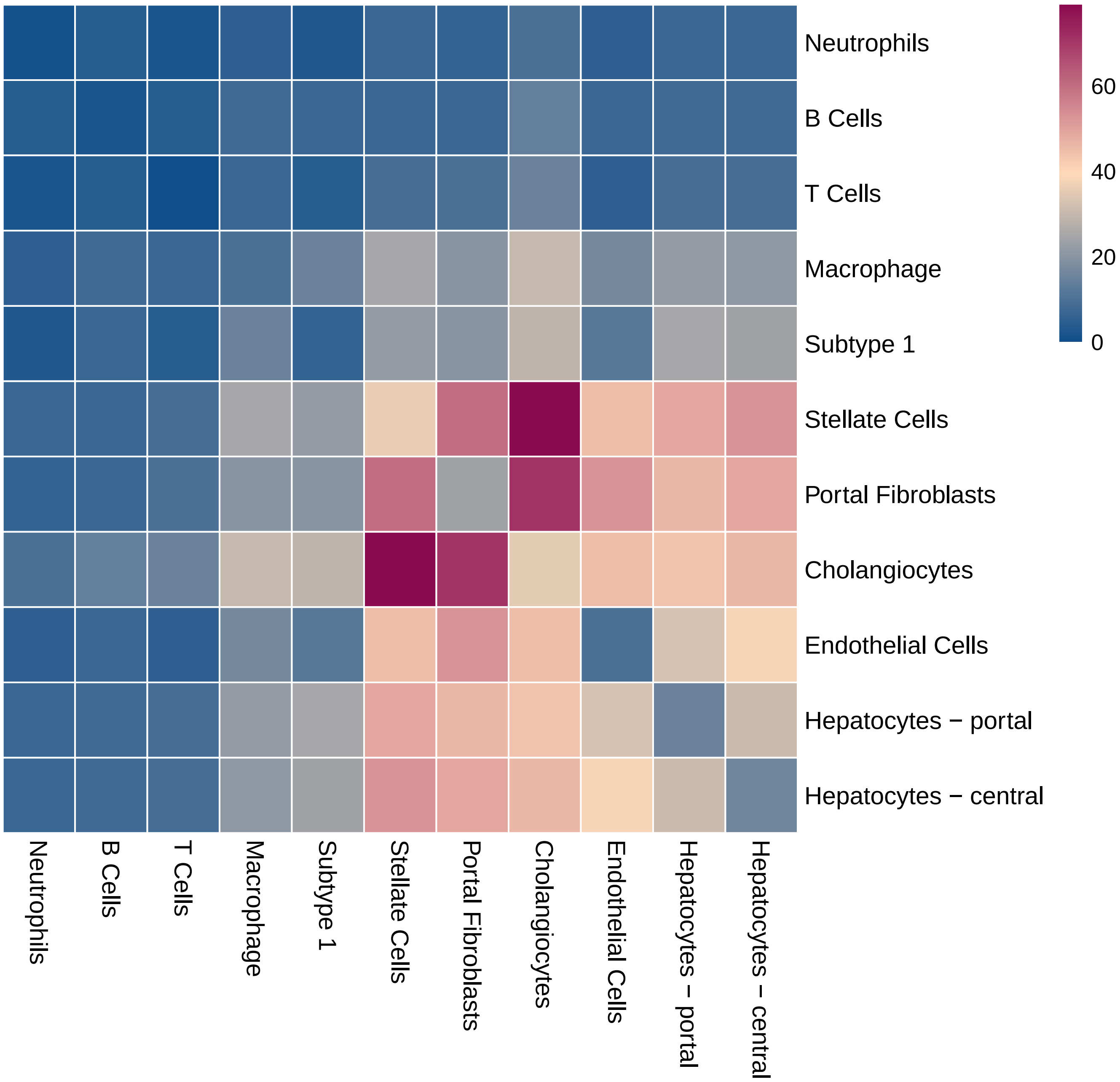

### heatmap_log_count.pdf

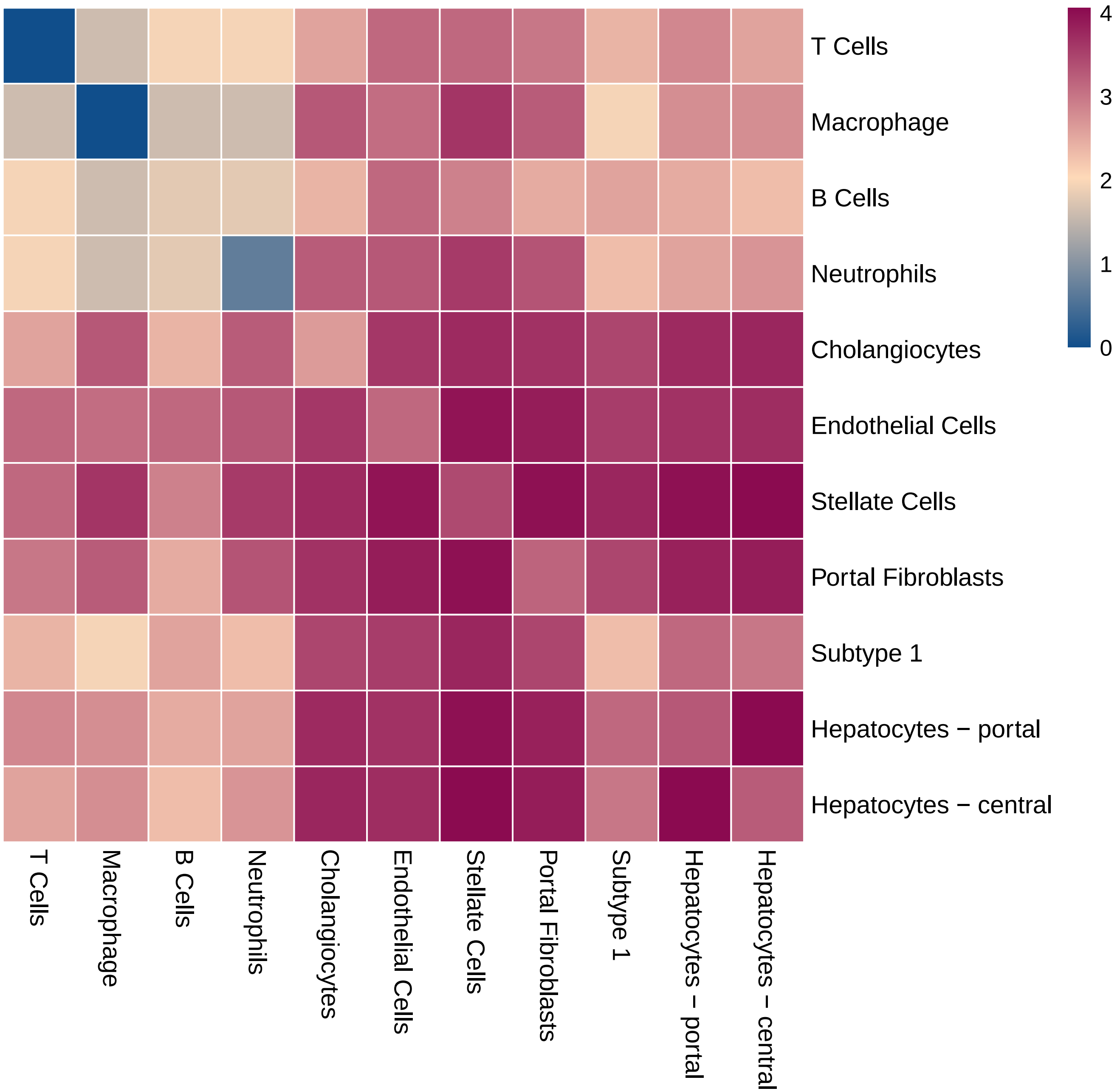

### heatmap_log_count.pdf

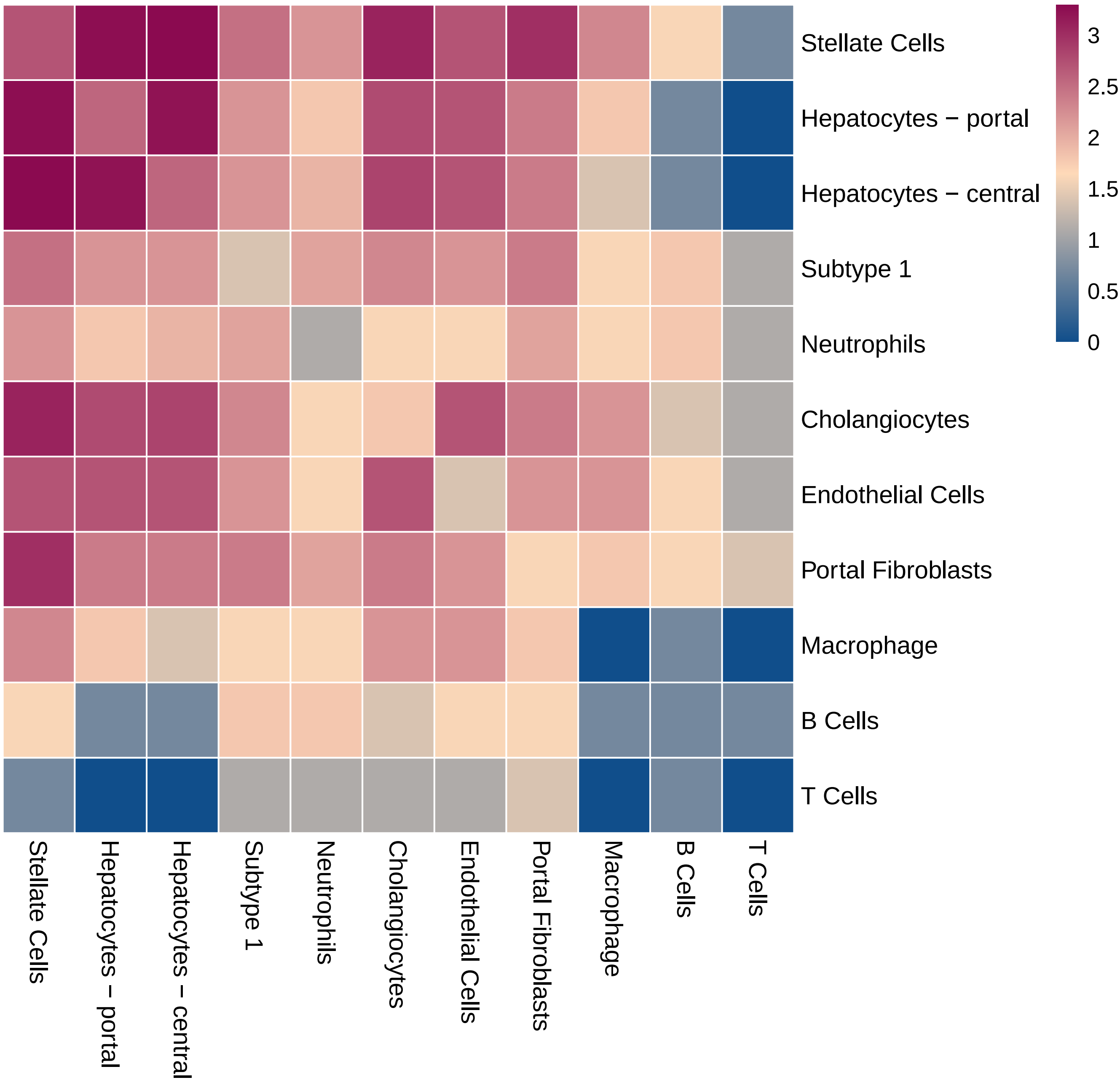

### heatmap_log_count.pdf

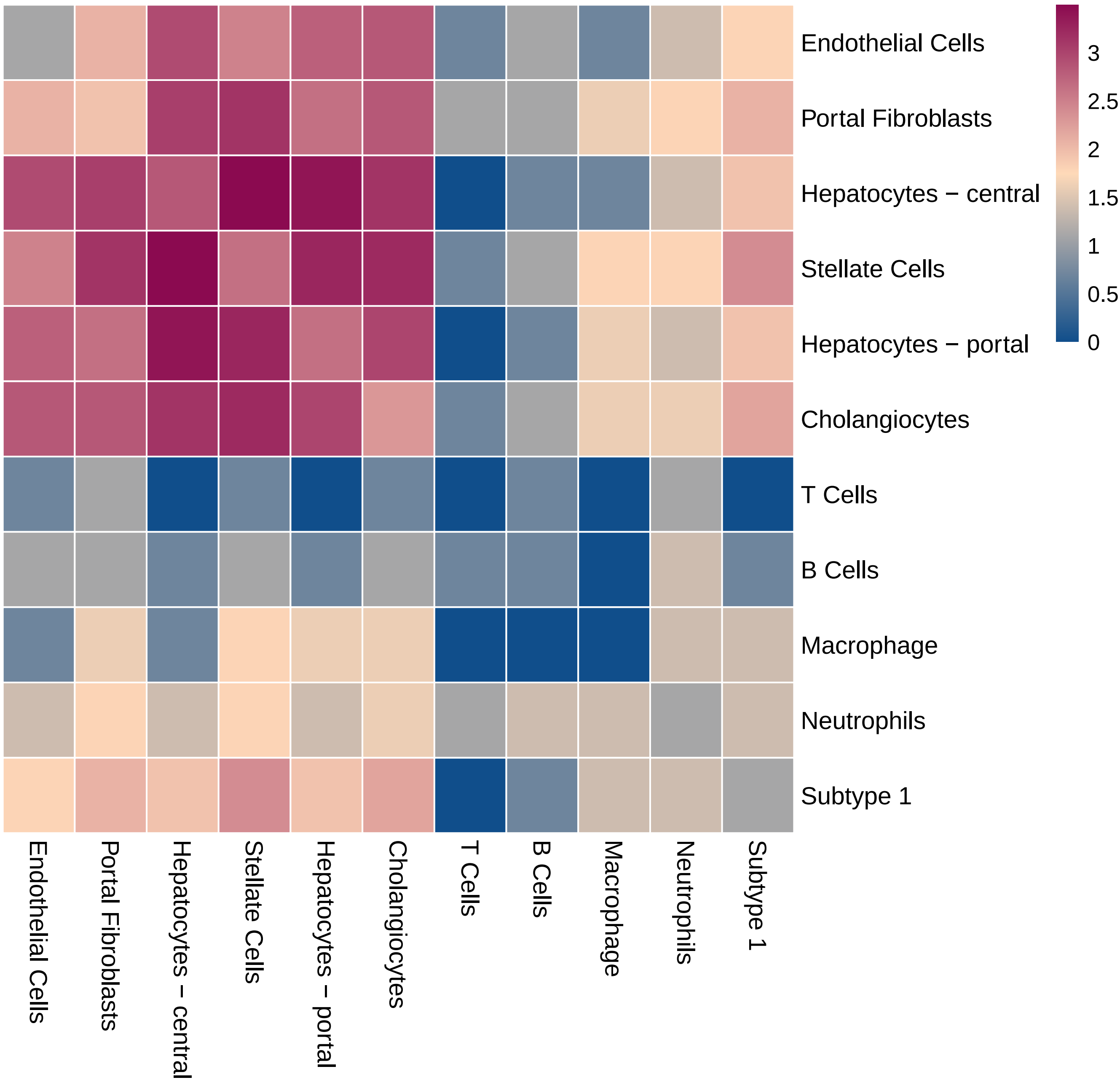

### heatmap_log_count.pdf

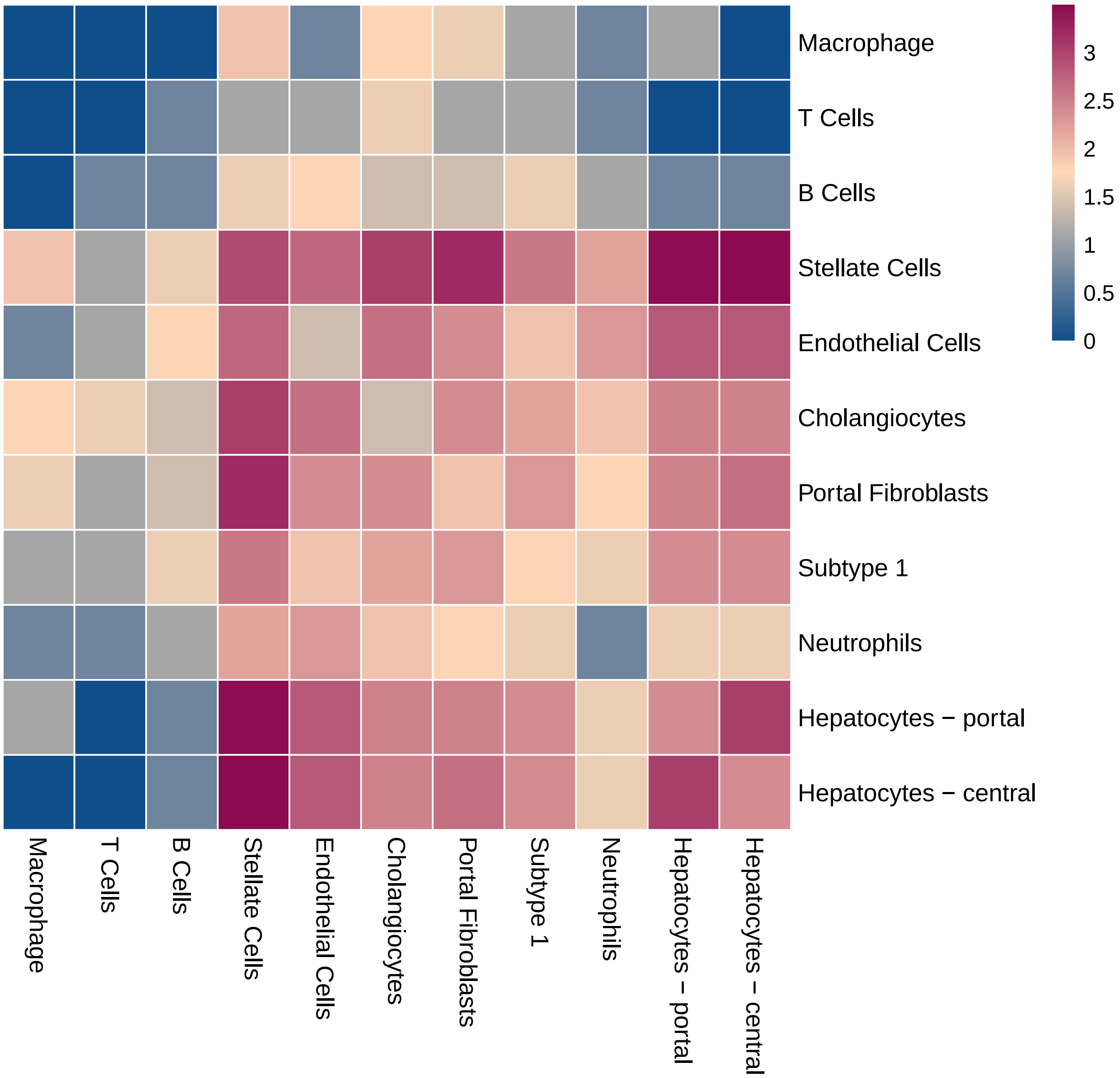

### heatmap_log_count.pdf

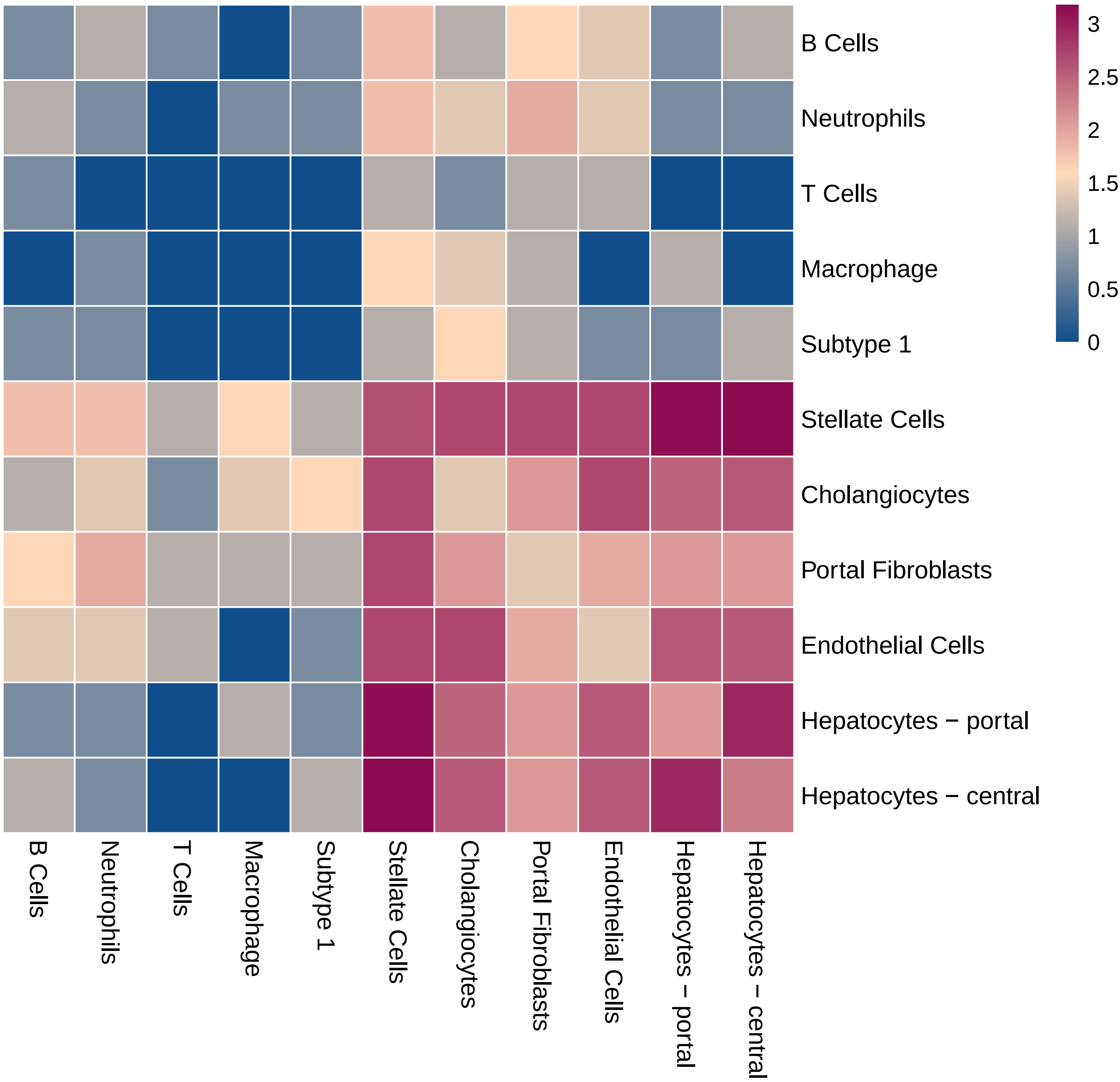

### heatmap_log_count.pdf

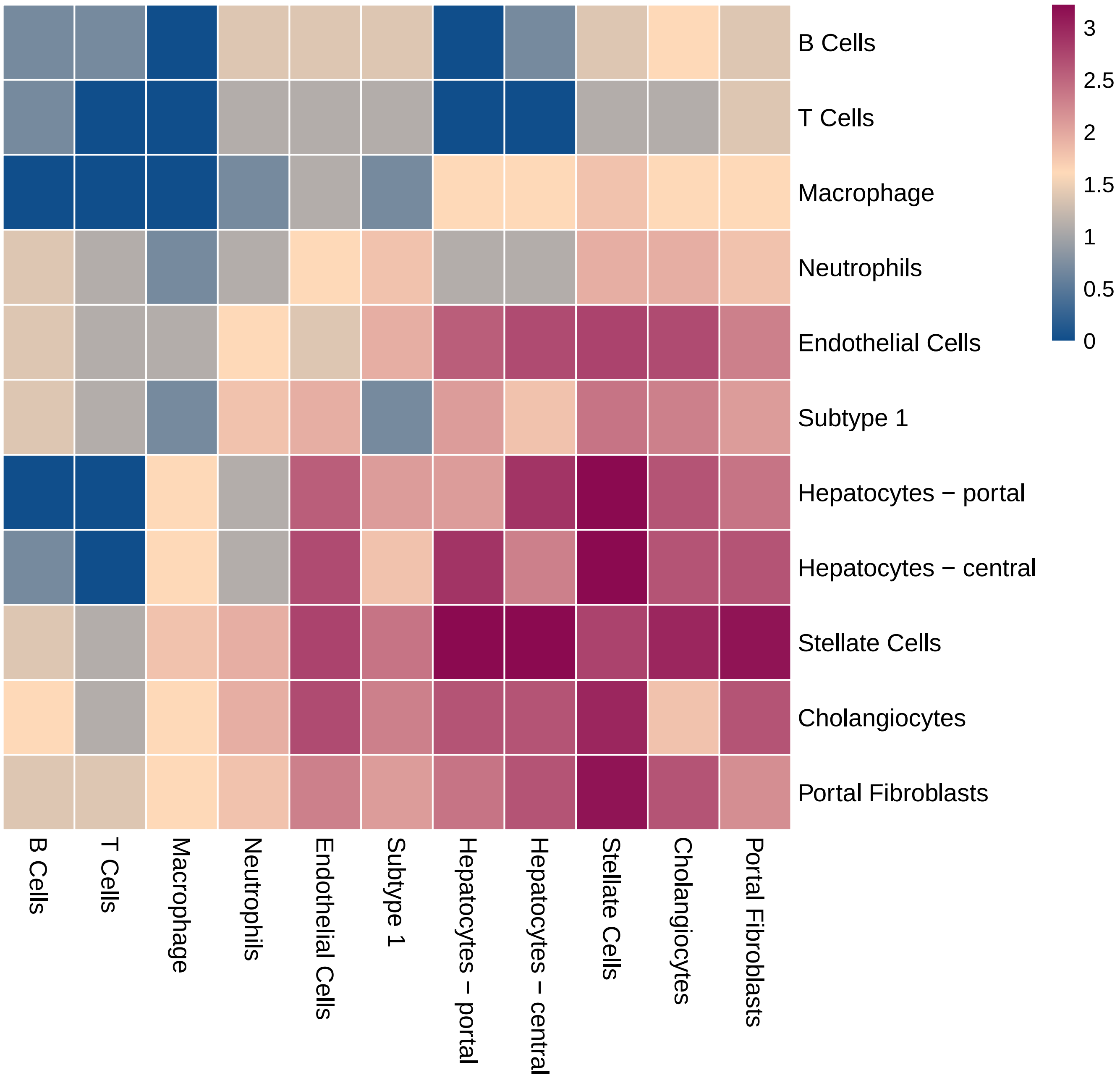

### heatmap_log_count.pdf

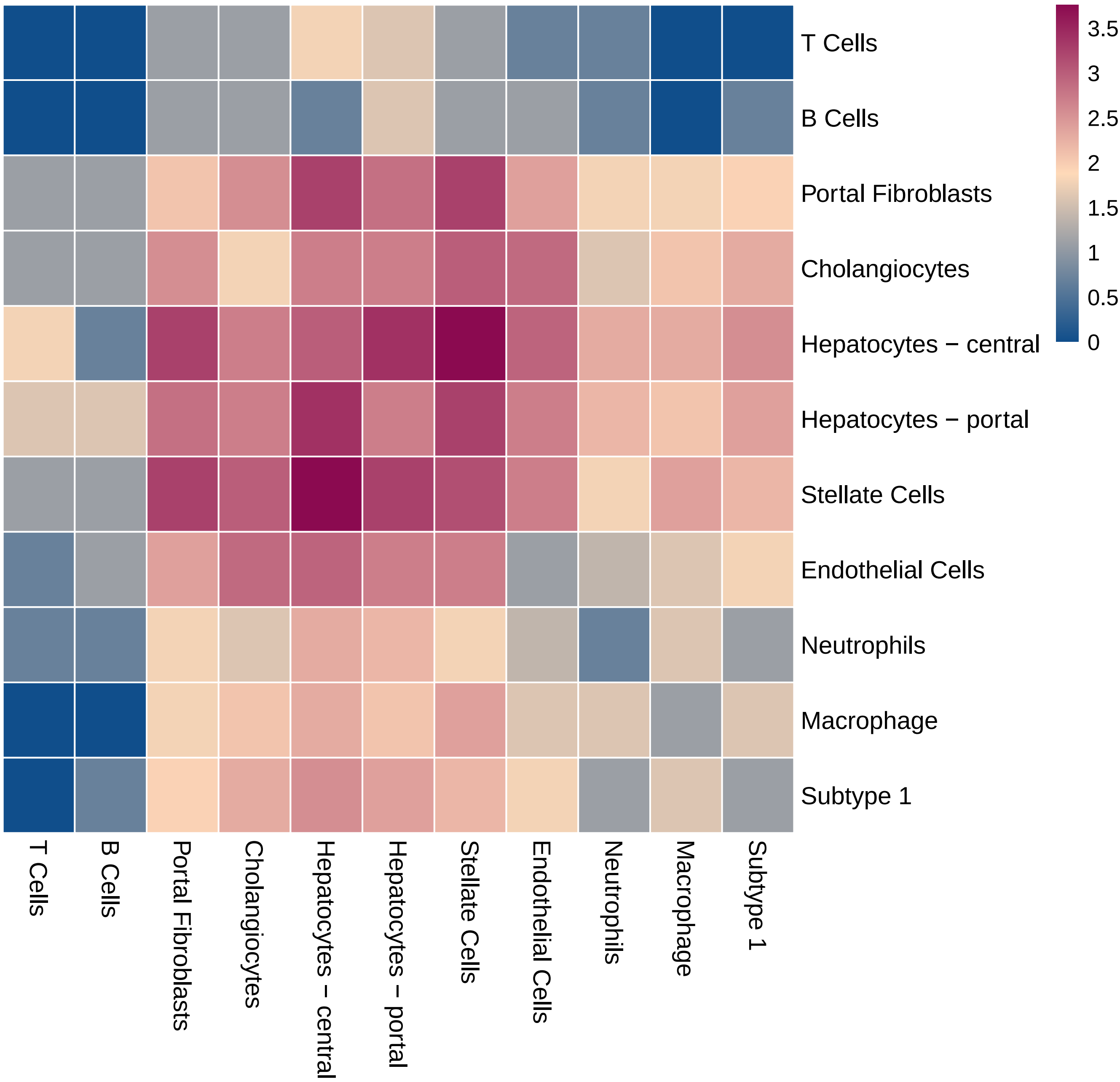

### heatmap_log_count.pdf

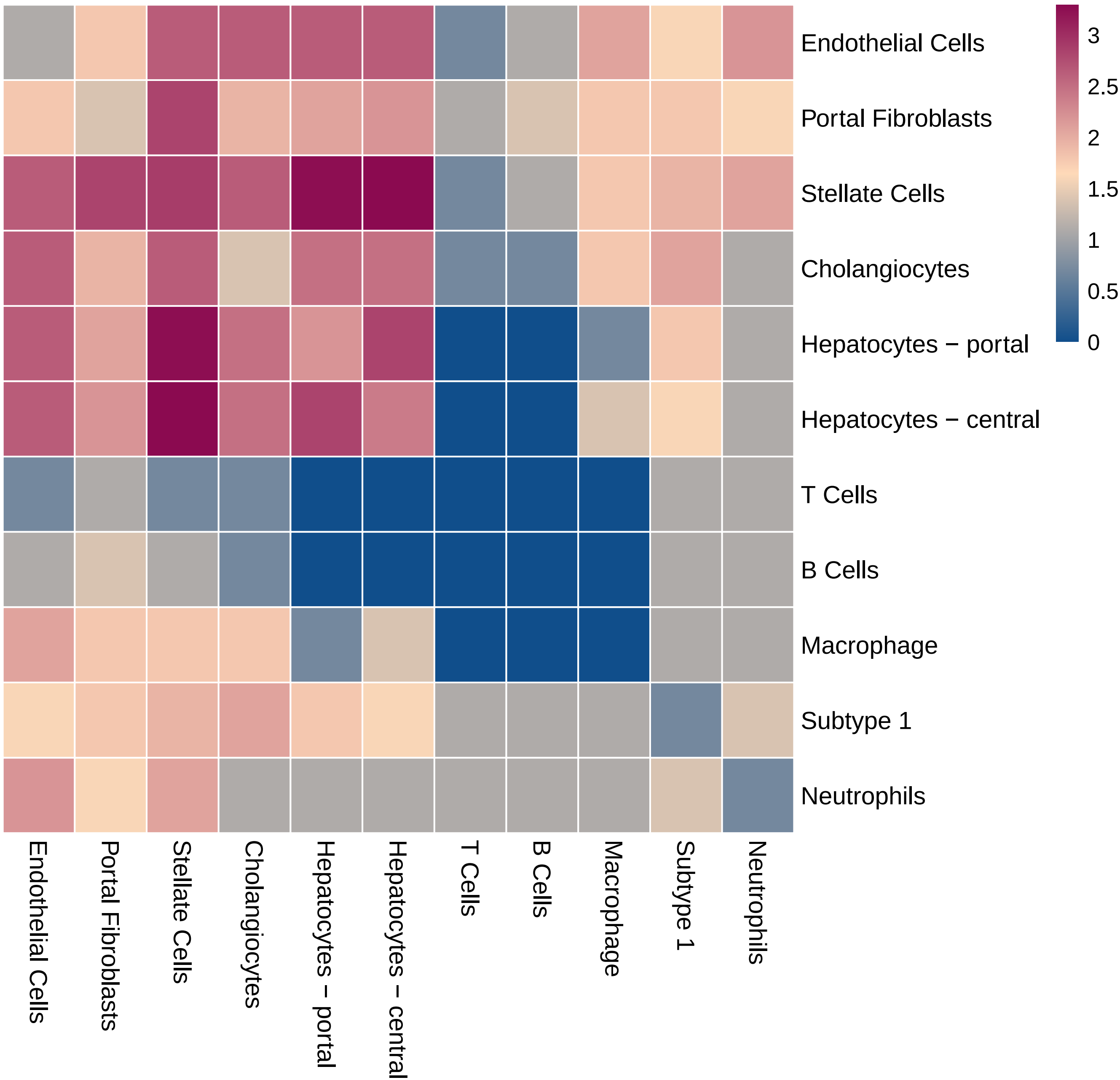

### heatmap_log_count.pdf

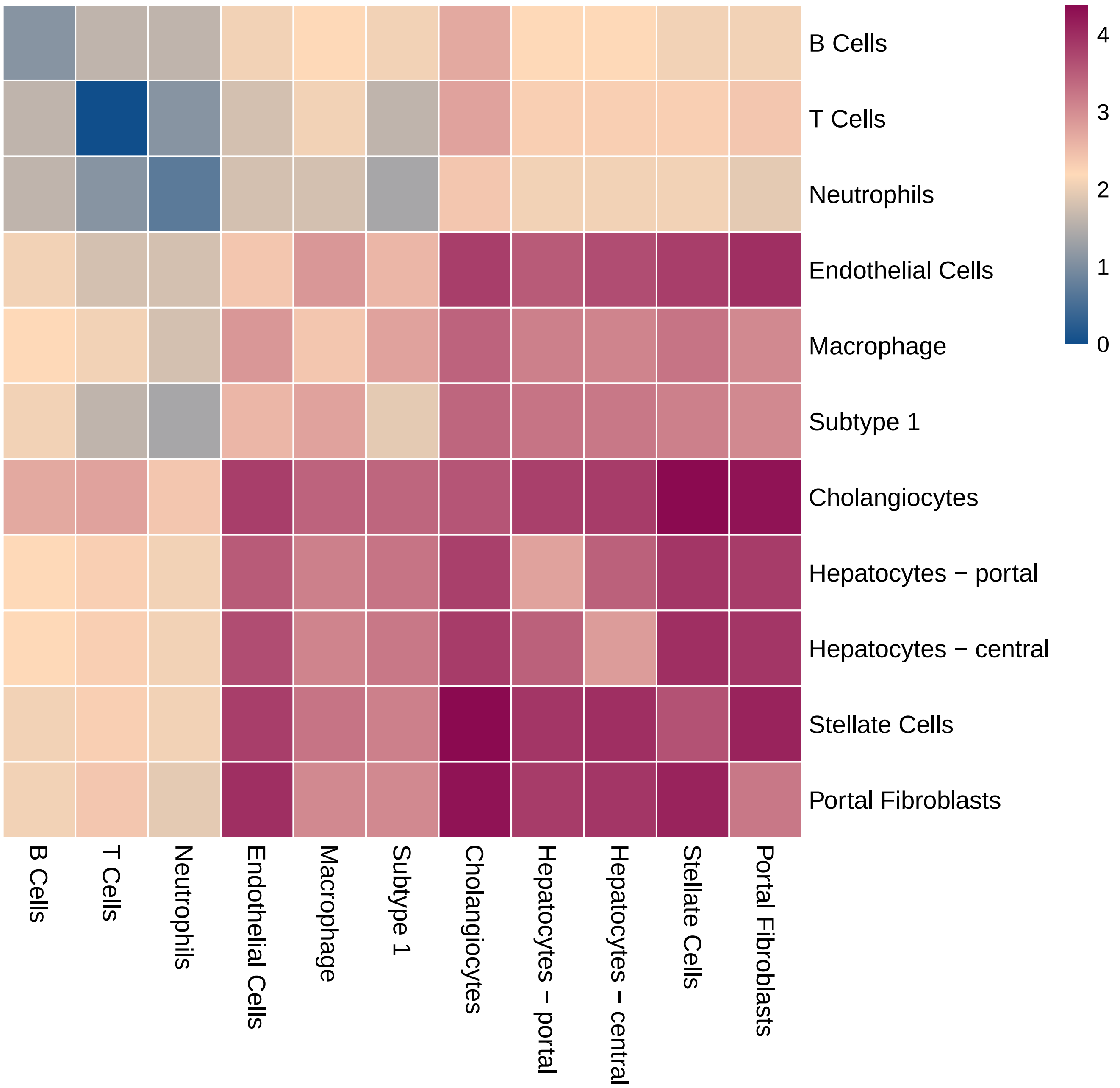
