## Supplementary material for "Dose-dependent disruption of hepatic zonation by 2,3,7,8-tetrachlorodibenzo-*p*-dioxin in mice: integration of single-nuclei RNA sequencing and spatial transcriptomics": Figure S6: FigS6-CellPhoneDB network.nb.html


Code 

- Show All Code
- Hide All Code
- Download Rmd

### Figure S6


**Figure S6.** Network visualization of ligand-receptor pair interactions determined by using CellPhoneDB. Interactions were identified for each individual dose group independently including genes expressed in at least 25% of nuclei. Mean values for ligand-receptor pairs (mean of the means for each member of the interaction) was calculated at each dose and only interactions exhibiting a ≥ 2-fold increase relative to control and p-value ≤ 0.05 were kept.
